# Spatially organized inflammatory myeloid-CD8^+^ T cell aggregates linked to Merkel-cell Polyomavirus driven Reorganization of the Tumor Microenvironment

**DOI:** 10.1101/2025.06.06.657162

**Authors:** Maximilian Haist, Magdalena Matusiak, Yuqi Tan, Stefanie Zimmer, Henner Stege, Tim Noah Kempchen, Silke Mitschke, Pauline Chu, Beate Weidenthaler-Barth, Graham L. Barlow, Friederike Rogall, Antonio Delgado Gonzalez, Marc-A. Baertsch, Silvia Albertini, Marisa Gariglio, Cinzia Borgogna, Yury Goltsev, Stephan Grabbe, John W. Hickey, Garry P. Nolan

## Abstract

Merkel cell carcinoma (MCC) is an aggressive skin cancer with high propensity for metastasis, caused by Merkel-cell-polyomavirus (MCPyV), or chronic UV-light-exposure. How MCPyV spatially modulates immune responses within the tumor microenvironment and how such are linked to patient outcomes remains unknown. We interrogated the cellular and transcriptional landscapes of 60 MCC-patients using a combination of multiplex proteomics, *in-situ* RNA-hybridization, and spatially oriented transcriptomics. We identified a spatial co-enrichment of activated CD8^+^ T-cells and CXCL9^+^PD-L1^+^ macrophages at the invasive front of virus-positive MCC. This spatial immune response pattern was conserved in another virus-positive tumor, HPV^+^ head-and-neck cancer. Importantly, we show that virus-negativity correlated with high risk of metastasis through low CD8^+^ T-cell infiltration and the enrichment of cancer-associated-fibroblasts at the tumor boundary. By contrast, responses to immune-checkpoint blockade (ICB) were independent of viral-status but correlated with the presence of a B-cell-enriched spatial contexts. Our work is the first to reveal distinct immune-response patterns between virus-positive and virus-negative MCC and their impact on metastasis and ICB-response.

## 1. Introduction

Merkel cell carcinoma (MCC) is an aggressive neuroendocrine skin cancer characterized by its propensity to metastasize following primary diagnosis^1^. Specifically, about one-third of MCC patients with initial local disease will eventually develop distant metastasis that results in a high mortality rate of 33-46%. This highlights the urgent need for biomarkers that would identify patients at high risk of metastasis^2^.

MCC is classically divided into two distinct molecular subclasses: virus-positive (VP) and virus-negative (VN), which are morphologically indistinguishable by conventional histology. VP-MCC results from the integration of the Merkel cell polyomavirus (MCPyV) into the host genome and expression of viral oncoproteins that cause the inactivation of tumor suppressor genes^3^. By contrast, VN-MCC is caused by UV-induced DNA damage and is characterized by a high mutational burden*^4^*. Several reports have documented more favorable survival outcomes for VP-MCC patients^4-6^. However, whether the difference in survival between VP- and VN-MCC is due to comorbidities often found in the predominantly older VN-MC population or inherent biological differences associated with MCPyV infection remains unknown.

Our current understanding of anti-tumor immune responses in MCC is largely derived from flow cytometry, T-cell receptor sequencing, low-dimensional immunohistochemistry (IHC), and bulk sequencing data that lack either spatial context or single-cell resolution^4,7,8^. Hence, critical aspects of the *in-situ* tumor ecosystem can only be inferred. To capture the complexity of tumor-immune interactions within the native TME directly and on the single-cell level, we here apply the first comprehensive multiplex imaging study of the MCC-TME utilizing co-detection by indexing combined with in-situ RNA hybridization (RNAscope) imaging and targeted RNA sequencing through laser-capture microdissection (LCM). This allowed us to address two questions: (1) How MCPyV modulates anti-tumor immune responses within the TME and (2) how these tissue signatures are linked to patient outcomes.

Our results demonstrate that the VP-TME is characterized by a spatial co-enrichment of CD8^+^ T-cells with CXCL9^+^PD-L1^+^HLA-DR^+^ macrophages and dendritic cells (DCs) at the tumor invasive front. We show that virus-positivity and high CD8^+^ T cell infiltration at the invasive front predict prolonged metastasis-free survival in MCC patients. By contrast the enrichment of cancer-associated fibroblasts (CAFs) at the tumor boundary associated with higher risk for distant metastasis. Importantly, we found that responses to immune-checkpoint-blockade (ICB), being the current standard-of-care in advanced MCC patients, associated with a B-cell enriched spatial context that was independent of viral-status.

In summary, we have created the first single-cell spatial map of MCPyV-mediated immunomodulation of the TME, that informs novel strategies for patient stratification at early disease stages and provide a rationale for biomarker-informed adjuvant ICB therapy of MCC patients at high-risk of metastasis.

## 2. Results

### MCPyV-positive MCC is associated with favorable survival outcomes

We applied a multi-modal experimental framework including CODEX multiplex imaging, RNAscope, and LCM-guided bulk sequencing (LCMseq) (**Fig.1A**) in conjunction with an analytical framework to dissect spatial imaging data (**Fig.1B**). This strategy allowed us to uncover MCPyV-associated anti-tumor immune responses within the native tissue context and delineate how those are linked to patient outcomes (**Fig.1C**). From a database of 242 patients with MCC, we selected a cohort of 60 MCC patients with detailed clinical documentation, long-term follow up and known MCPyV-status (29 VN and 31 VP patients) (**Supplemental Fig.1A-B, Fig.1D**). Importantly, VP and VN groups were matched for main clinical parameters such as demographics, primary tumor characteristics, tumor stage, treatments, investigated tissues and follow-up (**Supplemental Table1, Supplemental Fig.1C-D**). In line with previous reports, we observed a trend towards a longer overall survival (OS) in VP-MCC patients as compared to VN-MCC patients (**Fig.1E**, **Supplemental Tables1-2**)^4,6^. In addition, we found that VP-MCC patients with local stage I/II disease had a significantly longer distant-metastasis-free survival (DMFS) (median DMFS: not reached vs 7 months, *p*=0.0034), that was confirmed in multivariate testing (**Supplemental Fig.1E-F**, **Supplemental Tables3-4**).

**Fig. 1:**
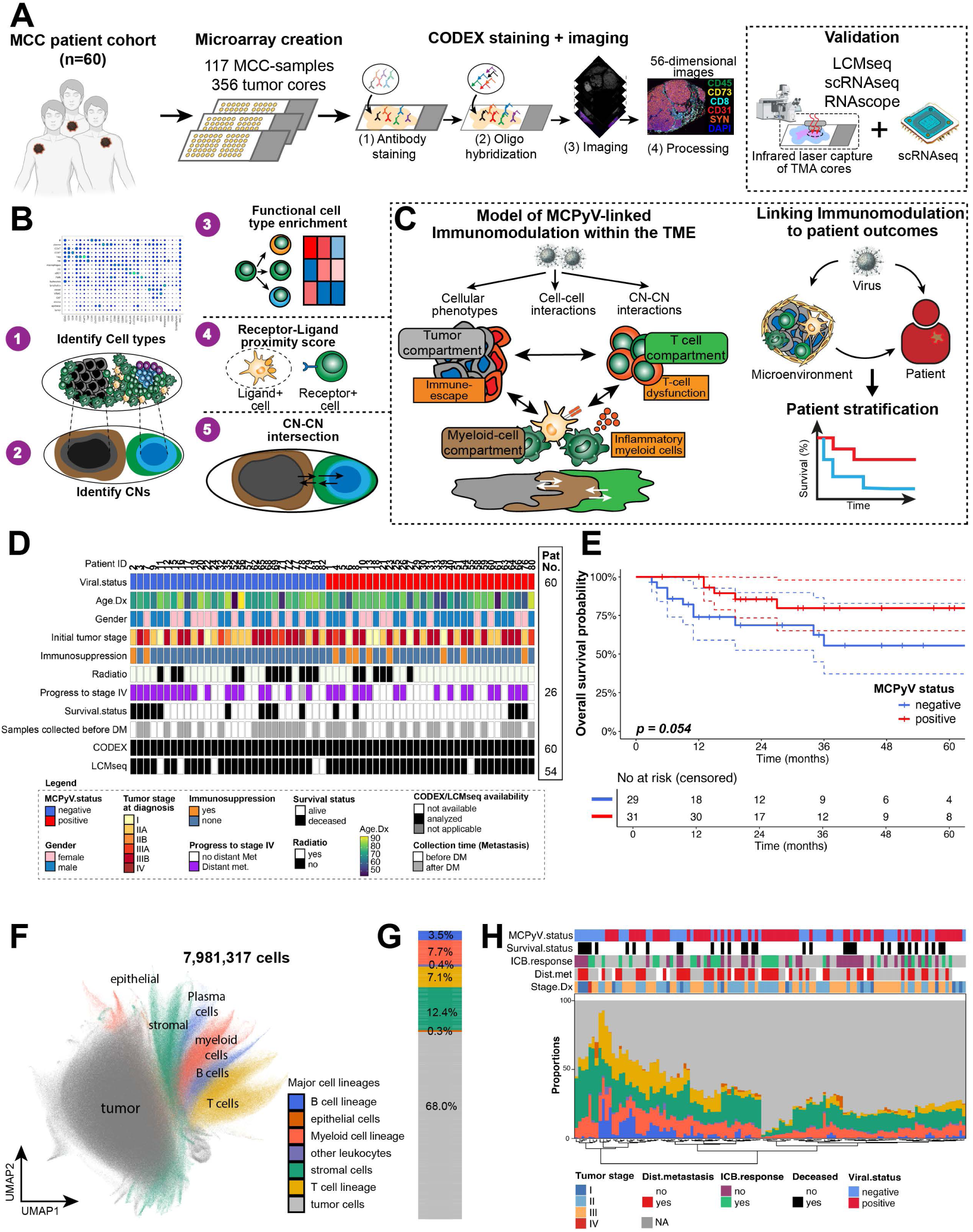
Study design, workflow and profiling of the MCC-TME. **(A)** Conceptual framework and experimental setup. **(B)** Analytical setup. **(C)** Left: Illustration of key characteristics of the VP MCC-TME. Right: Cartoon depicting the proposed link between viral infection, immunomodulation within the TME and patient outcomes. **(D)** Summary of patient and sample characteristics within the investigated patient cohort stratified by viral status. **(E)** Kaplan Meier plot depicting prolonged overall survival for patients with MCPyV-positive MCC as compared to virus-negative disease (p-value determined by log-rank test). **(F)** Major cell types identified within the CODEX data set represented as UMAP projection. **(G)** Proportions of major cell types in the overall data set. Color-code from Fig.1F applies. **(H)** Mean proportions of major cell types per investigated tumor sample with corresponding clinical annotations for location of tissue biopsies, viral status, immunotherapy response, tumor stage at diagnosis and the event of distant metastasis. Each column depicts a single MCC tumor sample. Samples are ordered by Euclidian clustering. Color-codes for major cell types shown in Fig.1F.

### *In-Situ* Profiling of the MCC-TME using CODEX multiplex imaging

We reasoned that the identification of virus-specific immune signatures and their spatial organization might help explain the differential survival outcomes found between VP and VN-MCCs. To identify such tissue signatures, we, determined MCPyV-status for all MCC samples and constructed tissue microarrays (TMA) of the MCC-TME that incorporated representative tumor regions from both the tumor-invasive front and the tumor core for each patient (**Supplemental Fig.2A-F**). We developed and validated a 56-marker CODEX antibody panel that includes antibodies for cell phenotyping and functional assessment such as markers for macrophages (CD68, CD163, CD206), B cells, T cell-specific markers, tumor markers (CK20, Synaptophysin, CD56, Chromogranin A), markers of T cell dysfunction (EOMES, PD1, LAG3), and markers of MCPyV-infection (MCPyV LT antigen) (**Supplemental Fig.3A-C**, **Data Fig.S1-4**). We assessed binding specificity and optimized signal-to-noise ratios of the antibodies in a multi-Tumor TMA before optimizing our panel for use in MCC (**Supplemental Fig.3C-D**).

**Fig. 2:**
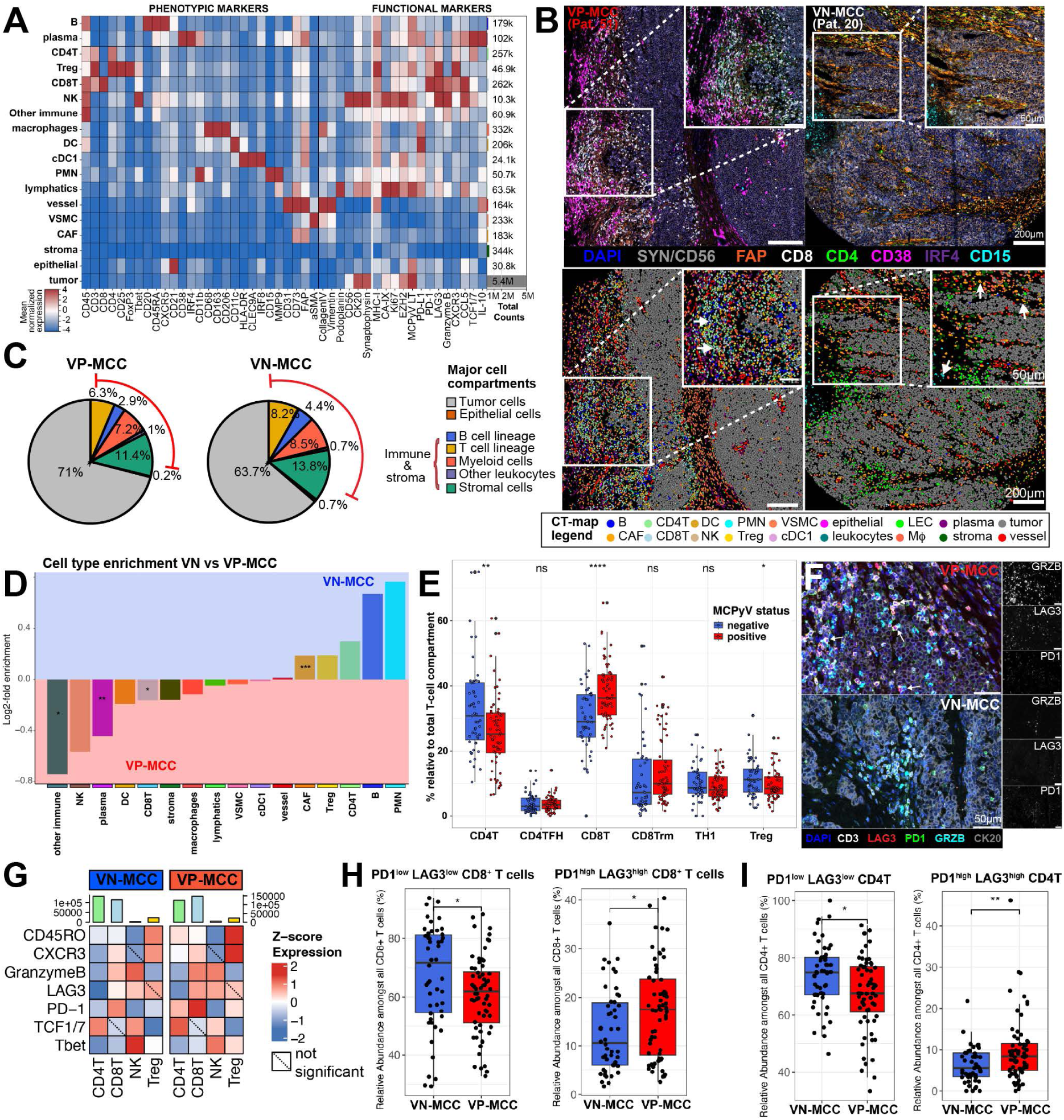
Distinct cellular composition and T-cell phenotypes define the tumor microenvironment of MCPyV-positive and MCPyV-negative MCC. **(A)** Heatmap illustrating log-normalized mean expression of phenotypic (left) and functional markers across the identified cell types of CODEX multiplex imaging with corresponding cell counts. **(B)** Representative images of the VP-MCC and VN-MCC TME (top) with corresponding cell type maps (bottom). Scale bars, 200 µm and 50µm. Black arrows highlight cell types specifically enriched in VP vs VN-MCC. **(C)** Cell type frequencies of the major cell compartments in VP-MCC and VN-MCC normalized to the total cell counts, brackets indicate cell types within immune/stroma compartments. **(D)** Log2-fold enrichment of immune and stroma cell types between VP-MCC and VN-MCC. Tests were adjusted for multiple comparisons using the Bonferroni method with a targeted false discovery rate (FDR) at 0.05 (* *p ≤* 0.05, ** *p ≤* 0.01). **(E)** T cell subtype abundances normalized by the total number of T cells and grouped by viral status. Each point represents the mean cell frequency per MCC-sample (n=117). Statistical significance was determined using t-tests adjusted for multiple comparisons using the Bonferroni method with a targeted FDR at 0.05; * *p ≤* 0.05, ** *p ≤* 0.01, **** *p ≤* 0.0001. **(F)** Representative images of T-cell phenotypes found within VP (top) and VN-MCC with corresponding greyscale images of selected functional markers. Scale bar of 50µm applies to all panels. **(G)** Heatmaps depicting mean z-normalized expression of functional markers across T-cell subtypes between VP and VN-MCC samples. Mean expression values of each marker in a given cell-type was compared between VP and VN-MCC samples using Wilcoxon-rank sum test adjusted for multiple-hypothesis testing (BH). Non-significant interactions are indicated by dotted lines. **(H)** Relative abundance of checkpoint-protein positive and negative CD8^+^ T cell states normalized to the total number of T cells in VP-MCC vs VN-MCC. Each point represents the mean cell frequency per MCC-sample. Significance was determined using Bonferroni-adjusted t-test with FDR <0.05. * *p ≤* 0.05, ** *p ≤* 0.01, **** *p ≤* 0.0001. **(I)** Relative abundance of checkpoint-protein positive and negative CD8^+^ T cell states normalized to the total number of T cells in VP-MCC vs VN-MCC. Each point represents the mean cell frequency per MCC-sample. Significance was determined using Bonferroni-adjusted t-test with FDR <0.05.

**Fig. 3:**
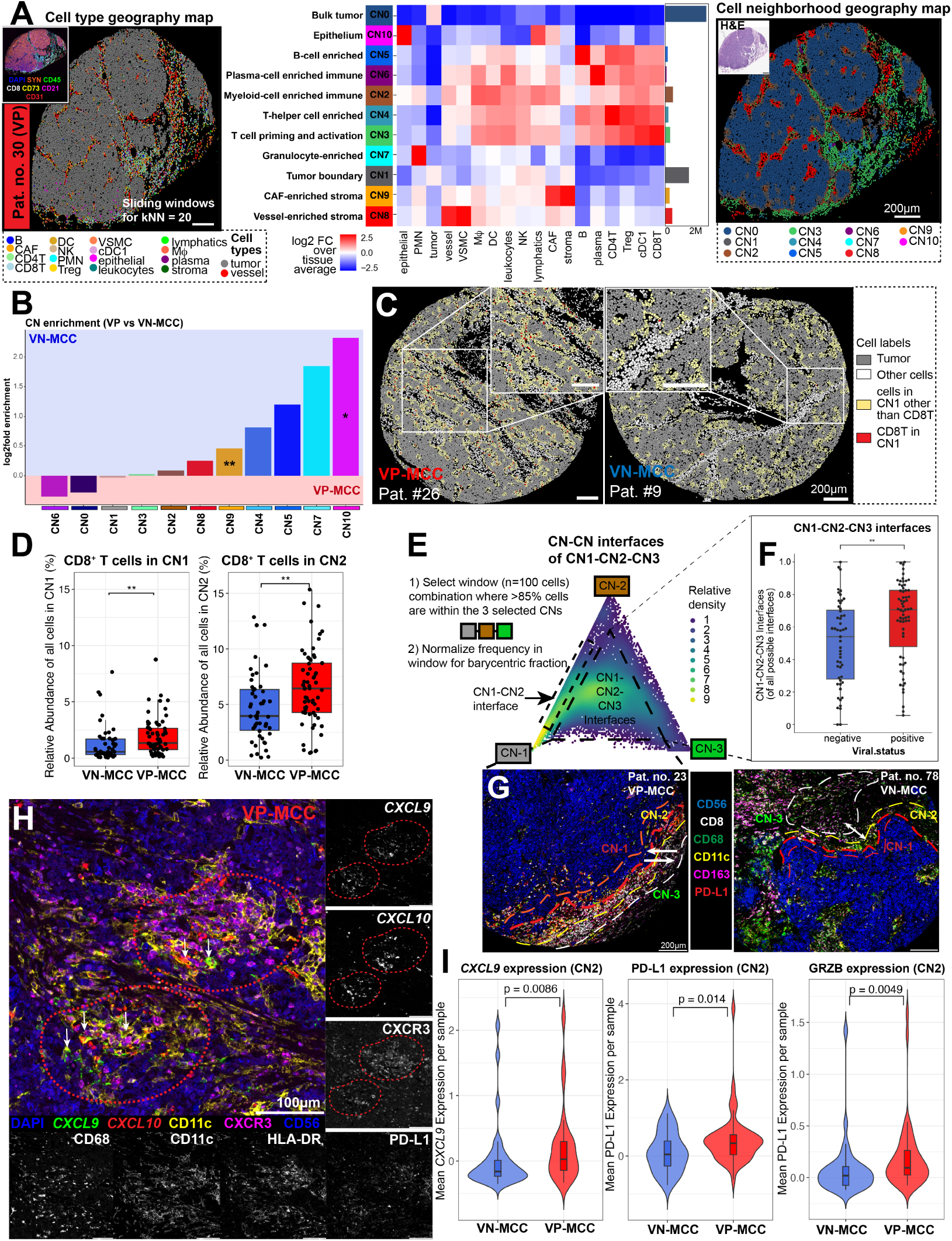
Spatially-restricted expression of immunoregulatory molecules associates with enhanced CD8^+^ T cell infiltration at the tumor invasive front of VP-MCC. **(A)** Left: Cell-type geography map. Center: Identification of 11 distinct CNs based on the 18 granular cell types and their enrichment within each CN (aggregated data from both VN and VP-MCC). Right: CN geography map. **(B)** Mean Log2-fold enrichment of CNs between VP-MCC and VN-MCC tumor samples (n=117). Statistical significance was determined using t-tests adjusted for multiple comparisons using the Bonferroni method with a targeted FDR at 0.05; * *p ≤* 0.05, ** *p ≤* 0.01. **(C)** Representative examples for the spatial distribution of CD8+ T cells within the MCC-TME in VP and VN-MCC. Scale bar of 200µm applies to both panels. **(D)** Relative enrichment of CD8^+^ T cells within CNs at the tumor boundary (CN1, left) and the myeloid cell enriched CN2 (right). Statistical significance was determined using Wilcoxon-rank sum test corrected for multiple-hypothesis testing (BH). ** *p ≤* 0.01. **(E)** Left: Schematic of steps used to generate barycentric plots. Right: Barycentric coordinate projection of the interaction of the invasive tumor CNs CN1, CN2, and CN3 across all tumor samples depicted as density plot. **(F)** Degrees of intermixing amongst CN1, CN2, ad CN3 in VP-MCC vs VN-MCC samples. Each point represents the mean degree of intermixing per MCC tumor sample (n=117). Statistical significance was determined by t-test was adjusted for multiple comparisons using the Bonferroni method with a targeted FDR at 0.05; ** *p ≤* 0.01. **(G)** Representative examples of intersection of CN1-CN2 and CN-3 at the tumor invasive front in VP-MCC and VN-MCC. Scale bars, 200 µm. **(H)** Representative example of CXCL9+ / CXCL10+ DC-T cell aggregates at the tumor border as shown through combined CODEX and multiplex RNAscope imaging within a VP-MCC tumor core. Scale bar of 100µm applies to all panels. **(I)** Violinplots depicting the mean expression of immunomodulatory molecules within the myeloid-cell enriched CN2 across VN and VP-MCC samples. Wilcoxon-rank sum test corrected for multiple hypothesis testing (BH) was used to determine statistical significance. *Abbreviations: CN, cellular neighborhood*.

We used this panel to stain and image 356 tumor cores with a total of 7.9M cells. Following tissue extraction, cell segmentation, quality control, and data normalization as described previously^9,10^, we performed Leiden-based clustering that enabled the identification of 7 major cell lineages (**Fig.1F**) including T and B-cells, myeloid cells, stromal cells, as well as epithelial and tumor cells that comprised the majority of cells within the MCC-TME (**Fig.1G**). We observed a strong heterogeneity in the cellular composition of the investigated MCC tumors but did not detect a clear correlation between the abundance of the major cell lineages with clinical parameters such as tumor stage or viral status (**Fig.1H**).

### The TME of virus-positive MCC is characterized by a CD8-dominant T cell infiltrate exhibiting an effector phenotype while CAFs are enriched within the VN-MCC TME

To better characterize the cellular composition of the MCC-TME we used key phenotypic markers to identify 18 granular cell types (**Fig.2A**). Stratification by MCPyV status highlighted differences in the cellular composition between VP-MCC and VN-MCC tumors (**Fig.2B**): In particular, we observed a higher proportion of tumor cells within VP-MCC, whereas VN-MCC exhibited a higher proportion of stromal cells (**Fig.2C**). To enable a more balanced representation of cellular changes driving anti-tumor immune responses within the MCC-TME, we analyzed the cellular composition of the immune and stroma compartment between VN and VP tumors. We discovered that CD8^+^ T cells and plasma cells were significantly enriched in VP-MCCs, whereas we detected a higher abundance of cancer-associated fibroblasts (CAFs) within VN-MCC tumors (**Fig.2D**). We further interrogated the T-cell compartment, to characterize the differential impact of MCPyV-infection on the abundance of T-cell subtypes (**Fig.2E**) and their functional polarization in more detail. We found that both CD4^+^ and CD8^+^ T-cells within the VP-TME exhibited an elevated expression of proteins, implicated in T-cell cytotoxicity and effector function, including Granzyme B, PD1 and LAG3 (**Fig.2F-G**). To map the organization of different T cell states within the TME, we identified states of T cell differentiation based on previously defined marker expression **(Supplemental Fig.4A**)^11^. Using these T-cell subsets we found an enrichment for both CD4^+^ and CD8^+^ T cells that expressed high levels of the checkpoint molecules (**Fig.2H-I**) in VP-MCC. This observation reflects on the known role of chronic viral infections and cancer in promoting T-cell exhaustion^11^. We expanded our results by analyzing a publicly available scRNAseq dataset^12^ (**Supplemental Fig.4B-D**). In line with the results from our CODEX multiplex imaging, we identified therein a subset of T-cells enriched for genes implicated in T-cell activation and exhaustion (**Supplemental Fig.4E**). We detected a relative enrichment of these PD1^high^, LAG3^high^CD8^+^ T cells in VP-MCC (**Supplemental Fig.4F**) that were characterized by an elevated expression of cytotoxic markers such as *PRF1* and *IFNG*, as well as immune-checkpoint molecules *PDCD1*, *CTLA4* and *TIGIT* (**Supplemental Fig.4G-H**).

**Fig. 4:**
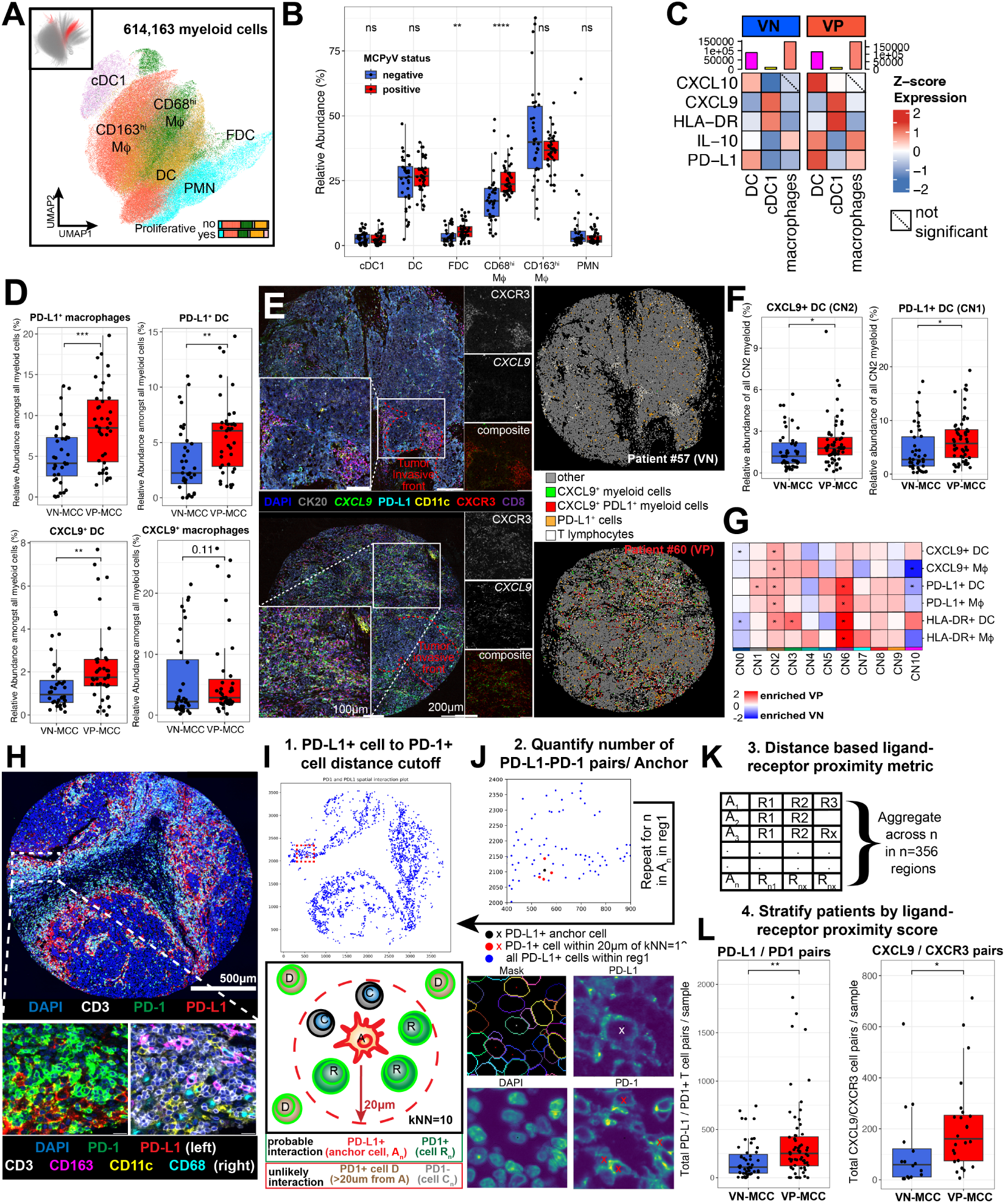
Spatial organization of CXCL9^+^ PD-L1^+^ macrophage and DC aggregates with CD8^+^ T cells at the tumor invasive front. **(A)** UMAP projection of myeloid cell subtypes with relative frequency of Ki67^+^ myeloid cell subtypes (bottom right). **(B)** Comparison of the myeloid cell type frequencies normalized to the overall myeloid cell counts stratified by MCPyV-status. Each point represents the mean cell frequency per MCC tumor sample (n=117). Significance was determined by Bonferroni-corrected t-test comparisons with FDR < 0.05. ∗*p* < 0.05, ∗∗*p* < 0.01. **(C)** Heatmaps depicting mean z-normalized expression of functional markers across myeloid cell subtypes between VP and VN-MCC samples. Mean expression values of each marker in a given cell-type were compared between VP and VN-MCC samples using Wilcoxon-rank sum test adjusted for multiple-hypothesis testing (BH). Non-significant interactions are indicated by dotted lines. **(D)** Boxplots depicting relative abundance of inflammatory myeloid cell subtypes in VP-MCC vs VN-MCC. Each point represents the mean cell frequency from all investigated MCC-samples (n=117). Significance was determined by Bonferroni-corrected t-test with FDR < 0.05. * *p ≤* 0.05, ** *p ≤* 0.01. **(E)** Representative examples from combined CODEX and RNAscope imaging for the CXCL9^+^ myeloid cells (green) located at the invasive tumor front that colocalize with PD-L1^+^ myeloid cells (red) and are in proximity to T cells (black). Scale bars, 200 µm. **(F)** Boxplots depicting relative abundance of CXCL9+ / PD-L1+ DCs within CNs at the tumor boundary (CN1, CN2) in VP-MCC vs VN-MCC. Each point represents the mean cell frequency from all investigated MCC-samples (n=117). Significance was determined by Bonferroni-corrected t-test with FDR < 0.05. * *p ≤* 0.05. **(G)** Relative enrichment of marker-positive myeloid cell subtypes across CNs. Asterisks indicate CNs and cell types with a regression p-value < 0.05 (not adjusted for multiple testing). Red boxes highlight the enrichment of inflammatory CXCL9^+^, HLA-DR^+^, PD-L1^+^ macrophages and DC within CNs of the invasive tumor front (CN1-3). **(H-L)** Schematic step-by-step approach used to quantify receptor-ligand interactions within the TME. This includes the identification of PD1^+^ cells within 20µm of the kNN=10 of a given PD-L1^+^ anchor cell **(I)**, the quantification and visual confirmation of PD1^+^ cells in proximity to PD-L1^+^ anchor cell **(J)**, the quantification of ligand-receptor pairs across all anchors within a region and aggregation over all investigated regions **(K)**. This metric was used to stratify all MCC samples (n=117) by the number of PDL1/PD1 pairs per tumor core based on viral status. Boxplots comparing the number of PDL1/PD1 and CXCL9/CXCR3 ligand-receptor pairs are shown in L including results of BH-adjusted Wilcoxon rank sum test **(L)**. ∗*p* < 0.05, ∗∗*p* < 0.01.

### Neighborhood analysis reveals a spatial co-enrichment of immunomodulatory CXCL9 and CD8^+^ T cells at the invasive front of VP-MCC

In order to spatially contextualize the enrichment of CD8^+^ T cells and their functional states within the VP-MCC TME, we grouped the TME into spatially organized functional units, termed cellular neighborhoods (CNs). We identified 11 unique CNs characteristic of VP-MCC and VN-MCC using a k-nearest neighbors’ approach (**Fig.3A** and **Supplemental Fig.5A-D**). While some CNs were unique to a given tissue, such as an *epithelium CN* (skin) or *LN-follicles* (LN), the majority of CNs was shared between tissues, such as a B-cell enriched CN, vessel-enriched stroma, and more granular CNs such as T cell priming and activation zones and the tumor boundary (**Supplemental Fig.5E-H**).

**Fig. 5:**
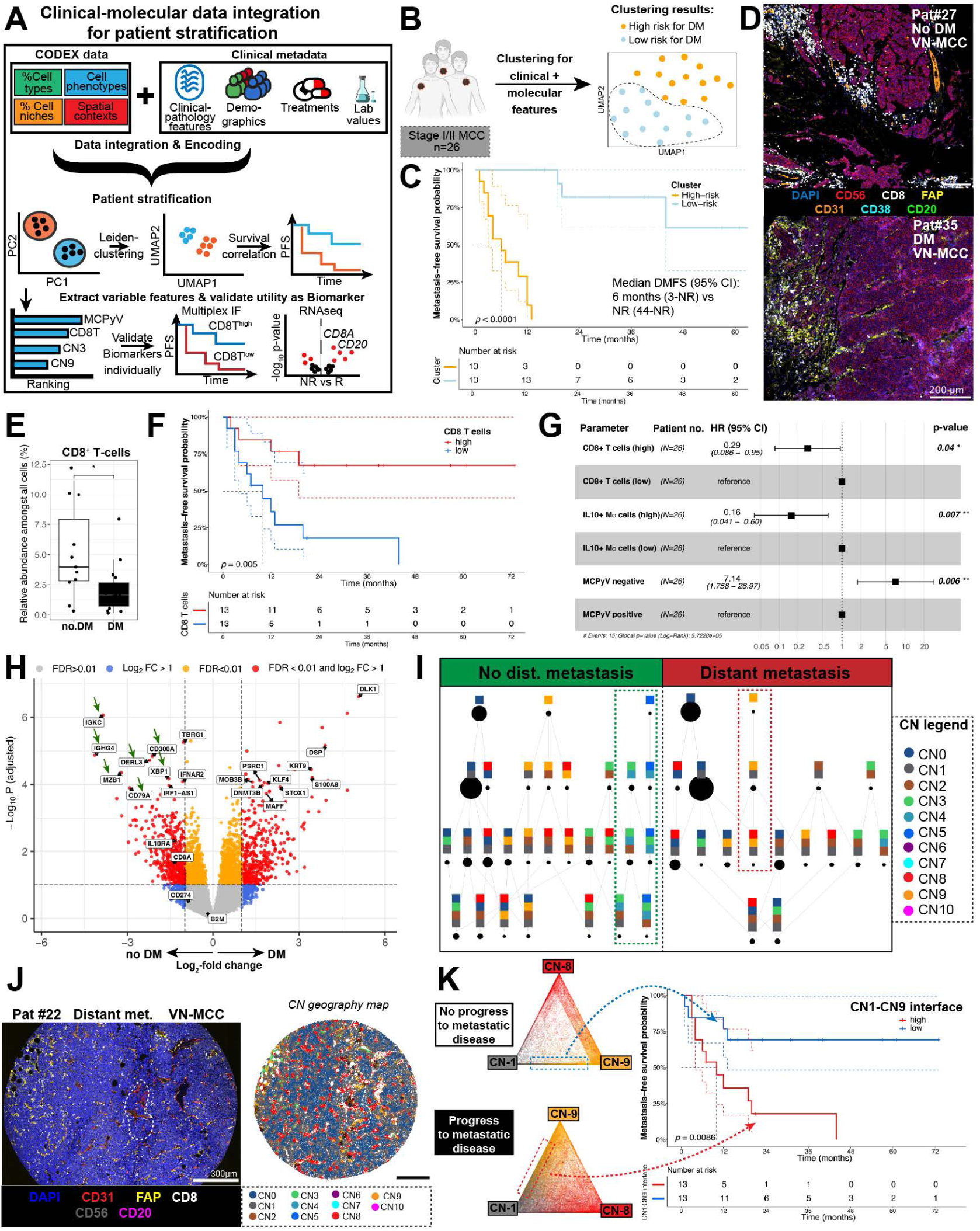
MCPyV-positivity is linked to prolonged distant-metastasis-free survival in local stage MCC through enhanced CD8^+^ T cell infiltration. **(A)** Analytical framework for the identification of biomarkers associated with distant metastasis. **(B)** UMAP representation of identified patient clusters separating patients with local stage I/II MCC into high- and low-risk (demarcated by dotted line) groups for distant metastasis. **(C)** Kaplan-Meier survival plot for distant-metastasis-free survival stratified by high/low risk clusters were based on combined clinical and molecular patient features. **(D)** Representative images of two MCC samples from patients without and with distant metastasis (DM). Scale bars, 200µm. **(E)** Boxplot depicting CD8^+^ T frequencies in all patients stratified by occurrence of distant metastasis. Each point represents the mean cell frequency from all TMA cores sampled before distant metastasis per patient (n=26). **(F)** Kaplan-Meier plot for DMFS in patients that were dichotomized by median CD8^+^ T infiltration. Statistical significance was determined by log-rank test. **(G)** Lasso-regressed multivariate CoxPH model depicting variables significantly associated with DMFS amongst the collected clinical and molecular parameters from CODEX multiplex imaging. **(H)** Volcano plot illustrating genes significantly enriched in in patients with or without distant metastasis at a false-discovery rate (FDR) < 0.01 as assessed by LCM-seq. Green arrows indicate genes implicated in B-cell-driven immune responses or T cell activation. **(I)** Spatial context map revealing CN-CN interfaces stratified by patients with and without distant metastases. Dotted lines indicate unique spatial contexts that were only observed in patients with (center) or without (left) distant metastasis. Color-codes specified on the right. **(J)** Representative fluorescence image (left) with corresponding CN geography map (right) illustrating the spatial context involving the interaction of vessel-enriched CN8 with a CAF-enriched CN9 at the tumor boundary CN1. **(K)** Barycentric coordinate projection depicting the interactions between CN1-CN8-CN9 in patients without (top left) or with distant metastasis (bottom left), as well as corresponding Kaplan-Meier plot for DMFS in patients with high and low levels of CN1-CN9 intersection (right). Statistical significance was determined through log-rank test. *Abbreviation: DM, distant metastasis*.

We compared the abundance of CNs between VP-MCC and VN-MCC and observed a higher abundance of the *CAF-enriched stroma* CN9 in VN-MCC (**Fig.3B**). More importantly, we found that CD8^+^ T cells were particularly enriched in CN1 (*tumor boundary*) and CN2 (*myeloid-cell-enriched*) of VP-MCC (**Fig.3C-D, Supplemental Fig.6A**). This indicates that MCPyV-presence does not result in a homogenous enrichment of CD8^+^ T cells but that CD8T cell recruitment is confined to restricted tumor niches. When further analyzing the distribution of CD8^+^T-cell subclusters (**Supplemental Fig.6B-C**), we found a relative enrichment of PD1^high^, LAG3^high^ CD8^+^ T cells within the tumor compartment of VP-MCCs (**Supplemental Fig.6D**). To quantify whether the regionalized CD8^+^ T cell infiltration correlates with the organization of CN-CN interactions at the tumor invasive front, we computed the intersections of CNs located at the tumor invasive front (CN1, CN2 and CN3) (**Fig.3E**). Regions of such CN-CN interactions, that are termed spatial contexts, are locations where the characteristic local processes of each CN are interacting through local molecular reactions. For example, CN-CN interfaces are sites where a molecule from one CN contacts cells from another CN that enables unique activation processes^13^. We observed that the spatially restricted influx of CD8^+^ T cells into CN1 (*tumor boundary*) and CN2 (*myeloid-cell-enriched*), associated with a stronger intermixing of CN1 and CN2 with the CN3 (*T-cell-enriched*) in VP-MCC (**Fig.3F-G**).

**Fig. 6:**
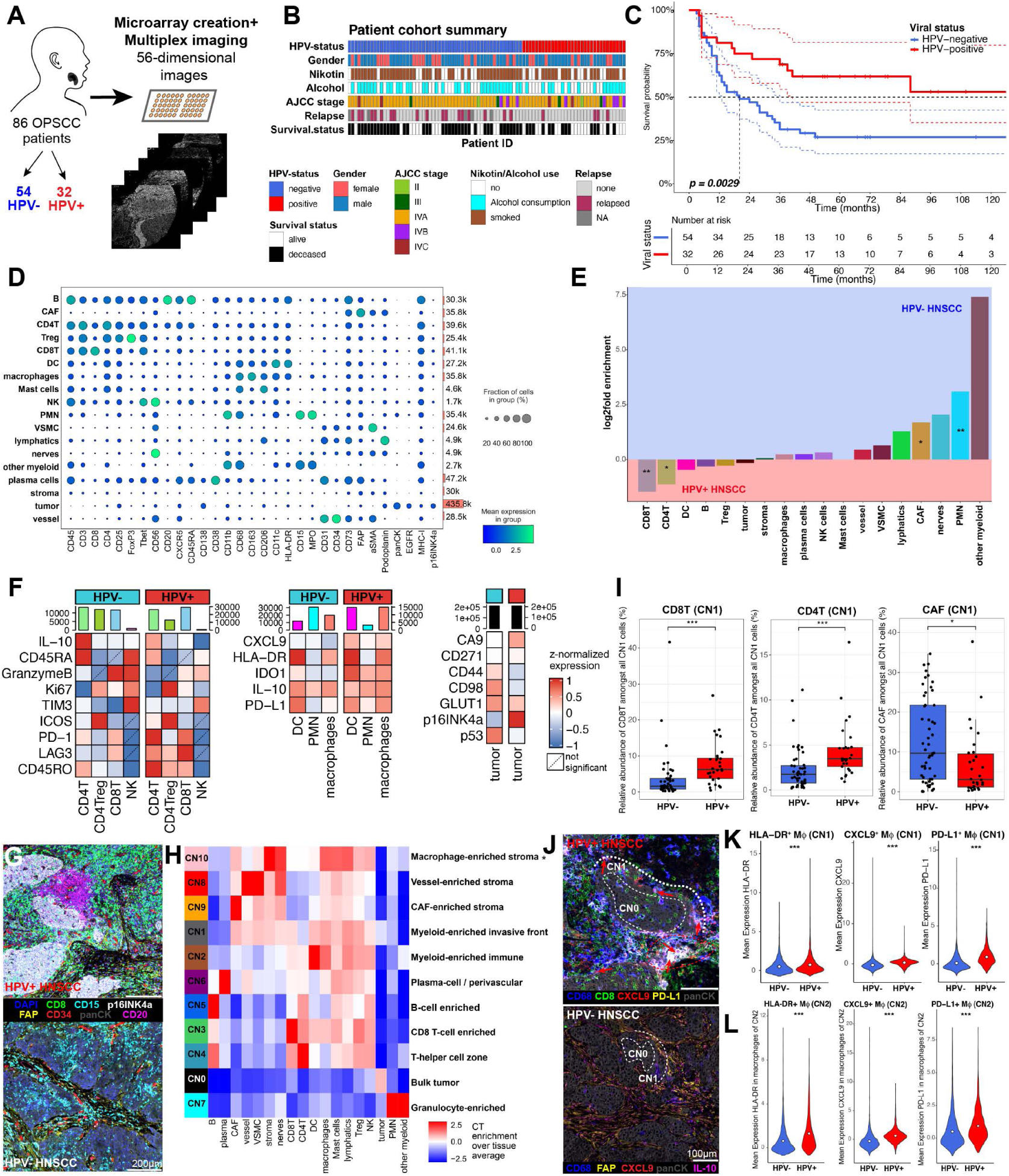
CXCL9+ macrophages and CD8^+^ T cells are spatially co-enriched at the tumor invasive front of HPV-positive head-and-neck carcinoma. **(A)** Conceptual and analytical framework for the investigation of the OPSCC patient cohort using CODEX multiplex imaging. **(B)** Summary of clinical patient characteristics of the investigated OPSCC patient cohort. **(C)** Kaplan-Meier survival plot for overall survival stratified within the OPSCC cohort separated by HPV-viral status. Statistical significance was determined using log-rank test. **(D)** Dotplot summarizing the mean log-normalized expression values of key phenotypic and functional markers amongst the classified cell-types within the CODEX dataset. **(E)** Waterfall plot depicting the mean log2-fold enrichment of major cell-types in HPV+ vs HPV-OPSCC patients. Statistical significance was determined using Wilcoxon-rank sum test corrected for multiple-hypothesis testing (BH). ∗*p* < 0.05, ∗∗*p* < 0.01. **(F)** Left: Heatmaps comparing the mean z-normalized expression of functional markers in selected cell-types between HPV- and HPV+ OPSCC samples, with corresponding total cell counts of each cell-type shown as barplots above. Non-significant differences in functional marker expression for a given cell-type between viral-status groups are indicted by dotted lines. Statistical significance was determined using Wilcoxon-rank sum test corrected for multiple hypothesis testing. **(G)** Representative examples illustrating the cellular composition of the TME in HPV+ vs HPV-OPSCC. Scale bars, 200µm. **(H)** Heatmap depicting the relative enrichment of cell-types within the identified OPSCC CNs compared to the global tissue average. CNs were computed using window sizes of k=15 nearest neighbors. **(I)** Boxplots depicting the relative enrichment of CD8+ and CD4+ T cells in the tumor boundary CN1 of HPV+ OPSCC (left and center); while CAFs were enriched in CN1 of HPV-OPSCC (right). Statistical significance was determined using Wilcoxon-rank sum test corrected for multiple hypothesis testing (BH). ∗*p* < 0.05, ∗∗*p* < 0.01, *** *p*<0.005. **(J)** Representative images depicting the enrichment of CD8+ T cells and CXCL9+ macrophages in CN1 of HPV+ HNSCC while in HPV-OPSCC CAFs were enriched in the tumor boundary CN1. Scale bar of 100µm applies to both panels. **(K)** Violinplots comparing the mean log-normalized expression of HLA-DR, CXCL9 and PD-L1 in macrophages of the tumor boundary CN1 between HPV+ and HPV-OPSCC. Statistical significance was determined using Wilcoxon-rank sum test corrected for multiple hypothesis testing (BH). ∗*p* < 0.05, ∗∗*p* < 0.01, *** *p*<0.005. **(L)** Violinplots comparing the mean log-normalized expression of HLA-DR, CXCL9 and PD-L1 in macrophages of the tumor boundary CN2 between HPV+ and HPV-OPSCC. Statistical significance was determined using Wilcoxon-rank sum test corrected for multiple hypothesis testing (BH). ∗*p* < 0.05, ∗∗*p* < 0.01, *** *p*<0.005.

We reasoned that the stronger infiltration of CD8^+^ T cells into CNs of the invasive front might be linked to the presence of immunomodulatory molecules within those CNs. To quantify the distribution of both immunomodulatory surface markers and soluble mediators, we established a protocol in which we combined both multiplex RNA *in-situ hybridization* with CODEX multiplex imaging (**Fig.3H, Supplemental Fig.6E-G**, **METHODS**). We carried out cell-type annotations and CN identification of this combined CODEX-RNAscope experiment similar to the previously described CODEX experiment (**Supplemental Fig.6H-J**). Using this orthogonal dataset we found a strong co-localization of PD-L1^+^ and CXCL9^+^ expression in cells at the tumor invasive front (CN1, CN2, CN3). Critically, we also found a significant enrichment of the immunomodulatory molecules PD-L1 and *CXCL9*, as well as T-cell cytotoxicity marker Granzyme B in CN2 (*myeloid-cell-enriched*) of the tumor invasive front of VP-MCC tumors (**Fig.3I, Supplemental Fig.6K**). This indicates that those CNs might exhibit higher levels of T-cell activation and recruitment in VP-MCC. Notably, immunomodulatory molecules *CXCL9, CXCL10* and PD-L1 were predominantly co-localized within cells of myeloid origin, specifically macrophages and DC that were in close proximity to T-cell aggregates (**Fig.3H, Supplemental Fig.7B**), indicative of a central role of these myeloid cells in the spatial organization of T-cell responses within the MCC-TME.

**Fig. 7:**
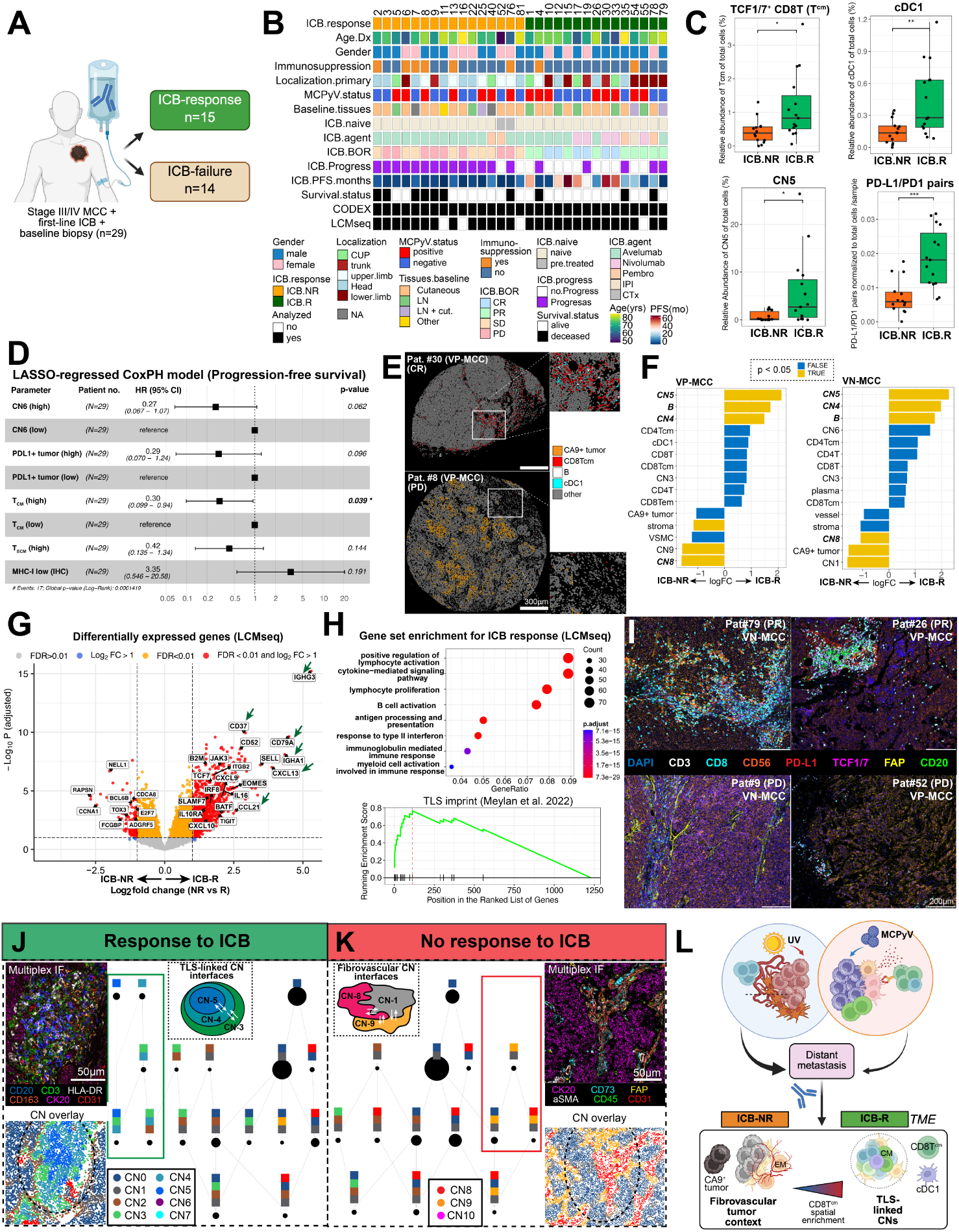
Predominance of central memory CD8T-cells and B-cell enriched niches are shared features predicting response to ICB in VP and VN-MCC. **(A)** MCC patient cohort used for investigation of predictive molecular biomarkers. **(B)** Summary of clinical patient characteristics of the investigated OPSCC patient cohort. **(C)** Boxplots depicting relative abundance of molecular features associated with ICB-response. Statistical significance was determined using Wilcoxon-rank sum test corrected for multiple-hypothesis testing (BH). ∗*p* < 0.05, ∗∗*p* < 0.01, *** *p*<0.005. **(D)** Lasso-regressed CoxPH model summarizing features significantly associated with PFS amongst the collected CODEX parameters and clinical parameters. **(E)** Representative cell-type geography maps highlighting enrichment of cellular features associating with ICB response. Scale bars, 300 µm. **(F)** Barplots summarizing the top10 and bottom 5 features sorted by log-fold change difference associating with ICB-response in VP-MCC patients (left) and VN-MCC patients (right). Statistical significance was determined using a generalized linear model from the glm-package in R. **(G)** Volcano plots of selected genes significantly enriched in ICB responders with FDR < 0.01 as assessed by LCM-seq. Genes implicated in TLS formation and antigen presentation pathways are highlighted. **(H)** Dotplot summarizing results gene-set enrichment analysis for ICB-responders with selected non-redundant top-20 gene sets shown (top). Bottom gene-set enrichment plot using the TLS imprint signature described in Meylan et al. ^21^. **(I)** Representative images highlighting characteristic TME composition and architectural features across VP and VN-MCC, that associate with ICB response. Scale bars, 200 µm. **(J)** Left: Representative multiplex immunofluorescence image of the spatial contexts unique to ICB-responders with corresponding CN-overlays. Right: Spatial context map depicts the enrichment of characteristic CN-interfaces in ICB-responders including the CN3-CN4-CN5 spatial context (highlighted in green box). Size of nodes depicts the relative abundance of these CN-CN interfaces amongst the overall dataset. **(K)** Left: Spatial context map depicts the enrichment of characteristic CN-interfaces in ICB-non-responders including the spatial context encompassing CN1-CN8-CN9 interfaces (highlighted in red box). Right: Representative multiplex IF image of the spatial context specified on the left unique to ICB-non-responders with corresponding CN-overlays. Color-codes as specified in Figure 7J-K apply. **(L)** Cartoon depicting key differences between the VN and VP-MCC TME that results in differential risk for distant metastasis (top). Despite these distinct TME compositions, VN and VP-MCC share cellular and architectural features that predict ICB response (bottom). *Abbreviations. PD, progressive disease; PR, partial response; CR, complete response; R, response; NR, no response; ICB, immune checkpoint blockade; Tcm, central memory CD8T cells; PFS = progression-free survival; cDC1 = conventional dendritic cell type 1*.

### CXCL9^+^PD-L1^+^ macrophages and DC spatially co-localize with CD8^+^ T cells at the tumor front that associates with enhanced receptor-ligand interactions in VP-MCC

To better characterize the phenotype of these myeloid cells, and their interaction with CD8^+^ T cells within the MCC-TME, we analyzed the myeloid compartment in more detail. The myeloid compartment is known to show high expression of PD-L1 and has been implicated in T cell priming and recruitment as reflected by its spatial location at the interface of the tumor boundary and T cell CNs^14,25,26^, but little data exist on their role in MCC. In our CODEX dataset, we detected six clusters of myeloid cells annotated as DC, conventional DC type 1 (cDC1), follicular DC (FDC), CD68^hi^ and CD163^hi^ macrophages, and neutrophils (PMN) clusters (**Fig.4A**). Each of these clusters revealed a distinct expression pattern of functional markers (**Supplemental Fig.7A, supplemental Table 5**) that was in agreement with previous reports from scRNAseq data^15,16^. We observed an enrichment of CD68^hi^ macrophage and FDC clusters (**Fig.4B**) and elevated expression of immunoregulatory molecules within myeloid cell subtypes of VP-MCC compared to VN-MCC (**Fig.4C**), suggesting that the functional phenotype of myeloid cells is substantially altered within the VP-TME.

When characterizing these changes in more detail, we found a global enrichment of PD-L1^+^ and HLA-DR^+^ macrophages and DC in VP-MCC (**Fig.4D**). Spatially, those myeloid cells were predominantly located at the tumor invasive front (**Fig.4E**): In line, CXCL9^+^/PD-L1^+^/HLA-DR^+^ macrophages and DC were significantly enriched in CNs at the tumor-invasive front (CN1-CN3) of VP-MCC (**Fig.4F-G**), implying that the VP-MCC TME might show a stronger interaction of PD-L1^+^ and CXCL9^+^ macrophages and DC with PD1^+^ and CXCR3^+^ T cells. Indeed, distance-based interaction analysis showed that PD-L1^+^ macrophages and DC were more likely to interact with PD1^+^ T cells compared to their ligand-negative counterparts (**Supplemental Fig.7C**). Similarly, we observed a significantly stronger interaction between CXCL9^+^ myeloid cell subtypes and CXCR3^+^ T cells compared to their CXCL9^-^ counterparts (**Supplemental Fig.7D**). To account both for the distance and the direct neighbors of a cell, we developed a novel analytical tool that enabled the identification of all PD1-receptor^+^ T cells within a given distance of a PD-L1^+^ anchor cell (**Fig.4H, METHODS**). We computed the number of PD-L1/PD1 pairs for all PD-L1^+^ anchor cells within a given tumor sample (**Fig.4I-J**) and finally aggregated this distance-based metric across all investigated tumor samples (**Fig.4K**), thereby quantifying the potential for direct receptor-ligand interactions between PD-L1^+^ cells and PD1^+^ cells within the MCC TME. Using this distance-based metric we observed a higher degree of PD-L1 engagement with PD1^+^ cells in VP-MCC than in VN-MCC that was independent from the total number of PD-L1^+^/PD1^+^ cells (**Fig.4L** left**, Supplemental Fig.7E**). Similarly, we observed an enrichment of *CXCL9^+^* macrophages/DC pairs with CXCR3^+^ T cells (**Fig.4L**, right). Given the role of CXCL9 in T-cell recruitment^17^, our findings suggest that the enrichment of mature (HLA-DR), inflammatory (PD-L1, CXCL9) macrophages and DC in VP-MCCs might be involved in the enhanced T-cell recruitment into VP-MCC. In line, we detected significant correlations between the relative abundances of PD-L1^+^ and CXCL9^+^ macrophages and DC with CD8^+^ T cells (**Supplemental Fig.7G**). While PD-1/PD-L1 and CXCL9/CXCR3 pairs predominantly formed within immune-cell centric CNs (CN2-4) in both VP and VN-MCC (**Supplemental Fig.7H**), we observed that within tumor-centric CNs (CN0-CN2) the formation of CXCL9-CXCR3 pairs was significantly elevated in VP-MCC as compared to VN-MCC (**Supplemental Fig.7I**), suggesting a spatial link of inflammatory myeloid cell enrichment and T-cell recruitment into the tumor of VP-MCCs. These findings were reflected by the scRNA-seq dataset, which confirmed elevated expression of *CXCL9-CXCL11*, and *HLA-DR* in myeloid cells of VP-MCCs (**Supplemental Fig.7J-K**).

Collectively, these observations support a model of MCPyV-driven TME immunomodulation where MCPyV-infection is linked to the polarization of macrophages and DC towards an inflammatory CXCL9^+^ and PD-L1^+^ phenotype particularly in CNs at the invasive front. The spatially restricted enrichment of these inflammatory myeloid cells is associated with an enhanced infiltration of CD8^+^ T cells and particularly effector CD8^+^ T cells into the VP-TME. Specifically, we find an enhanced interaction of mature HLA-DR^+^ and inflammatory CXCL9^+^PD-L1^+^ macrophages and DC with early activated PD-1^+^ CD8^+^T cells in VP-MCC. Meanwhile, CD8^+^ T cells with elevated expression of PD1 and LAG3 were predominantly enriched in the tumor compartment of VP-MCC (**Supplemental Fig.6D**), indicative of a tumor-specific T-cell response in this compartment. By contrast, VN-MCC is characterized by a CD4-dominant T cell infiltrate with a stronger infiltration of CD163^hi^ macrophages and a selective enrichment of CAFs within the TME (**Supplemental Fig.8**). Importantly these observations were even more pronounced within cutaneous tissue samples only (**Supplemental Fig.9**), which agrees with the skin as primary site of MCPyV-infection.

### MCPyV-positivity and enhanced CD8^+^ T-cell infiltration are linked to prolonged distant-metastasis-free survival in local stage MCC

We next investigated which features of the MCPyV-specific immune signatures might be linked to the differential survival outcomes of VP-MCC and VN-MCC patients with the aim to identify biomarkers associated with metastasis-free survival. We employed *ehrapy^18^* to integrate the clinical and molecular data available at initial MCC diagnosis, which allowed us to account for the heterogeneity of clinical and high-dimensional molecular data from multiplex imaging analysis (**Fig.5A**). Following feature-based clustering that enabled the stratification of patients into distinct clusters, we identified cluster-defining features and explored their association with DMFS (**Supplemental Fig.10A-C**).

This strategy separates two patient clusters that associated with DMFS, which we hence classified as high and low risk (**Fig.5B-C**). These clusters associated with MCPyV status on a clinical level. On a molecular level we found that the abundance of CD8^+^ T cells, CN3 (*T-cell-enriched*), plasma cells, and IL10^+^ myeloid cells associated with prolonged DMFS (**Fig.5D-F** and **Supplemental Fig.10D, F**). Strikingly, we found that specifically the enrichment of CD8^+^ T cells in CN3 (*T-cell-enriched*) associated with lower risk to metastasis, thereby highlighting the relevance of the spatial context of cells on patient outcomes (**Supplemental Fig.10E**). To identify independent determinants of DMFS, we applied a lasso-regressed Cox proportional hazards (CoxPH) model, which enabled us to study of the relationships between multiple molecular predictor variables and survival time^19^. The CoxPH model confirmed MCPyV-positivity as independent determinant of DMFS (HR for VN-MCC: 7.14, 95% CI: 1.76-29.0, *p<*0.006) and highlighted the association of CD8^+^ T cell infiltration with low risk of distant metastasis (**Fig.5F**). In agreement, LCMseq confirmed an enrichment of transcripts implicated in T cell-based immune responses (**Supplemental Fig.10G-H**), as well as the upregulation of transcripts associated with B-cell driven immune responses, including *IGHG4* and *CD79A* within the MCC-TME (**Fig.5H, Supplemental Fig.10G-H**). This transcriptional signature has previously been linked to the formation of tertiary-lymphoid structures (TLS) ^20,21^, which frequently associate with improved patient outcomes. To identify such structures within our multiplex imaging dataset and determine their importance in predicting DMFS, we investigated the MCC-TME of patients with initial local disease on a tissue architecture level: This approach revealed unique spatial contexts that associated with DMFS (**Fig.5I**). These spatial contexts are defined by the intersection of CNs within a given window size and included the intersection of T-and B-cell enriched niches (CN3, CN4, CN5) resembling TLS in structure and composition (**Fig.5I**, left; **Supplemental Fig.10I**), which we found specifically in patients without distant metastasis. By contrast, a stronger intersection of CN9 (*CAF-enriched*) with CN1 (*tumor boundary*) correlated with inferior DMFS in these patients (**Fig.5I** right, **Fig.5J-K)**, thereby providing a spatial correlate for the inferior outcomes observed in VN-MCC patients (**Supplemental Fig.10J**). In agreement, the relative abundance of CN1-CN9 interfaces negatively correlated with the infiltration of CD8^+^ T-cell infiltration highlighting the adverse role of this CAF-tumor-context in MCC (**Supplemental Fig.10K**). Our observations also align with a previously described xenograft model in which patient-derived CAF and the MCC cell line MKL-1 were co-injected into SCID mice. In that study, the co-injection of CAF and MKL-1 cells led to a significantly higher incidence of distant metastases compared to the injection of MKL-1 cells alone or in combination with skin-derived fibroblasts^22^. To better characterize the phenotype of these CAFs we re-analyzed tumors of this study using CODEX multiplex imaging (**Supplemental Fig.10L**). In line with our human MCC-CODEX data (**Supplemental Fig.10M**), we observed that specifically xenografts derived from CAF-MLK1 co-cultures that were enriched in CAFs at the tumor boundary were more likely to metastasize. Furthermore, the CAF-enriched niches in these metastatic xenografts showed a higher expression of immunomodulatory CD73 (**Supplemental Fig.10N**), that has previously been linked to adverse clinical outcomes^23^.

Collectively, these data provide a link between our proposed model of MCPyV-driven immunomodulation of the MCC-TME and differential patient outcomes through the selective enrichment of CD8^+^ T cells within the TME: Furthermore, we provided both molecular and spatial features that allow for the identification of a group of patients at high risk for metastasis including virus-negativity, low CD8^+^ T-cell infiltration and a CAF-enriched tumor boundary that associates with poor DMFS.

### Spatially coordinated enrichment of inflammatory myeloid cells with CD8^+^T-cells at the tumor invasive front is conserved in HPV^+^ head-and-neck cancer

Given the link of MCPyV-driven immunomodulation and patient outcomes, we interrogated if our model of viral-linked spatial immunomodulation might also be observed in other virus-associated cancers. To achieve this, we investigated the TME from primary tumor tissues of a cohort of 86 patients with metastatic oropharyngeal squamous cell carcinoma (OPSCC) and known status of human papillomavirus infection (HPV) using CODEX multiplex imaging (**Fig.6A**). Next to classical risk factors such as alcohol and tobacco use, HPV has emerged as another risk factor for OPSCC. Notably, HPV^+^ OPSCC are characterized by a distinct molecular profile compared to its HPV-negative counterpart^24^. HPV^+^ and HPV^-^ patients were matched for main clinical parameters such as demographics, primary tumor characteristics, tumor stage, treatments and follow-up (**Fig. 6B**, for details see^25^). In line with prior data, we observed a significantly better prognosis in HPV-positive OPSCC patients compared to their HPV-negative counterparts (**Fig.6C**). Similar to our MCC cohort, we constructed a 56-marker panel for CODEX multiplex imaging, which we used to interrogate the TME in this OPSCC cohort (**Supplemental Fig.11A**, **METHODS**). Following initial cell-type annotation using major cell type markers (**Fig.6D**), we compared the cellular composition of the TME outside of the tumor compartment stratified by HPV-status. Similar to the observations from our MCC-cohort (**Supplemental Fig.8**) we observed a significant enrichment of CD8^+^ T cells in HPV^+^ OPSCC, whereas CAFs where enriched within the TME of HPV-patients (**Fig.6E**). More specifically, we found that CD8^+^ T cells within the HPV^+^ TME exhibited elevated levels of immune-checkpoint molecules including PD1, LAG3 and TIM3 (**Fig.6F**, left; **Supplemental Fig.11B**). In line with our previous observations from MCPyV+ MCC, we also detected an enrichment of immunomodulatory IDO1, PD-L1 and CXCL9 expression in macrophages and DC from HPV+ OPSCC (**Fig.6F**, center).

To better characterize the spatial organization of HPV-linked anti-tumor immune responses within the OPSCC-TME, we identified 11 CNs based on the cellular composition within the k=15 nearest-neighbors (**Fig.6G-H**). In accordance with our findings from MCC tumors, we found that HPV-OPSCC tumors were characterized by the relative enrichment of a CAF-enriched CN (**Supplemental Fig.11C**). More importantly, we again observed that T-cells were specifically enriched within CNs at the tumor invasive front (CN1) in HPV+ OPSCC (**Fig.6I**, **Supplemental Fig.11D**). In agreement with our findings in VP-MCC, we observed a spatial co-enrichment of CXCL9^+^, PD-L1^+^ macrophages and CD8^+^ T cells at the invasive front of HPV+ OPSCC (**Fig.6J**), as well as a significantly elevated expression of immunomodulatory IDO1, PD-L1 and CXCL9 in macrophages and DC within CN1 (*tumor boundary*) and CN2 (*myeloid-cell enriched immune*) of HPV+ OPSCC (**Fig.6K-L, Supplemental Fig. 11E**).

By contrast, HPV-OPSCC was characterized by the enrichment of CAFs at the tumor invasive front (**Fig.6I, J**). This selective enrichment of CAFs at the tumor boundary, was associated with the formation of a distinct spatial context, characterized by the intersection of CN9 (*CAF-enriched*) with CN0 (*Bulk tumor*) and CN1 (*tumor boundary*). Similar to our MCC-cohort, this spatial context associated with an adverse outcome in OPSCC patients (**Supplemental Fig.11F-I**), supporting the notion that the CAF-tumor context might be a shared predictor of adverse outcomes in both viral cancers.

### Central memory CD8^+^ T cells and B-cell linked spatial contexts predict response to ICB across advanced VP and VN-MCC

ICB is the current standard-of-care for advanced MCC patients^26,27^. However, it is not known whether the differential TME organization in VP and VN-MCC might impact the efficacy of the ICB. While the ICB efficacy in advanced MCC has been linked to various clinical and molecular biomarkers^4,28,29^, there is yet a limited understanding if those biomarkers might apply for both in VP or VN-MCC patients. To determine features predictive of ICB response, we investigated a subset of our MCC cohort that received ICB therapy for advanced MCC (**Fig.7A-B**, **Supplemental Fig.12A-B**). We integrated clinical, molecular and architectural features for patients with available baseline tissue biopsies. This approach yielded two clusters, reflecting patients with durable ICB response or ICB failure (**Supplemental Fig.12C-D**).

Patients with durable response to ICB therapy were characterized by the enrichment of a TCF1/7^+^ subset of CD8^+^ T cells (CD8T^cm^) (**Supplemental Fig.12E**), cDC1 and B-cell enriched CNs (CN4, CN5), as well as a higher abundance of direct PD-L1/PD1 pairs within the MCC-TME (**Fig.7C**). To assess the relative importance of these cellular biomarkers in a multivariate model we used a lasso-regressed CoxPH model for progression-free survival (PFS) on those clinical and molecular features. This model recapitulated the relevance of CD8T^cm^ as potential predictors of PFS (**Fig.7D-E**). Notably, MCPyV-positivity, although associated with DMFS, was not associated with ICB response or PFS (**Supplemental Fig.12F-G**). Rather, VP and VN-MCC patients exhibited a shared set of both favorable and adverse predictive features, including the presence of B-cell enriched CNs and central memory T-cells (**Fig.7F**, **Supplemental Fig.12H**) that correlated globally on a patient-wide level (**Supplemental Fig. 12I**).

We orthogonally validated those molecular signatures using LCM-seq, which confirmed an enrichment of genes implicated in immune cell recruitment (*CCL19, CCL21, CXCL13*), T cell memory formation (*TCF7, BATF*), and B cell activation (*IGHG3, CD79A*) amongst ICB-responders (**Fig.7G-H**). As suggested by the cellular and transcriptional signatures, we observed that the enrichment of adaptive immune cells in ICB-responders often occurred in lymphoid-rich clusters resembling TLS in structure and composition (**Fig.7I**).

To better contextualize these interactions on the level of functional tissue units, we computed spatial contexts, that summarize CN-CN intersections within the TME. Using this approach, we detected a unique spatial context amongst ICB-responders characterized by the intersection of T and B-cell enriched niches reminiscent of TLS (**Fig.7J**). By contrast, patients who did not responds to ICB often showed an intersection of the CAF-enriched CN9 with the vessel-enriched CN8 at the tumor boundary CN1 (**Fig.7K**). These higher-order interactions were corroborated on the level of cell-cell interactions, where patients who responded to ICB presented with a stronger interaction of B-cells with DCs and CD4Tcm (**Supplemental Fig.14D**). Notably, the presence of the TLS-context not only correlated with the numeric abundance of CD8T^cm^ (**Supplemental Fig.13A**), but CD8T^cm^ were in fact spatially enriched within the TLS-context (**Supplemental Fig.13B-E**), indicating that T-cell function and activity is differentially regulated by their spatial context. Importantly, we confirmed that these cellular, transcriptional and architectural signatures were not only shared between VP and VN-MCC patients but were also identified when only investigating cutaneous tissue biopsies from patients who received ICB for advanced MCC (**Supplemental Fig.14A-C, and 14E-I**).

Collectively, our data demonstrate that VP and VN-MCC patients share CD8T^cm^ and TLS spatial contexts as features predictive of ICB-response (**Fig.7L),** despite the distinct TMEs of VP and VN-MCC that resulted in a differential risk for distant metastasis in early-stage MCC patients. We further identified a fibrovascular motif at the tumor boundary that was linked to both risk of metastasis and failure of ICB. Those spatial and molecular biomarkers could be employed for the design of novel MCC therapeutics and personalized management strategies in MCC patients.

## 3. Discussion

Despite significant advances in uncovering MCC tumor biology, it remains poorly understood how immune-evasive mechanisms contextualize within the native TME of VP and VN-MCC. Further it is unknown how those relate to favorable patient outcomes for VP-MCC partly due to technological limitations to dissect the TME at single-cell resolution^4,6^.

Here we present a multi-modal framework including a combination of first-in-class multiplex imaging alongside laser capture micodissection-guided RNAseq of the MCC-TME. This framework informed our understanding of how MCPyV+ MCC re-organizes spatial behavior within the TME.

One striking observation from our study is the presence of a CD8^+^ dominant T cell infiltrate within the VP-MCC TME, but not the VN-MCC TME. These T cells were predominantly of an effector phenotype with elevated levels of Granzyme B and checkpoint molecules such as PD1, LAG3 or TIM3. These PD1^high^, LAG3^high^ CD8^+^ T cells were spatially enriched within tumor-centric CNs specifically of VP-MCC tumors. Given previous reports documenting that MCC-oncoprotein-specific T-cells exhibit a signature similar to exhausted CD8^+^ T cells^30-32^, our results indicate an enrichment of cancer-specific, cytotoxic CD8^+^ T cells inside the VP tumor compartment thus providing a possible link to favorable patient outcomes.

Second, we detected a significant enrichment of inflammatory CXCL9^+^, HLA-DR^+^ macrophages and DCs specifically at the tumor invasive front of VP-MCC. The enrichment of these inflammatory myeloid cells spatially coincided with an enhanced infiltration of CD8^+^ T-cells in the same spatial niches, a finding which we confirmed in HPV+ OPSCC, suggesting a common mechanism through with the immune system organizes anti-tumor immune responses in virus-associated cancers. Our receptor-ligand interaction analysis further indicates that specifically the interaction of CXCL9^+^ macrophages and DC with CXCR3^+^ T-cell at the tumor boundary is characteristic to VP-MCC and might drive the prognostically favorable infiltration of CD8^+^ T cells. This observation highlights the importance of appreciating cells not only in terms of simple phenotypic descriptors but also in terms of their functional characteristics^33^ to understand coordinated cellular behavior within the TME. In fact, inflammatory macrophages and DCs have recently been implicated in the recruitment and activation of CD8^+^ T cells^14,34-36^, suggesting a potential link to favorable survival outcomes found in VP-MCC - a finding that merits further investigations on the role of myeloid cells in re-shaping CD8^+^ T-cell-based anti-tumor responses in virus-associated solid malignancies^37^.

An urgent unmet need in the clinical management of local MCC is to identify patients who will develop distant metastasis and offer them an additional treatment strategy that would prolong DMFS. We here identified virus-positivity and high CD8^+^ T cell infiltration as predictors of prolonged metastasis-free survival^4^. Those biomarkers are easily applicable in a clinical setting and may inform clinical decision-making to identify patients at risk for distant metastasis. Given the promising results of the recent ADMEC-O interim-analysis on the efficacy of adjuvant nivolumab in local stage MCC^38^, our results suggest that particularly such patients at high-risk of distant metastasis should be considered for adjuvant systemic treatments such as ICB to prevent disease progression.

While ICB is currently not approved for use in local stage MCC, it is the only FDA-approved treatment for advanced MCC patients that significantly prolongs survival. However, many of these patients present either with primary (∼50%) or acquired resistance ^39,40^ emphasizing the need for actionable biomarkers to maximize clinical benefit. While prior observations revealed that MHC-class I expression within the TME, as well as the abundance of memory T cells and γδ-T-cells is associated with ICB-response in advanced MCC-patients^7,8,41-44^, less is known on how these features spatially contextualize within the MCC-TME and whether those markers might be applicable across VP and VN-MCC patients. Importantly, spatial approaches have shown that cells do not function independently within the TME but rather act in concert as multicellular units (CNs)^13,44^. Identifying characteristic interactions of such CNs, termed spatial contexts, in effective vs ineffective anti-tumor immune responses has been shown to be vital to reveal biological rules governing these interactions^33,45^. Through our enhanced spatial analysis, we identified unique spatial contexts linked to ICB-response, that included the intersection of T/B-cell enriched CNs within the MCC-TME. Our data thereby lay the foundation for future strategies aimed at enhancing the ICB-efficacy by fostering a supportive environment for structures like TLS through targeted therapeutic interventions. Furthermore, our observations strongly support that this spatial ICB-response signature might be applicable to both VP and VN-MCC patients alike, thereby offering a personalized treatment management approach.

Conversely, we detected that a spatial context defined by the enrichment of a CAFs at the tumor boundary associated both with distant metastasis and low response to ICB. This finding is in line with recent *in-vivo* data from a MCC xenograft model, in which mice injected with the MCC tumor cell MLK-1 in combination with human CAFs more often developed distant metastasis compared to mice injected with tumor cells alone^22^. The CAF-cancer cell interaction may therefore represent a novel therapy target that might complement adjuvant ICB or improve treatment outcomes. In agreement, prior reports documented that the combination of PD-L1 and TGF-ß targeting therapies to modulate CAF functionality, was able to reverse T-cell exclusion and re-instate an effective immune responses ^46-48^.

In summary, we have presented a multi-modal framework to dissect the native TME of two common virus-linked cancers at single-cell resolution, that provides a large (*8,000,000* cells), first-of-its-kind, and critically spatial publicly available dataset of MCC for the research community. Using this unique spatial dataset, we provide a comprehensive model of MCPyV-driven-immunomodulation and how it is linked to patient outcomes, thereby identifying a group of patients at high-risk for distant metastasis. We validated our model in HPV-linked OPSCC, suggesting common spatially organized mechanism of immunomodulation across viral cancers. Future work will require analysis of a larger independent series of MCC and deeper characterization of the proposed biomarkers for improved patient prognostication, as well as functional characterization of the CAF-tumor interaction to fully understand their role in metastasis development. Given the efficacy of ICB in MCC in both VP and VN MCC patients, adjuvant ICB^49,50^ represents a promising treatment strategy to prolong metastasis-free survival that might be particularly impactful in this high-risk group of local stage MCC patients^38^.

### Limitations of the study

This study has several limitations. While our study represents the first, and so-far largest exploration of virus-associated immunomodulation within the MCC-TME, it was single-institution with a modest patient cohort size and might therefore not represent the full diversity of Merkel cell carcinoma. It will therefore be imperative to validate our findings in larger, multi-institutional cohorts to confirm the suitability of the proposed biomarkers and tissue signatures for predicting distant metastasis and ICB-response. Our deep, multiplexed spatial analysis in combination with validation using an independent scRNAseq dataset and spatial profiling of a second cohort of virus-associated solid tumors (HNSCC) helps to mitigate this issue. Second, although our data suggest virus-specific modulation of T-cell responses between virus-positive and virus-negative MCC, we did not employ spatial transcriptomics or T-cell receptor (TCR) sequencing approaches, which could provide deeper insights into T-cell clonality, antigen specificity, and functional states *in situ*. Third, this study lacks in vivo models to directly assess the causal role of Merkel cell polyomavirus (MCPyV) in shaping immune responses within the TME. The future application of recently developed immunocompetent mouse models of MCC may help to address this mechanistic gap in the future^51^. Finally, the interpretation of highly multiplexed spatial datasets remains methodologically complex. To facilitate further discovery, we released all raw and processed data publicly available, including high-resolution images, segmentation masks, and single-cell datasets comprising over 10 million cells from human MCC and OPSCC (10.6019/S-BIAD1607). We anticipate that these datasets will spur the development of novel computational and AI-assisted tools in digital pathology.

## Acknowledgements

This work was supported by the US National Institutes of Health (P01HL108797,U01AI101984, 5U54CA209971, 5U01AI140498, U54HG010426, U19AI100627, 5P01AI131374, UH3DK114937, U19AI135976, U2CCA233238, U2CCA233195, U19AI057229, U54HG012723); the US Food and Drug Administration (HHSF223201610018C, DSTL/AGR/00980/01); Cancer Research UK (C27165/A29073); the Bill and Melinda Gates Foundation (OPP1113682); the Cancer Research Institute; the Parker Institute for Cancer Immunotherapy (PICI0025); Hope Realized Medical Foundation (209477); the Kenneth Rainin Foundation (2020-1463); the Beckman Center for Molecular and Genetic Medicine; Celgene (133826, 134073); Vaxart, Inc. (202627); and the Rachford Carlotta A. Harris Endowed Chair to G.P.N. M.H. is supported by the Deutsche Forschungsgemeinschaft (DFG, German Research Foundation) (project number: HA 9793/1-1). M.M. is supported by R01 CA229529 06. Y.T. is supported by a Stanford Dean’s Fellowship and the Stanford Cancer Institute Cancer Innovation Award. A.D.G. is supported by the Ovarian Cancer Research Alliance (Mentored Investigator Grant, MIG-2023-2-1015). We thank the Stanford Genomics Service Center for providing technical and analytical expertise. We also thank Angelica Trejo and Gustavo Vazquez (Stanford University) for excellent technical assistance. We are grateful for the patients who consented to use their tissues for research.

This article reflects the views of the authors and should not be construed as representing the views or policies of the FDA, NIH, BMGF, or other institutions that provided funding. Some figures were created with BioRender.com.

## Ethics declaration

The study was reviewed and approved by the Institute of Pathology, University Medical Center Mainz and the local ethics committee (Ethik-Komission der Landesärztekammer Rheinland-Pfalz, No: 2020–14822). Patients gave informed consent for contributing biomaterial and clinical metadata to the study before taking part. Analysis of the anonymized patient data was conducted in accordance with the guidelines of the declaration of Helsinki.

## Author information

### Author contributions

M.H. conceived and coordinated the study. M.H. M.M., and J.W.H. designed and performed experiments. M.H., Y.T., G.L.B. and M.M. analyzed and interpreted data, created figures. M.H., M.M., and Y.T., and wrote the manuscript. Y.T., M.M., G.L.B., M.H. and T.N.K. conceived and computationally implemented the conceptual framework. M.H., S.Z., P.C. and S.M. designed and created the TMAs. M.H., H.S., F.R. obtained clinical and pathological information. P.C., M.B. and A.D.G. helped with methodological validation. C.B., S.A. and M.G. generated MCC-xenograft data and helped analyze the data. Y.G. conceived CODEX, built the computational image processing pipeline, and provided technical advice. S.G., J.W.H. and G.P.N. supervised the study and wrote the manuscript. All authors revised the manuscript and approved its final version.

### Corresponding author

Correspondence to Garry P. Nolan

## Data Availability

A Single cell data table of the pre-processed and normalized MCC-CODEX imaging data, as well as combined CODEX and RNAscope imaging of the MCC-TME, in addition to the CODEX OPSCC dataset has been deposited at Mendeley data and will be publicly available as of the date of publication at: DOI: <u>10.17632/2m8fs89st7.1</u>.

LCMseq data have been deposited at GEO and will be publicly available as of the date of publication with accession number: GSE281260

Raw CODEX and RNAscope imaging data, alongside imaging masks will be publicly available at The BioImage Archive.

DOI: 10.6019/S-BIAD1607

Accession numbers for existing, public datasets are listed within the main text and include GSE226438.

Anonymized clinical metadata are reported in Supplemental Table 6.

Any additional information required to reanalyze the data reported in this paper is available from the lead contact upon request.

## Code Availability

Original code is deposited at https://github.com/nolanlab/Viral_TME_Reorganization_MCC/ and will be publicly available as of the date of publication.

## 4. Methods

### Study design and study cohort

A cohort of 242 patients who were diagnosed with primary MCC between August 2007 and August 2022 at the University Medical Center Mainz, Germany were screened for eligibility. Of these 242 patients, 60 patients with survival-follow-up until August 2023 were retrospectively identified according to the following selection criteria (**Supplemental Fig.1A**): histopathological confirmed diagnosis of MCC, sufficient pre-treatment FFPE tissue available for construction of tissue microarrays, DNA isolation, and laser-capture microdissection, follow-up longer than 3 months, and complete documentation of treatment outcomes. Clinical data on baseline patient and tumor characteristics, as well as treatment specifics, toxicity, the time and patterns of tumor recurrence, subsequent disease management (i.e., additional treatments following first-line treatment failure), and survival outcomes were collected. The study was reviewed and approved by the Institute of Pathology, University Medical Center Mainz and the local ethics committee (Ethik-Komission der Landesärztekammer Rheinland-Pfalz, No: 2020–14822). Patients gave informed consent for contributing biomaterial and clinical metadata to the study before taking part. Regional cutaneous, soft tissue and lymph node metastases were recorded as locoregional recurrences, all other as distant metastases. Primary clinical endpoint of this study was Distant-metastasis free survival (DMFS) for patients with initial local stage I/II disease. DMFS was calculated from primary diagnosis of MCC until the time of distant organ metastasis or death of any cause. Real-world tumor response as assessed by the investigators was categorized into complete response (CR), partial response (PR), stable disease (SD) and progressive disease (PD) as described earlier^52^.

Secondary endpoints included OS as defined as time from initial diagnosis of MCC until the last day of follow-up or death of any cause.

For DMFS analysis, we excluded patients who had no initial local stage I/II disease (n=26). Analysis of the anonymized patient data was conducted in accordance with the guidelines of the declaration of Helsinki.

### Construction of tissue microarrays

FFPE tissue blocks were retrieved from the tissue archives of the Institute of Pathology, University Medical Center Mainz, Germany and the Department of Dermatology, University Medical Center Mainz, Germany. The TMA contained 1-mm diameter cores from 51 malignant and semi-malignant samples and 15 non-malignant tissues samples; for details see **Supplemental Table 6**. Representative tumor and normal tissue regions were annotated on corresponding H&E-stained sections by a board-certified surgical pathologist (S.Z.). The TMA was sectioned at 3 µm thickness either onto SuperFrost Plus microscopy slides or square glass coverslips. Square glass coverslips (Electron Microscopy Sciences) were pre-treated with Vectabond (Vector Labs) according to the manufacturer’s instructions. Briefly, coverslips were immersed in 100% acetone for 5 min and then incubated in a mixture of 2.5 mL Vectabond and 125 mL 100% acetone in a glass beaker for 30 min. Coverslips were washed in 100% acetone for 30 s and air-dried, baked at 70 °C for 1 h, and stored at room temperature. For the MCC study, four independent 94-core TMAs, one 18-core TMA and one 9-core TMA were created, containing four 1.5mm cores per tumor block. TMA cores were annotated by S.Z., and M.H., under the supervision of B.W.B., as follows: For each tumor block, two representative areas at the tumor invasive front and two representative areas from the tumor core were annotated. TMAs were sectioned at 3 μm thickness and mounted onto SuperFrost Plus microscopy slides (CODEX analysis) or UV-activated membrane-coated slides for subsequent laser-capture microdissection. The 3-μm thick sections of the TMAs were stored at 4°C in a vacuum desiccator (Thermo Fisher) containing Drierite desiccant (Thermo Fisher) until analysis.

### Generation of CODEX DNA-conjugated antibodies

Buffers and solutions for CODEX antibody conjugation and multiplex imaging were prepared as previously described ^53^. For CODEX antibody conjugation we used a previously described protocol^33^. Maleimide-modified short DNA oligonucleotides were purchased from Biomers.net. Conjugations were performed at a 2:1 weight/weight ratio of oligonucleotide to antibody with at least 50 μg of antibody per reaction. All centrifugation steps were at 12,000 g for 8 min, unless otherwise specified. Purified, carrier-free antibodies (for details on clones and manufacturers see **Supplemental Table 6**) were concentrated on 50-kDa filters, and sulfhydryl groups were activated using a mixture of 2.5 mM TCEP and 2.5 mM EDTA in PBS, pH 7.0, for 30 min at room temperature. After washing the antibody with buffer C, activated oligonucleotides were resuspended in buffer C containing NaCl at a final concentration of 400 mM. Oligonucleotides were then added to the antibody and incubated for 2 h at room temperature. The conjugated antibody was washed by resuspending and spinning down three times in PBS containing 900 mM NaCl. It was then eluted by centrifugation at 3,000 g for 2 min in PBS-based antibody stabilizer (Thermo Fisher) containing 0.5 M NaCl, 5 mM EDTA, and 0.02% w/v NaN_3_ (Sigma), and stored at 4 °C.

### Manual and robotic IHC

The antibodies used for CODEX were cross-validated and optimized by manual DAB-IHC (**Data Fig.S1**). All validation was performed under the supervision of a board-certified pathologist (S.Z.) and collated with known expression patterns published online (The Human Protein Atlas, Pathology Outlines) as well as the published literature.

Chromogen-based manual IHC was carried out according to a procedure described previously ^25^. Briefly, sections were cut to 4 μm thickness and placed on frosted histology glass slides (Thermo Fisher). Deparaffinization, rehydration, and heat-induced epitope retrieval were performed on an ST4020 small linear stainer (Leica). Activity of endogenous peroxidase was blocked via incubation with 3% H_2_O_2_ in PBS for 5 min. After subsequent washing steps, slides were incubated for 20 min with 2.5% normal horse serum to block non-specific binding. Antibodies were diluted in antibody diluent (DAKO) and sections were stained overnight in a sealed humidity chamber at 4 °C on a shaker. After staining, slides were washed for 10 min in 1x TBS IHC wash buffer with Tween 20 (Abcam), and specimens were covered with an HRP-conjugated polymer detection reagent (VectorLabs), followed by washing for 10 min. Bound antibodies were then visualized using the HRP/liquid DAB+ substrate chromogen system (Vector Labs) according to the manufacturer’s instructions. Sections were counterstained with hematoxylin, followed by dehydration, mounting, and imaging in brightfield mode on a BZ-X710 inverted fluorescence microscope (Keyence) with subsequent whole-slide scanning on an APERIO AT2 brightfield scanner.

In order to allow for a reproducible workflow and streamlined process, we established an automatic protocol on the Omni-Stainer platform (Parhelia Biosciences) using the same reagents as applied above. Briefly, following deparaffinization and heat-retrieval steps, slides were loaded into the Omni-Stainer and a 96-well-plate was prepared with the IHC antibodies, chromogens, and blocking reagents. Staining buffers were were added into separate wells of a liquid storage container. Detailed protocols for IHC and CODEX uniplex stains can be found on Stainworks platform.

In addition to antibody validation, we used chromogenic IHC to assess the expression of MCPyV-LT using clone Ab3 at a concentration of 1:100 and MHC class I expression using clone EMR8-5 at a concentration of 1:20.000 (for details see **Data Fig.S1**). Expression levels were assessed within the tumor compartment. MCPyV-LT expression was categorized as positive or negative, whereas MHC-I expression was determined on a semi-quantitative scale (low, medium, high). Evaluation of expression levels was assessed by M.H. prior to CODEX multicycle experiments and independently confirmed by a board-certified dermatopathologist (B.W.B.) who was blinded to clinical metadata and CODEX information.

### CODEX antibody screening, validation and titration

Conjugated CODEX antibodies were tested and titrated in low-plex fluorescence assays and resulting expression patterns, as well as signal-to-noise-ratio were cross-verified with manual IHC results (see **Data Fig.S1**). Antibody-oligonucleotide conjugates were tested together in a single CODEX multicycle and optimal dilution, exposure time and appropriate imaging cycle was determined for each conjugate (**Supplemental Table 6** specifies CODEX multi-cycle reaction and image acquisition details and **Data Fig.S2-4** shows staining results for CODEX multicycle imaging validated in a multi-Tumor TMA and MCC). All validation was performed under the supervision of a pathologist (S.Z.) and confirmed with online databases (The Human Protein Atlas, Pathology Outlines) and the published literature.

### CODEX FFPE tissue staining and fixation

Coverslips and or microscope slides were handled using Dumont coverslip forceps (Fine Science Tools). For deparaffinization, coverslips were baked at 70 °C for at least 1 h, followed by immersion in fresh xylene for 30 min. Sections were rehydrated in descending concentrations of ethanol (100% twice, 95% twice, 80%, 70%, doubly distilled H_2_O (ddH_2_O) twice; each step for 3 min). Heat-induced epitope retrieval was performed in a Lab Vision PT module (Thermo Fisher) using target retrieval solution of pH 9 (DAKO) or pH 6 (Akoya Biosciences) at 97 °C for 10 min. After cooling to room temperature for 30 min, coverslips were washed for 10 min in 1x TBS IHC wash buffer with Tween 20 (Cell Marque). Non-specific binding was blocked for 1 h at room temperature using 100 μL of blocking buffer [S2 buffer containing B1 (1:20), B2 (1:20), B3 (1:20), and BC4 (1:15)] as described previously ^33^. For each coverslip, DNA-conjugated antibodies were added to 50 μL of blocking buffer on a 50-kDa filter unit, concentrated by spinning at 12,000 g for 8 min, and resuspended in blocking buffer to a final volume of 100 μl. This antibody cocktail was then added to the coverslip and staining was performed in a sealed humidity chamber overnight at 4 °C on a shaker. After staining, coverslips were washed for 4 min in S2 and fixed in S4 containing 1.6% paraformaldehyde for 10 min, followed by three washes in PBS. Then, coverslips were incubated in 100% methanol on ice for 5 min, followed by three washes in PBS. Fresh BS3 fixative was prepared immediately before final fixation by thawing and diluting one 15 μL aliquot of BS3 in 1 mL PBS. Final fixation was performed at room temperature for 20 min, followed by three washes in PBS. Thereafter, coverslips or microscope slides were stored in S4 at 4°C for up to two weeks, or further processed for imaging.

### CODEX multi-cycle reaction and image acquisition

We used coverslips for validation of antibodies in low-plex and multicycle reactions. Following CODEX staining coverslips were removed from storage buffer S4, rinsed in ddH_2_O to remove salt residues and mounted onto custom-made CODEX acrylic plates (Bayview Plastic Solutions) using coverslip mounting gaskets (Qintay), creating a well in the acrylic plate above the tissue section for liquid storage and exchange. Tissue stained with Hoechst nuclear stain at a dilution of 1:600 in H2 buffer for 1 min, followed by three washes with H2 buffer. The CODEX acrylic plate was mounted onto a custom-designed plate holder and secured onto the stage of a BZ-X710 inverted fluorescence microscope. Fluorescent oligonucleotides (concentration: 200 nM) were aliquoted into Corning black 96-well plates in 250 μL H2 buffer containing Hoechst nuclear stain (1:600) and 0.5 mg/ml sheared salmon sperm DNA. Black plates were sealed with aluminum sealing film (VWR Scientific) and kept at room temperature during the multi-cycle reaction.

All final multicycle panels were performed on SUPERFROST Plus microscopy slides with a total imaging area of 18 x 35 mm. Following staining, slides were removed from S4, brought to room temperature in PBS, and rinsed in ddH2O to remove salt residues. Meanwhile a Flow cell (Akoya Biosciences Cat. 240205) was prepared so that the pressure sensitive adhesive side of the flow cell faced up in the flow cell assembly device. Subsequently, slides were thoroughly dried, except for a thin PBS film on the tissue. The slide was inverted and aligned with the flow cell. To ensure proper seal, the flow cell and slide were then aligned for 30 s using slight pressure from the flow cell assembly device (Akoya Biosciences). Upon proper sealing, the slide with the attached flow cell were stored in H2 buffer for 10 min. Prior to assembly onto the PhenocyclerFusion, slides were cleaned with 70% ethanol to remove any residual debris. Similar to the CODEX multi-cycle reaction with the coverslips, fluorescent oligonucleotides were aliquoted into Corning black 96-well plates in 235 µl H2 buffer and 0.5 mg/ml sheared salmon sperm DNA. DAPI (ThermoFisher Scientific, D1306) was used at a concentration of 165 µg/ml.

Automated image acquisition and fluidics exchange were performed using an Akoya PhenocyclerFusion instrument with Akoya Biosciences driver software. Briefly, this entailed cyclic stripping, annealing, and imaging of fluorescently labeled oligonucleotides complementary to the oligonucleotide conjugated to the antibody. Metadata from each CODEX run can be found in **Supplemental Table 6**. For multi-cycle imaging of the TMA spots, the “fine-core TMA function” was programmed to the center of each TMA spot, and an automatic focusing point was set for each imaging cycle. After each multi-cycle reaction, the flow cell was removed. Following pre-treatment in xylene, H&E staining was performed according to standard pathology procedures, and tissues were scanned in brightfield mode. For the MCC TMAs that used 56 different antibodies and DAPI as nuclear staining, the entire multicycle experiment took about 56 hours to complete per run and resulted in a total of 104.5 x 10^6^ single-cell protein readouts per TMA (30,000 cells per 1.5-mm diameter core). To ensure consistency and reproducibility of the results across multiple runs we followed previously described workflows ^33,54-57^. As such, we stained all 6 MCC-TMAs at the same time using the same antibody and imaging specifications across all TMAs. As imaging was limited to one sample at the time, the remaining samples were stored meanwhile in storage buffer at 4°C. Staining quality, marker expression, and distribution were verified on each individual TMA.

### CODEX post-imaging data processing

Raw imaging data were processed using the RAPID uploader for image stitching, drift compensation, deconvolution, and cycle concatenation^58^. Processed data were then segmented using the CellVisionSegmenter, a neural network R-CNN-based single-cell segmentation algorithm ^59^. Both the RAPIDS image processing software and Segmenter software can be downloaded from our GitHub site (https://github.com/nolanlab/CODEX), and the CellVisionSegmenter software can be accessed at https://github.com/michaellee1/CellSeg. After the upload, images were evaluated for specific signal. Any markers that produced an untenable pattern or a low signal-to-noise ratio were excluded from the ensuing analysis. Uploaded images were visualized in ImageJ (https://imagej.nih.gov/ij/).

### CODEX imaging data analysis

### Cell-type labels and identification

Cell type identification was done following the methods developed previously ^53,55^. Briefly, nucleated cells were selected by gating DAPI-positive cells, followed by z-normalization of protein markers for each TMA core and removal of cells with high normalized protein expression values in more than 90% of the detected proteins. The data was over-clustered with a Leiden-based clustering approach using the *scanpy* Python package. Cell types were assigned to each cluster based on average cluster protein expression and location within image. Clusters were mapped onto the tissue for manual verification of the assigned clusters. Impure clusters were split or re-clustered following mapping back to original fluorescent images. Following identification of major cell types, cell subtypes were identified by re-clustering the corresponding major cell type clusters for the phenotype-specific protein markers (i.e., CD45RO, CD45RA, PD-1, LAG-3, Granzyme B, CXCR3, CCL5, EOMES for T cell memory subtypes).

Neighborhood analysis was performed as described previously ^33^. Briefly a window size of 20 nearest neighbors was taken across the tissue cell-type maps. Cells of each type were quantified for each of these windows, and then these vectors were clustered into commonly composed neighborhoods. We employed k-means elbow plot and overclustering to inform our choice for the number of clusters of cellular neighborhoods. For overclustering, we generated 20-30 clusters and mapped the clusters back to the tissue and evaluated for cell-type enrichments to determine overall structure.

Cell distance matrices were calculated using squidpy^60^. Briefly, cells were detected within each field of view using the x, y position. Cell distances to an anchor cell (terminally exhausted CD8^+^ T cells) were identified for all cells that were more abundant than 10 cells / tissue sample Cell distances were finally compared using Bonferroni-adjusted t-test with FDR <0.05.

Investigators (M.H., Y.T.) were blinded to clinical metadata information during the single-cell spatial data analysis avoid classification bias.

### Differential enrichment analyses

Linear models *Y_n,c_* = β_0_ + β_1_*X* + β_3_*Y*_c_ + e were estimated, where *Y*_c_ is the log overall frequency of cell type c, *X* is an indicator variable for patient group, *Y_n,c_* is the log frequency of cell type c in CN n, β_i_ are coefficients, and e is mean zero Gaussian noise. A pseudo-count of 1e^-3^ was added prior to taking logs. These were estimated using the *statsmodels* Python package ^61^. The coefficient estimates and p-values for β_1_ were extracted and visualized.

Receptor-Ligand proximity analysis:

To analyze receptor-ligand interactions, we first identified ligand positive cells (e.g., PD-L1^+^ cells) and receptor positive cells (e.g., PD-1^+^ cells) based on their expression levels. PD-L1^+^ cells were then used as reference points to find their k-nearest neighbors (k = 10). Considering that PD1-PDL1 interactions require direct contact, we implemented a distance cut-off of 20 µm based on prior literature ^14^. This analysis was conducted across all TMA regions. For each TMA region, we quantified valid receptor-ligand interactions by retaining only those ligand-receptor pairs that met the distance criterion. The resulting valid pairs were then totaled for each TMA region to provide a measure of receptor-ligand interactions under different experimental conditions.

### Spatial context maps

Spatial context maps were created as previously described ^13^. First, we used a large window size (100 nearest neighbors) for each cell across the tissue neighborhood labels in order to create composition vectors. Second, instead of clustering the vectors of neighbor compositions, we determined the combination that included the fewest neighborhoods that made up more than 85% of the cells within that window. This combination informs about prominent associations of neighborhoods in the window, a feature we term spatial context. Third, we counted each combination and connected the most prevalent combinations together into a spatial context map.

### Barycentric coordinate projection

The windows computed for the spatial context maps were projected under the linear map sending the unit coordinate vectors to a point of an equilateral triangle^56^. The projected points were colored by the CN assignment at the center of the window. Barycentric density plots were colored by gaussian density. The frequencies of the CNs assigned to cells corresponding to different vertices, edges, and inter-mixing of three CNs mapped to a barycentric plot were computed and normalized by the total number of cells assigned to the corresponding three CNs per TME core. To determine significance, t-tests corrected for multiple hypothesis testing (Bonferroni) were calculated between the normalized CN-interface frequencies and the clinical outcome parameters of the same TMA core across all tissue samples.

### Cell-cell interaction analysis

Pairwise cell-cell interactions were calculated as described previously. Briefly, cell co-occurrences within 32 pixels were computed on a global scale for all unique regions. and then stratified by condition. The frequency of co-occurences was highlighted as edges and nodes as the average cell frequency within the investigated patient cohort. Results were plotted as a circlos plot^62^. In addition we used scimap to identify significant cellular interactions within tissue contexts by comparing observed adjacency frequencies to a random distribution^63^.

### DNA/RNA isolation from FFPE tumor blocks and PCR readout

Genomic DNA was extracted using the AllPrep DNA/RNA FFPE Kit (Qiagen, 80234) per manufacturer’s instructions. The MCPyV-status of each sample was initially screened for in conventional IHC using MCPyV large T-antigen antibody Ab3 followed by validation using PCR and qRT-PCR with primers listed in **Supplemental Table 6** as described previously^12^ (**Supplemental Fig.2C-F**). PCR was carried out in triplicate with a final volume of 25 µl using a PCT-200 (MJ-Research) under the following conditions for LT3 and LT1: 94 °C for 5 min, followed by 35 cycles of 94 °C for 30 s, 53.5 °C for 30 s, 68 °C for 30 s, then final 68 °C for 10 min, followed by electrophoresis on 2% agarose gel and visualized under ultraviolet light. Quantitative PCR was carried out in triplicate in a 384-well plate format using ABI 7900HT Real-Time PCR Detection System (Applied Biosystems) with a pre-treatment at 50 °C for 2 min, followed by denaturation at 95 °C for 10 min, and then by 45 cycles of 95 °C for 15 s and 60 °C for 1 min. The RT-PCR reaction had a final volume of 20 µl consisting of DNA (10 ng), primers and probe (5 µmol/L each), nuclease-free water, and 10 µl reaction buffer (LuminoCt Read Mix, Sigma Aldrich).

### Laser-capture microdissection

Two serial sections of the tissue microarray were taken at 7 µm thickness and mounted onto frame slides with polyethylene naphthalate membranes (Thermo Fisher Scientific, #LCM0521). Slides were immersed for 20 s each in xylene (three times), 100% ethanol (three times), 95% ethanol (two times), 70% ethanol (two times), water, hematoxylin (Dako, S3309), water, bluing reagent (Thermo Fisher Scientific, 7301), water, 70% ethanol (two times), 95% ethanol (two times), 100% ethanol (three times), and xylene (three times). Immediately after staining, cells were dissected from every tissue microarray spot on an ArcturusXT LCM System (Thermo Fisher Scientific) using the ultraviolet laser to cut out the desired region and the infrared laser to adhere the membrane to a CapSure HS LCM Cap (Thermo Fisher Scientific, LCM0215). A tissue area containing roughly 15,000 mononuclear cells was captured from each spot, with cell numbers determined based on density estimates by cell counting in an adjacent H&E-stained section. If a core had more than 1000 mononuclear cells, a tissue fragment containing around 1000 mononuclear cells was dissected from that core. If a core had less than 1000 cells, tissue fragments from corresponding cores on the serial section membrane were combined in the same LCM cap to obtain approximately 1000 cells. After microdissection, the caps were sealed using 0.5-ml tubes (Thermo Fisher Scientific, N8010611) and stored at −80 °C until cDNA library preparation.

### Preparation of cDNA libraries and RNA sequencing

Sequencing libraries were prepared according to the Smart-3Seq protocol for LCM HS caps as previously described ^54^. Briefly, 10 µl of lysis mix consisting of 40% (v/v) 5 M trimethylglycine solution (Sigma, B0300), 20% (v/v) 10 mM nuclease-free dNTP mix (Thermo Fisher Scientific, R0192), 10% (v/v) 20 µM first-strand primer in TE buffer (1 S, 5Biosg/GT GAC TGG AGT TCA GAC GTG TGC TCT TCC GAT CTT TTT TTT TTT TTT TTT TTT TTT TTT TTT TV; Integrated DNA Technologies), 10% (v/v) Triton-X 100 (Sigma #T8787; diluted to 0.5% v/v in molecular biology-grade water), and 20% (v/v) Proteinase K (New England Biolabs, P8107S; diluted to 0.125 mg/ml in in molecular biology-grade water) was added to the center of each LCM cap. Caps were sealed with 0.2 ml low-retention PCR tubes (Corning, PCR-02-L-C) and incubated on a pre-warmed metal CapSure incubation block (Thermo Fisher Scientific, LCM0505) at 60 °C in an incubator. Then, tubes were briefly centrifuged, and 10 µl of template-switching reverse-transcription (TS-RT) FFPE LCM mix consisting of 40% (v/v) 5x SMARTScribe first-strand reaction buffer (Clontech, 639537), 20% (v/v) 20 mM DTT (Clontech, 639537), 10% (v/v) 20x RNase inhibitor (Thermo Fisher Scientific, AM2694), 4% (v/v) 50 µM second-strand primer in TE buffer (2 S, /5Biosg/CT ACA CGA CGC TCT TCC GAT CTN NNN NrGrG rG; Integrated DNA Technologies), 4% (v/v) 200 mM MgCl_2_ (Sigma, 63069), 2% 5 mM proteinase K inhibitor (EMD Millipore, 539470), and 20% (v/v) 100 U/µl SMARTScribe reverse transcriptase (Clontech, 639537) was added. Samples were incubated in a programmable thermal cycler (42 °C for 30 min, 70 °C for 10 min, 4 °C hold), and 1.25 µl of a unique P5 primer and 1.25 µl of a universal P7 primer (2 µM in TE buffer each; Integrated DNA Technologies; sequences available upon request) and HiFi HotStart ReadyMix (Kapa, #KK2601) were then added, followed by 22 cycles of PCR amplification (98 °C for 45 s; 22 cycles at 98 °C for 15 s, 60 °C for 30 s, 72 °C for 10 s; then 72 °C for 60 s, and 4 °C hold). Amplified cDNA was purified with SPRI bead mix (Beckman Coulter, B23317) and a magnetic separation block (V&P Scientific, VP772F4). Finally, the samples were washed with 80% ethanol and resuspended in TE buffer to yield the sequencing-ready library. Libraries were profiled for size distribution on an Agilent 2200 TapeStation with High Sensitivity D1000 reagent kits and quantified by qPCR with a dual-labeled probe. The RNA libraries were sequenced on an Illumina NextSeq 500 instrument with a High Output v2.5 reagent kit (Illumina, #20024906) to a minimum sequencing depth of 1.5 M reads per sample.

### Processing of RNA-seq data

Base calls from the NextSeq were de-multiplexed and converted to FASTQ format with bcl2fastq (NextSeq system suite software, Illumina v2.20.0.422). The five-base unique molecular identifier (UMI) sequence and the G-overhang were extracted from FASTQ data, and A-tails were removed with umi_homopolymer.py (github.com/jwfoley/3SEQtools). Reads were aligned and further processed to remove duplicates using STAR v2.7.3a (github.com/alexdobin/STAR)^64^. Bulk gene expression profiles were transcript per million (TPM) normalized and log2 transformed.

### Principal component analysis (PCA) immune scores

PCA scores and PC1 coefficients were computed for the normalized bulk RNA-seq data on a per spot basis using the prcomp function in base R. The IFN-γ score was calculated using the six gene signature published by Ayers et al.^65^. The TGF-β score was calculated using the 15 gene signature published by Mariathasan et al.^46^. The immune activation and immunosuppression scores were computed using the genes listed in **Supplemental Table 6**. Differences in PC1 scores between patient groups were modeled using the Wilcoxon test.

### CIBERSORTx deconvolution

The LM22 signature matrix was used to deconvolve mixed gene expression profiles using the HiRes deconvolution mode using R scripts downloaded from the CSx website (arguments used: rmbatchSmode = T, QN = F)^66^. Log2 fold changes were computed for every deconvolved gene across patient groups. Differences in gene expression between patient groups were computed using the Wilcoxon test. The *p* values were adjusted with the Benjamini-Hochberg correction using the p.adjust function in R.

### scRNA-seq data processing

The publicly available scRNA-seq dataset used for external validation in our study is available in the NCBI Gene Expression Omnibus database under the accession code GSE226438 and was previously published to identify transcriptomic correlates of epithelial-mesenchymal transition in MCC ^12^. We obtained preprocessed scRNA-seq count data from this dataset. Data were filtered for cells containing a minimum of 1,000 and no more than 10,000 unique genes and containing less than 10% mitochondrial genes using the python package *scanpy*^67^ as described in ^68^. Doublets were identified using *scrublet*^69^ as counts with a predicted doublet score >0.2 and excluded from subsequent analysis Additionally, genes detected in fewer than 4 cells were also removed prior to data normalization. For the ‘treatment-naive’ dataset, the nine sample libraries were anchored and integrated using the top 2,000 variable features identified using *scVI*. The integrated object was log-normalized, and PCA was performed. Thereafter, the first 50 PCs were used for UMAP construction and non-linear dimensionality reduction followed by calculating the k-nearest neighbor graph and leiden clustering at varying resolution (0.3-1.0). The marker genes per cluster were identified using the rank_gene_groups function based on Wilcoxon Rank Sum test. Heatmaps and dotplots were created for the cell-type defining genes based on scaled expression values (**Data Fig.S5**). Marker gene expression was used to annotate cell-type specific clusters, and further sub-clustering was done in case of more granular phenotyping (i.e., subclassification of CD4^+^ T cell subsets and CD8^+^ T cell subsets). To compare treatment naïve and ICB-treated patients, we integrated 11 samples (9 treatment-naive and 2 ICB-treated tumors), followed by dimensionality reduction and cluster annotation results. Differential gene expression analysis was conducted using a Wald test from the *pydeseq2* and *diffxpy* packages^70^. Pseudo-bulk transcriptomes were generated by summing the UMI counts across cells as recommended for *DESeq2*, and mean expression values between the investigated conditions were compared using Wilcoxon-rank-sum test corrected for multiple hypothesis testing.

### Generation and Analysis of combined CODEX and RNAscope multiplex imaging

To validate the enrichment of CXCL9 and CXCL10-positive myeloid cells at the tumor invasive front we established a protocol of combined multiplex RNAscope imaging with CODEX multiplex imaging. Therefore, a 3µm thick section of the MCC-TMA was cut onto a SuperFrost PLUS slide and processed according to RNAscope^TM^ Multiplex Fluorescent Reagent Kit v2 (UM3223100). In brief, slides were baked for 1h at 63°C, followed by immersion in fresh xylene for 2x5min. Sections were washed in 100% ethanol for 2min before being air-dried for 5min on an adsorbent paper tissue. Prior to antigen retrieval using pH9 solution, the tissue was incubated for 10min with RNAscope Hydrogen peroxide and washed 3 times with ddH2O. Staining of the tissue with RNA probes CXCL9 (#1154158-C3) and CXCL10 (# 553198-C1) was carried out using a HybEZ oven set to 40°C according to manual protocol UM3223100. Probes were detected using OPAL dyes OPAL570 (1:120) and OPAL780 (1:25). Following staining and detection of RNA probes we continued with staining of CODEX antibody-conjugates (for panel information **see Supplemental Table 6**), followed by flow cell mounting and CODEX multiplex imaging using the Phenocycler Fusion imager.

Raw RNAscope and CODEX multiplex imaging data were registered, which allowed for the detection of both RNA signal and protein signal with spatial sub-cellular resolution. Processed data were then segmented using the CellVisionSegmenter, a neural network R-CNN-based single-cell segmentation algorithm ^59^. Segmentation resulted in a dataset encompassing 1,347,684 cells. Cells were annotated with support of an AI-agent (cfteammate), that was trained on scripts previously generated for the annotation of the CODEX-only MCC-dataset. The annotated dataset was analyzed similar as described above (see CODEX imaging data).

### Construction of the OPSCC TMA, and subsequent generation and analysis of CODEX multiplex imaging data

A cohort of 476 patients who were diagnosed with primary HNSCC between 2005 and 2019 at the University Medical Center Mainz, Germany, were screened. Eighty-six patients with survival follow-up data until April 2022 were retrospectively identified according to the following criteria: Histopathological confirmed diagnosis of oropharyngeal squamous cell carcinoma (OPSCC), the absence of secondary HNSCC tumors diagnosed within the observation period, sufficient pre-treatment tumor tissue available for the generation of a tissue microarray (TMA), >30% of uncompromised tumor tissue on TMA for each patient, treatment with primary radiotherapy or chemoradiotherapy (RTx/RCTx), complete follow-up documentation of treatment outcomes. Details on clinical-pathological parameters at initial diagnosis, subsequent treatments and survival data were collected from the patient’s medical records and have been described previously^25^.

Of the entire cohort of 476 patients, all 86 patients who met the selection criteria and for whom one or more paraffin tissue blocks were available were included in the present study. Formalin-fixed paraffin-embedded tissue blocks of these 86 OPSCC patients were retrieved from the archives of the Institute of Pathology. The area of malignancy was marked by a board-certified pathologist (S.Z.). The tissue microarray was constructed from 1,2mm diameter cores that were punched from a representative region of the tumor tissue blocks according to standard procedures.

The TMA was sectioned at 3µm thickness onto SuperFrostPLUS slides and processed following the procedures described above (Generation of CODEX multiplex imaging data). We established a 56-marker antibody panel for this OPSCC cohort and stained the TMA similar as described above. The stained TMA was imaged using the PhenocyclerFusion. Details on the panel used and on multicycle details are specified in **supplemental Table 6**. Raw CODEX multiplex imaging data were stitched, and background subtracted following the internal PCF protocols. We used MESMER – a pre-trained deep learning algorithm for automated, accurate cell segmentation^71^ – for the initial segmentation and marker quantification. Cell-type annotation and subsequent downstream analysis was performed as described before with support of the python package spacec^10^.

### Murine MCC-xenograft experiments and CODEX multiplex imaging

MCC-xenograft experiments were previously performed as described in detail in ^22^. In brief, CAFs were directly isolated from the MCC tissue of 9 MCC patients. For the in-vivo tumorigenic assay female severe combined immunodeficient mice were used (CB17/lcr-Prkdc scid/lcrlcoCrl; Charles River Laboratories, Wilmington, MA). HFFs (Human foreskin fibroblasts, code: SCRC-1041, ATCC) or CAFs from Patients were injected with MLK-1(MCPyV-positive MCC cell line, code: 09111801, Sigma-Aldrich) as described^22^. Tumors were collected, formalin-fixed and paraffin-embedded.

Tumor blocks were assembled as Tumor macroarrays with 5 samples of 3µm thickness each being sectioned onto a total of 2 SuperFrostPLUS slides.

Macro-TMAs were processed following the procedures described above (Generation of CODEX multiplex imaging data). We established a 20-marker antibody panel for this MCC-xenograft experiment including both mouse-specific and human-specific antibodies (**Supplemental Fig.10L**) as described above. The stained TMA was imaged using the PhenocyclerFusion. Details on the panel used and on multicycle details are specified in **supplemental Table 6**. Raw CODEX multiplex imaging data were stitched, and background subtracted following the internal PCF protocols. CAF phenotypes and spatial location were subsequently correlated with tumor kinetics and in-vivo endpoints as described in ^22^ using the open-source software QuPath.

### Clinical data analysis and multimodal data integration

Descriptive statistics were used to analyze the baseline characteristics of the study population. A chi-square test was used to assess the association between the compared groups (i.e., VP-MCC vs. VN-MCC; ICB responder vs. non-responder). The Clopper–Pearson method was used to calculate 95% confidence intervals for the categorical variables. As time-to-event endpoints, this retrospective cohort study used DMFS and OS, which were estimated using the Kaplan–Meier product-limit method and log-rank statistics using the *survminer* package in R. The median duration of follow-up was calculated using the reverse Kaplan–Meier method. Independent prognostic values of clinical patientś characteristics were estimated using univariate and multivariate Cox proportional hazard models. Multivariate analysis was calculated for the significant variables by the univariate test or *a priori* selection for biological relevance to evaluate their conjoint, independent effects on OS. In all cases, two-tailed p-values were calculated and considered significant with value of *p* < 0.05.

Integration of clinical metadata and molecular CODEX data for exploratory biomarker analyses was done in the Python package *ehrapy*^18^. Briefly, clinical and molecular data were combined into a single anndata object and subjected to initial quality checks and data normalization. For the clinical outcome variable (here: Progress to metastatic disease), we selected all clinical and molecular variables that were available at the baseline of the corresponding clinical outcome (i.e., at initial diagnosis) for subsequent dimensionality reduction. We then used a Leiden-based clustering approach to determine the proximity of the inputted variables and determined highly variable features of each group using t-tests corrected for multiple hypothesis testing. Those features were subsequently further explored using established approaches such as Cox-regression-analysis and Kaplan-Meier analysis to validate their impact on the investigated clinical outcome.

Lasso was used to identify markers associated with time-to-event clinical endpoints. To construct a Lasso model, we used the *glmnet* implementation of Lasso in R, including a built-in cross-validation function to tune the L1 regularization parameter λ. We used the minimum mean-cross validated error ^19^. Briefly, for each investigated endpoint (DMFS) we included all CODEX variables and those clinical variables that were found to be significantly associated with the corresponding endpoints variable into our regression model. Here, we obtained lambda.min and the model coefficients via the cross-validation function in *glmnet* as previously described to identify those parameters that were significantly associated with the investigated clinical endpoints ^72^. For all regularized Cox models, we only included patients for whom baseline tissue biopsies were available (i.e., biopsies from initial diagnosis; DMFS, n=26).

### Statistical analysis

Statistical analyses were performed with RStudio (version 2023.06.1) and Python (version 3.8.11). Results with *p* < 0.05 were considered significant unless otherwise stated. No statistical methods were used to predetermine sample size. For differences across the tested groups, including comparisons of VP vs VN MCC, patients with vs without distant metastasis, significance was tested using two-tailed t-test corrected for multiple hypothesis testing (Bonferroni). If Boxplots were used for comparing continuous variables between VP and VN-MCC tumors, all boxplots were structured as follows: Center line represents median of all MCC-tumor samples in that group. Box-limits denote interquartile range (1.-3. quartile), upper whiskers denote limits of Q3+1.5xIQR and lower whiskers denote limits of Q1-1.5xIQR. All boxplots show all individual datapoints from a given MCC-tumor sample that have been calculated as the average across all investigated tumor cores within that tumor sample (i.e. average from 3-4 tumor cores for each tumor sample). Tests involving comparisons between more than two groups were done using Kruskal-Wallis (across group). Tests involving comparison of groups with categorical values were performed using two-sided Fisher’s exact tests. All statistical tests were two-sided unless otherwise specified. Correlations were reported by the Spearman rank correlation and reported with Benjamini-Hochberg adjusted p-values. Hazard ratios and results from log-rank test for survival analysis were calculated using the survival package in R. For differential gene expression between patient groups LCM-seq data sets, the significance was tested using linear mixed-effect models, where *p* values were adjusted using Benjamini-Hochberg correction.

\**p* < 0.05, \*\**p* < 0.01, \*\*\**p* < 0.001, \*\*\*\**p* < 0.0001.

