## Supplemental Figs.1-14 for "Spatially organized inflammatory myeloid-CD8^+^ T cell aggregates linked to Merkel-cell Polyomavirus driven Reorganization of the Tumor Microenvironment"

[1] Department of Microbiology and Immunology, Stanford University School of Medicine, 94305, CA, USA
[2] Department of Pathology, Stanford University School of Medicine, 94305, CA, USA
[3] Department of Dermatology, University Medical Center Mainz, 55131, Germany
[4] Institute of Pathology, University Medical Center Mainz, 55131, Germany
[5] Molecular Biosciences/Cancer Biology Program, Heidelberg University, Heidelberg, 48232, Germany
[6] German Cancer Research Center, DKFZ, Heidelberg, 48232, Germany
[7] Department of Cancer Immunology and Virology, Dana Farber Cancer Institute, Boston, 02215, MA, USA
[8] Department of Hematology, University Medical Center Heidelberg, Heidelberg, 48232, Germany
[9] Institute of Molecular Virology and Tumor Virology, University Medical Center Hamburg-Eppendorf, Hamburg, Germany
[10] Virology Unit, Department of Translational Medicine, University of Piemonte Orientale, Novara, Italy
[11] Department of Biomedical Engineering, Duke University, 27708, NC, USA
[12] Lead contact
[13] These authors contributed equally

**Supplemental Fig.1: MCC study cohort and investigated tumor samples. Related to Fig.1.**

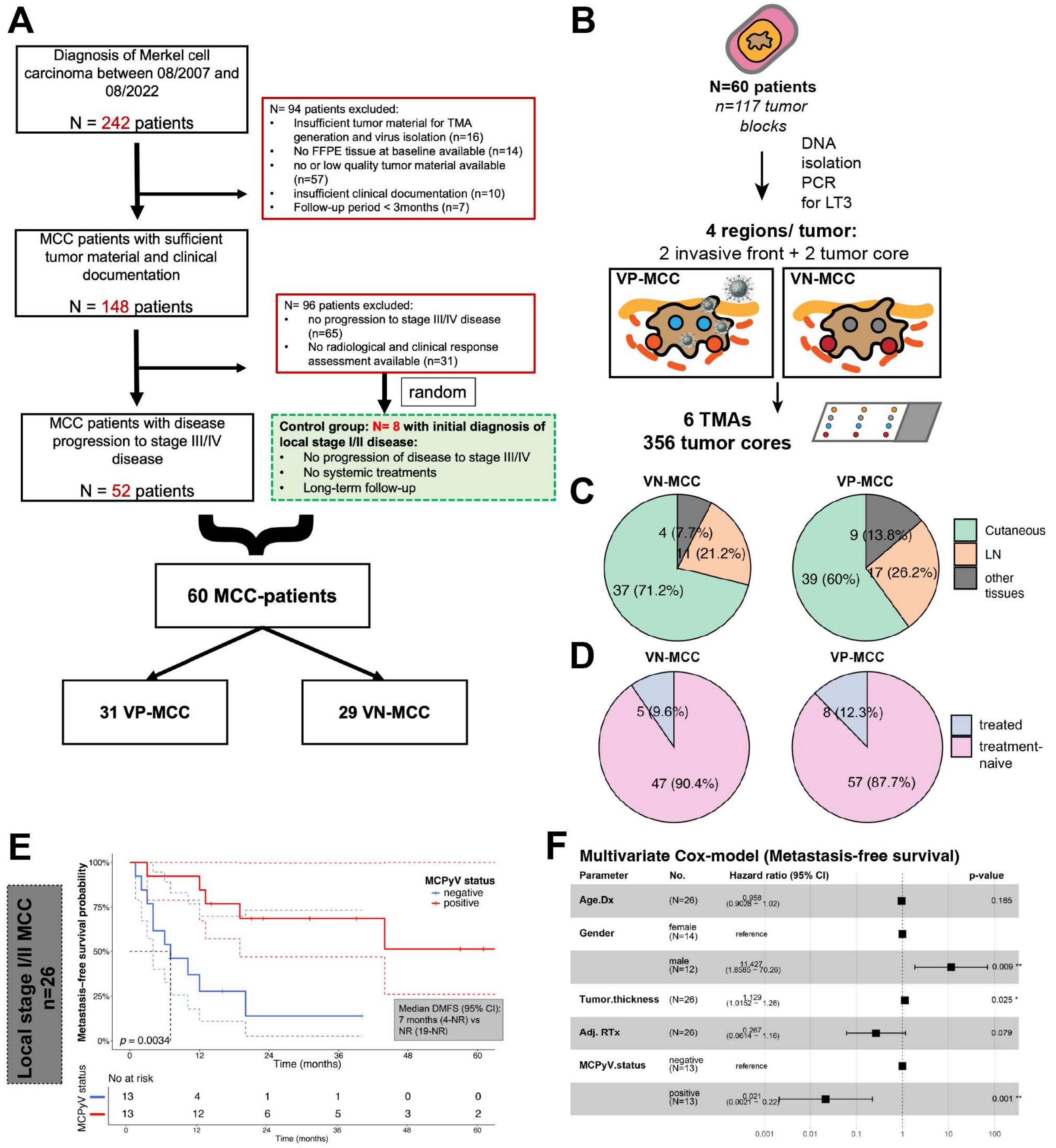


**(A)** CONSORT diagram depicting selection criteria for identification of the investigated MCC cohort. Exclusion criteria were insufficient tumor material for generation of TMA and viral isolation, no FFPE tissue available at baseline, low-quality tumor material available, insufficient clinical documentation, follow-up period < 3 months, unknown radiographic and clinical response assessment. **(B)** For MCC TMA assembly, longitudinally collected tissue samples were selected from 60 patients (step 1), DNA was isolated from FFPE blocks and viral status was determined using PCR and RT-PCR (step 2), and TMAs were assembled using four regions from each tissue block, two from the invasive tumor front and two from the tumor core (step 3). **(C)** Pie chart summarizing the relative distribution of tissue types within the investigated cohort stratified by viral status. **(D)** Pie chart summarizing the proportions of samples that were collected prior or after initiation of systemic treatments within the investigated cohort stratified by viral status. **(E)** Kaplan-Meier plot depicting DMFS probability in MCPyV-positive and MCPyV-negative patients with local stage I/II MCC who received standard-of care treatment. Statistical significance was determined using log-rank test. Median survival times indicated by black dotted line. 95% confidence intervals depicted as color-coded dotted lines. **(F)** Multivariate Cox-regression analysis for DMFS in patients with local stage I/II disease. Statistically significant results with $p<0.05$ are highlighted in bold. Related to Supplemental Tables 3-4.

**Supplemental Fig.2: ROI selection and identification of MCPyV status. Related to Fig.1.**

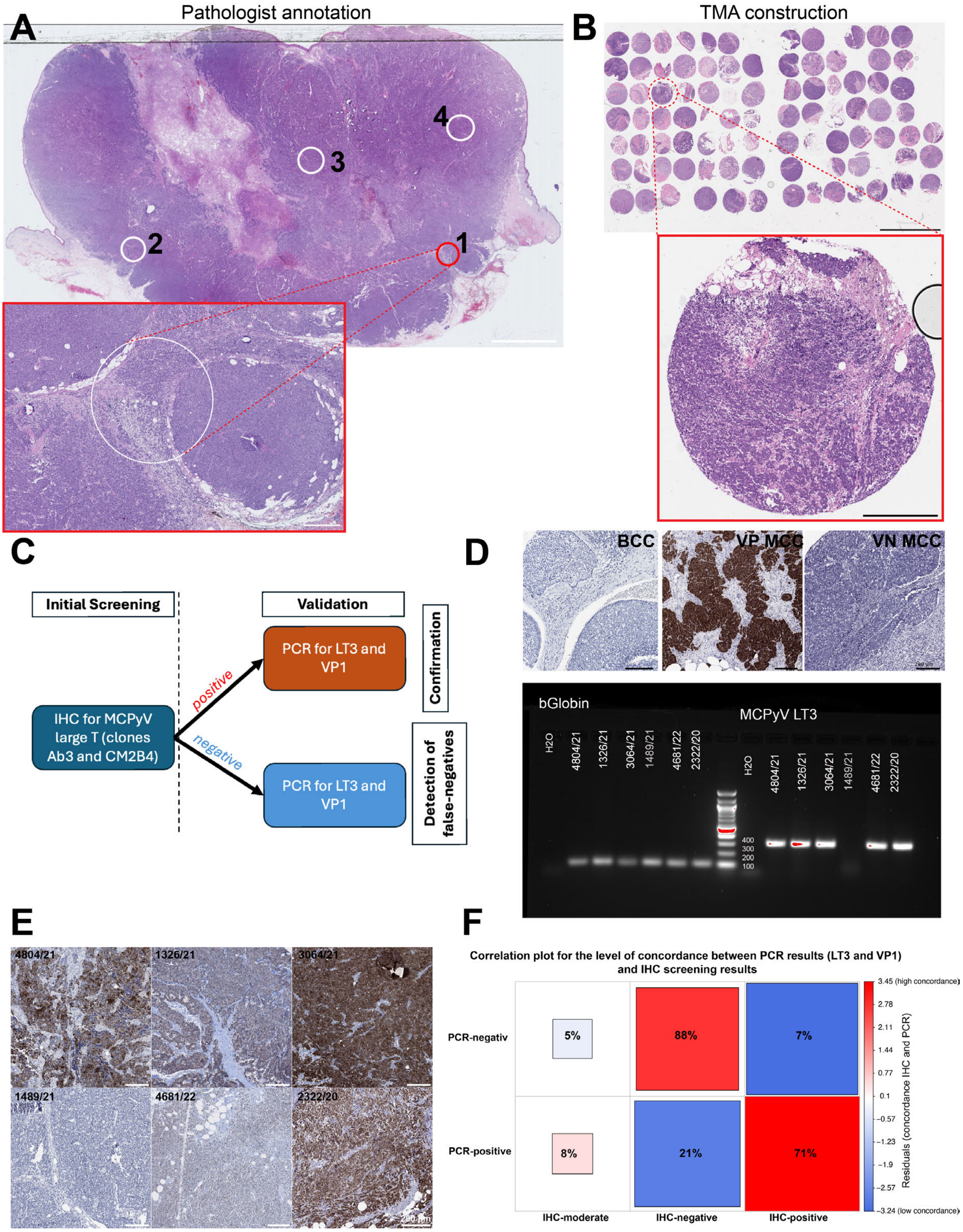


**(A)** Representative example of an MCC tissue sample showcasing the workflow for selecting cores to be included in the study. Hematoxylin and eosin (H&E) slides were digitized and subseqently annotated by a board-certified pathologist with demarcated with areas of interest (white circles). Scale bars, 3mm (top) and 500µm (bottom) apply. Each identified region was punched with a 1.5mm core and placed onto a tissue microarray (TMA). **(B)** Representative H&E stained tissue microarray of the MCC-cohort including the core punched from magnified region (1) shown in (A). Scale bars, 3mm (top) and 500µm apply. **(C)** Schematic of the workflow employed for determination of MCPyV status. Initial screening included IHC for MCPyV large T-antigen using clones Ab3 and CM2B4. Results were confirmed using PCR for LT3 and VP1 (see METHODS). **(D)** Representative results for initial IHC screening of MCPyV large T antigen using control basal cell carcinoma tissue (top, left), a VP-MCC sample (top, center) and VN-MCC sample (top, right). Representative results from PCR testing for MCPyV LT3 probe for the indicated samples (bottom). **(E)** Representative regions from MCC tumor samples stained for MCPyV large T-antigen clone CM2B4 in IHC corresponding to the samples tested for MCPyV LT3 in (D**)**. **(F)** Confusion matrix illustrating the correlation between results from IHC for MCPyV LT antigen and PCR using LT3 and VP1 primer pairs. Pearson residuals were calculated using chi-square test.

### Supplemental Fig.3: CODEX antibody validation. Related to Fig.1.

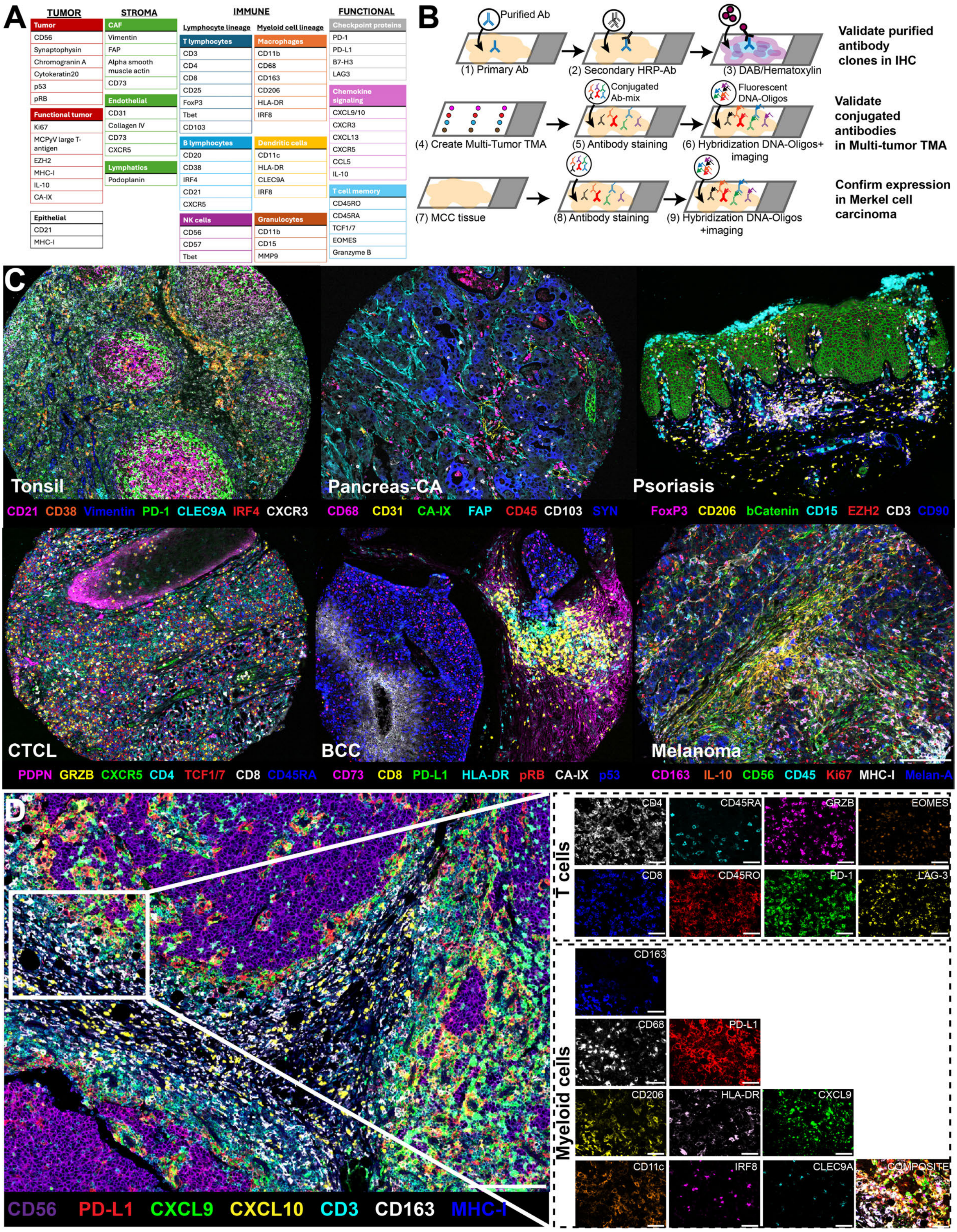


**(A)** CODEX antibody panel. **(B)** Experimental approach for validation of DNA-barcoded antibodies. **(C)** Images of selected cores from the multi-tumor TMA. Scale bar: 200 µm. Related to Data Fig.S1-4. **(D)** Left: Image of the tumor invasive front from selected tumor cores with markers for the T cell and myeloid cell compartments highlighted (left). Scale bar: 200 µm. Right: Images showing markers used for cell type identification in Leiden-based clustering (i.e., CD4 and CD8 in the T cell compartment and CD68, CD163, CD206, and CD11c in the myeloid cell compartment) and to identify functional subtypes (i.e., CD45RA and CD45RO for T cells and PD-L1 or HLA-DR for myeloid cells). Scale bars: 50 µm. *Abbreviations: SYN, Synaptophysin; GRZB, Granzyme B; PDPN, Podoplanin*.

**Supplemental Fig.4: Spatial distribution of CD8$^+$ T cells within the VP-MCC TME and transcriptional T-cell signatures characteristic for VP-MCC revealed by scRNAseq. Related to Fig.2.**

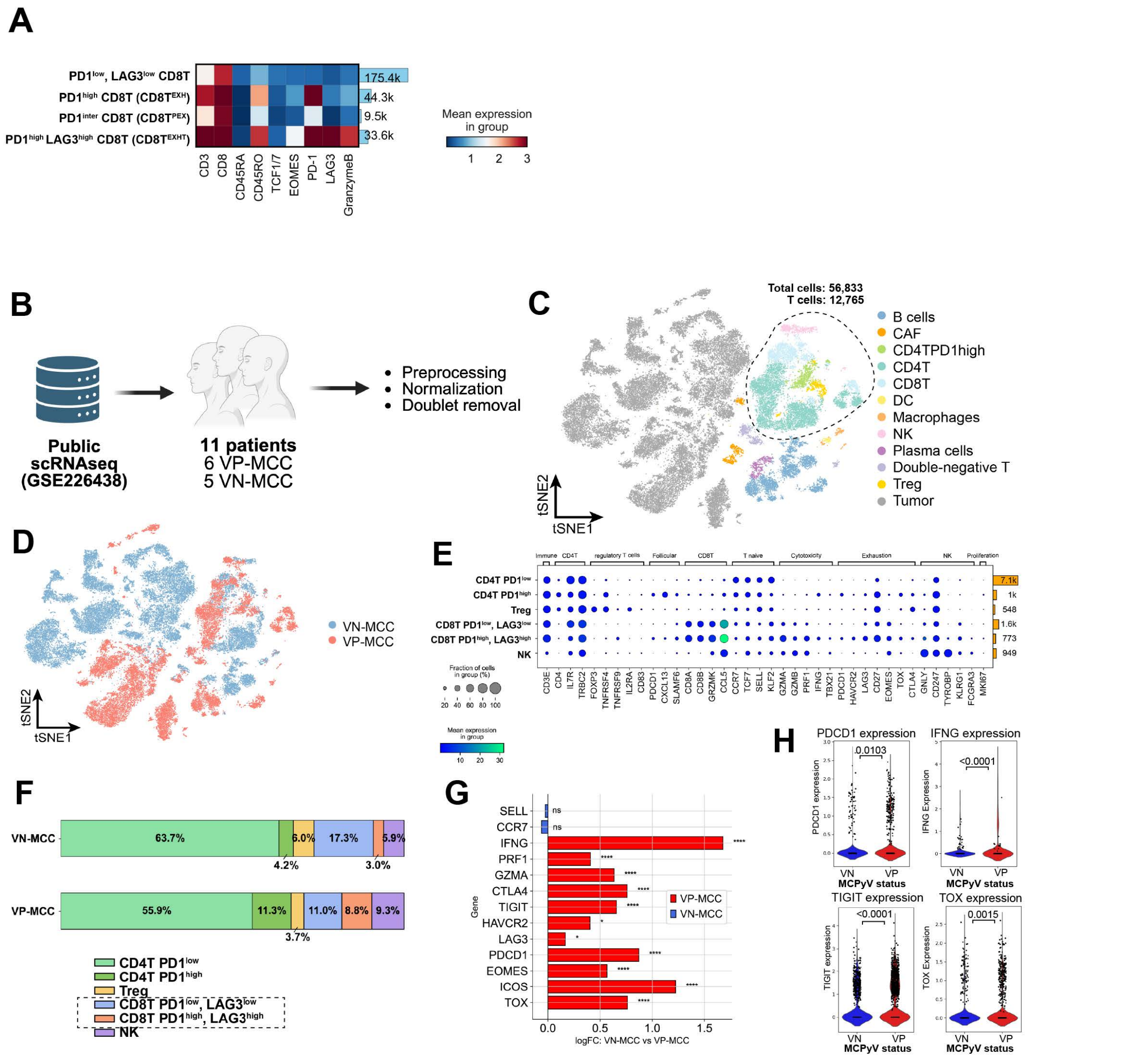

**(A)** Heatmap depicting T cell subsets defined by their expression of checkpoint molecules PD1 and LAG3, their absolute counts within the CODEX dataset and mean log-normalized expression of key phenotypic markers. **(B)** Analytical framework used to dissect the MCC-TME using scRNAseq (GSE226438). **(C)** tSNE projection of annotated cell types found in the scRNA-seq dataset. Related to Figure S4. **(D)** tSNE projection of all annotated cells found in the scRNAseq dataset colored by viral status. **(E)** Dotplot summarzing mean expression values amongst all T-cell subtypes with corresponding total counts in the scRNAseq dataset. **(F)** Stacked barplots comparing the relative abundances of T-cell subsets between VN and VP-MCC patients. CD8+ T cell subsets marked with dotted lines. **(G)** Waterfall plot depicting genes significantly enriched in VP vs VN-MCC samples amongst all CD8+ T -cells. Significance was determined using Wald-test from diffxpy package in python and was not corrected for multiple hypothesis testing. **(H)** Violinplots comparing mean expression of selected genes implicated in T cell activation and dysfunction shown as two-group comparison boxplots. Statistical significance was determined using Bonferroni-corrected Wilcoxon-rank sum test with FDR<0.05. *Abbreviations: log2FC, log(2) fold change; FDR, false-discovery rate.*

**Supplemental Fig.5: Identification of CNs within the TME stratified by viral status and tissue site, reveals CNs shared across MCC. Related to Fig.3.**

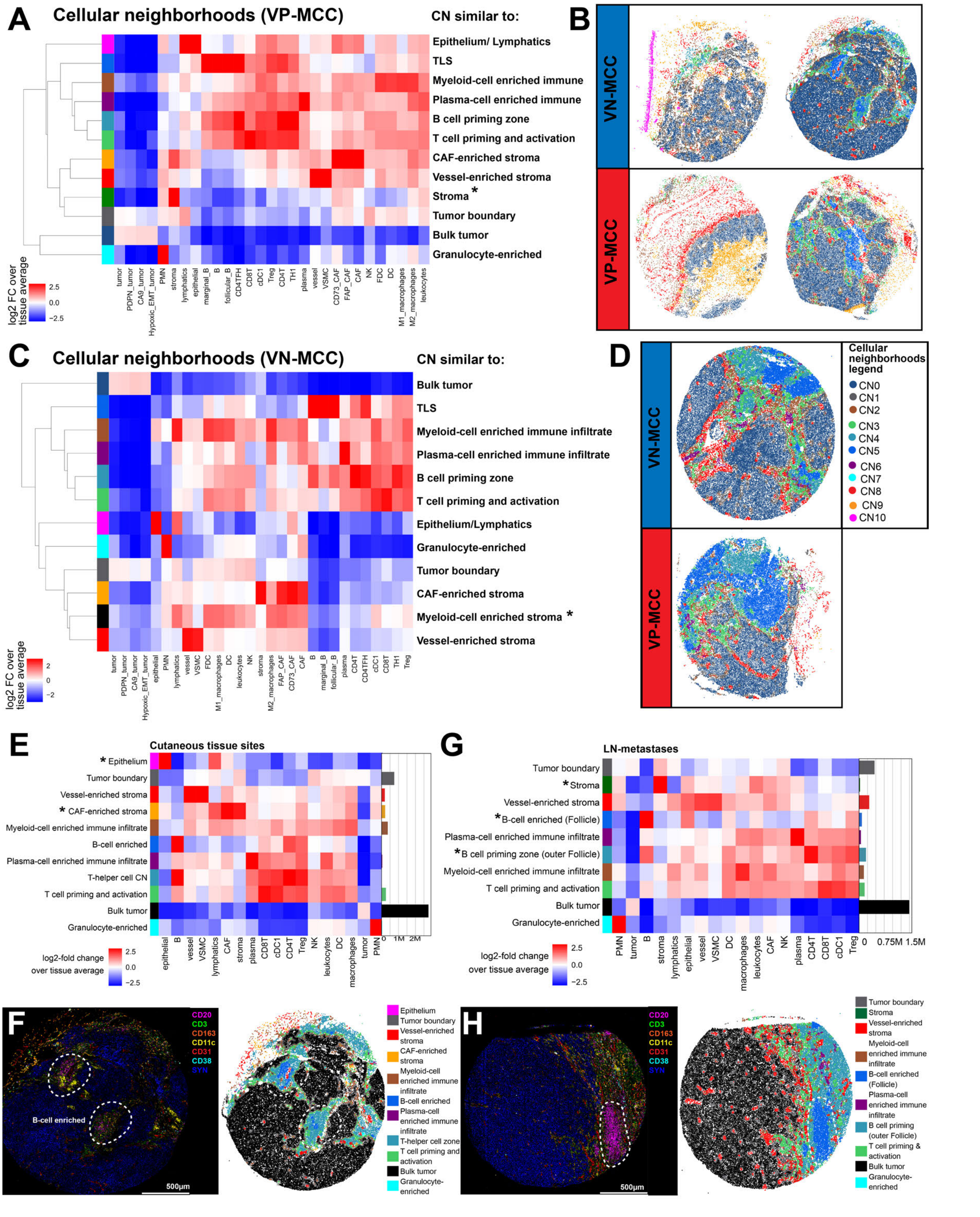

**(A)** CNs and their cell-type enrichment that were observed when clustering VP-MCC samples separately from VN-MCC. CNs different to those defined in Fig.3A are highlighted by Asterisks. **(B)** Representative images of VP-MCC and VN-MCC cutaneous tumors color coded by CN. Color codes from Supplemental Fig.5D apply. **(C)** CNs and their cell-type enrichment that were observed when clustering VN-MCC samples separately from VP-MCC. CNs different to those defined in Fig.3A are highlighted by Asterisks. **(D)** Representative images of VP-MCC and VN-MCC lymph node metastases color coded by CN. Color codes from Supplemental Fig.5D apply. **(E)** Heatmap depicting the log2-fold change in cell-type enrichment against the tissue average within each of the identified CNs in cutaneous tissue samples. CNs unique to cutaneous tissue sites are marked by asterixis. **(F)** Representative example of a cutaneous MCC-metastasis in both multiplex IF imaging (left) and corresponding CN geography map (right) with B-cell enriched CNs marked by dotted lines. Scale bars, 500µm. **(G)** Heatmap depicting the log2-fold change in cell-type enrichment against the tissue average within each of the identified CNs in LN metastases. CNs unique to LN metastases are marked by asterixis. **(H)** Representative example of an MCC-LN-metastasis in both multiplex IF imaging (left) and corresponding CN geography map (right) with the B-cell enriched germinal center CN marked by dotted lines. Scale bars, 500µm.

**Supplemental Fig.6: Spatial distribution of CD8$^+$ T cells and immunomodulatory molecules within the MCC-TME revealed by RNAscope and CODEX multiplex imaging. Related to Fig.3.**

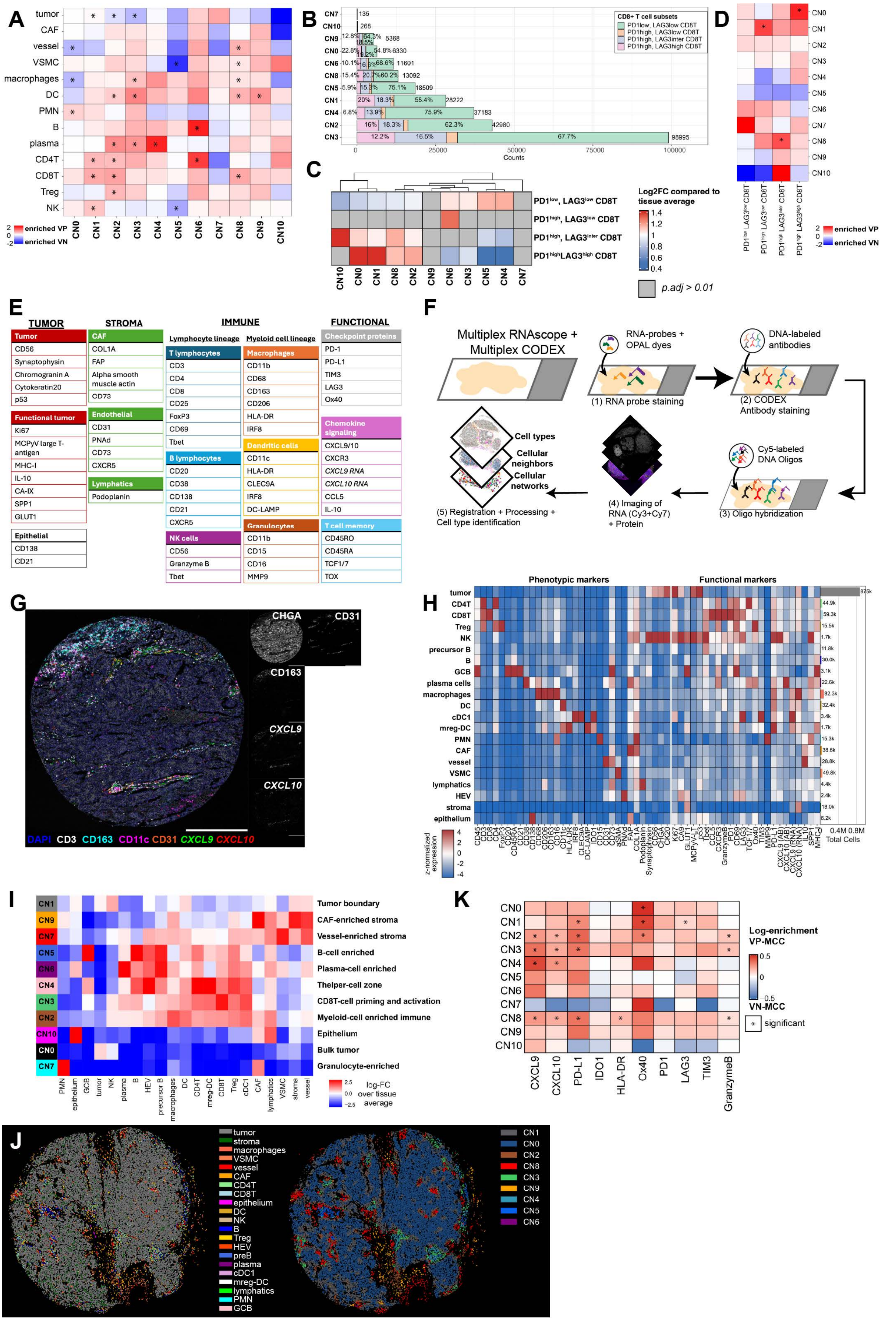

**(A)** Heatmap illustrating the enrichment of selected cell-types within each CN compared to the average global distribution of cells within the MCC CODEX data set. Asterixis indicate a significant enrichment of a given cell-type within this CN as computed through the statsmodel package in python (see METHODS). **(B)** Stacked barplot summarizing total counts for CD8 T cell subsets and their relative abundance across all CNs. Relative abundances below 5% are not printed. Total CD8 T cell counts per CN provided to the right of each stacked barplot. **(C)** Heat Heatmap depicting the relative enrichment of CD8+ T cell subsets across all CNs compared to the global tissue average of CD8+ T cell subsets within the entire MCC cohort. Statistical significance was determined using binomial testing corrected for multiple-hypothesis testing (Bonferroni). **(D)** Heatmap illustrating the enrichment of T cell dysfunction subtypes within each CN compared to the average global distribution of T cell dysfunction subtypes within the MCC CODEX data set. Asterixis indicate a significant enrichment of a given cell-type within this CN as computed through the statsmodel package in python (see METHODS). **(E)** Summary for the marker panel used for combined multiplex RNAscope and CODEX imaging. **(F)** Schematic illustrating the stepwise approach of combined multiplex RNAscope and CODEX multiplex imaging. **(G)** Representative tumor core from combined CODEX+multiplex RNAscope imaging with corresponding greyscale images of selected markers. Scale bar, 500µm. **(H)** Heatmap illustrating log-normalized mean expression of phenotypic (left) and functional markers across the identified cell types of combined CODEX and RNAscope multiplex imaging with corresponding cell counts. **(I)** Heatmap with CNs identified through combined CODEX and RNAscope multiplex imaging. CNs were identified similar as described in the CODEX multiplex imaging dataset using windows for kNN =20 cells. **(J)** Representative cell-type geography and corresponding CN geography maps derived from combined multiplex CODEX and RNAscope imaging. **(K)** Heatmap depicting the log2FC enrichment of a given marker across all identified CNs between VP and VN-MCC with red color indicating an enrichment in VP-MCCs and blue color indicating an enrichment in VN-MCC. Significant results are indicated by asterixis. Statistical significance was determined using Wilcoxon-rank sum test corrected for multiple-hypothesis testing (BH).

**Supplemental Fig.7: Phenotypic characteristics and spatial interactions of macrophages and dendritic cells within the MCC-TME. Related to Fig.4.**

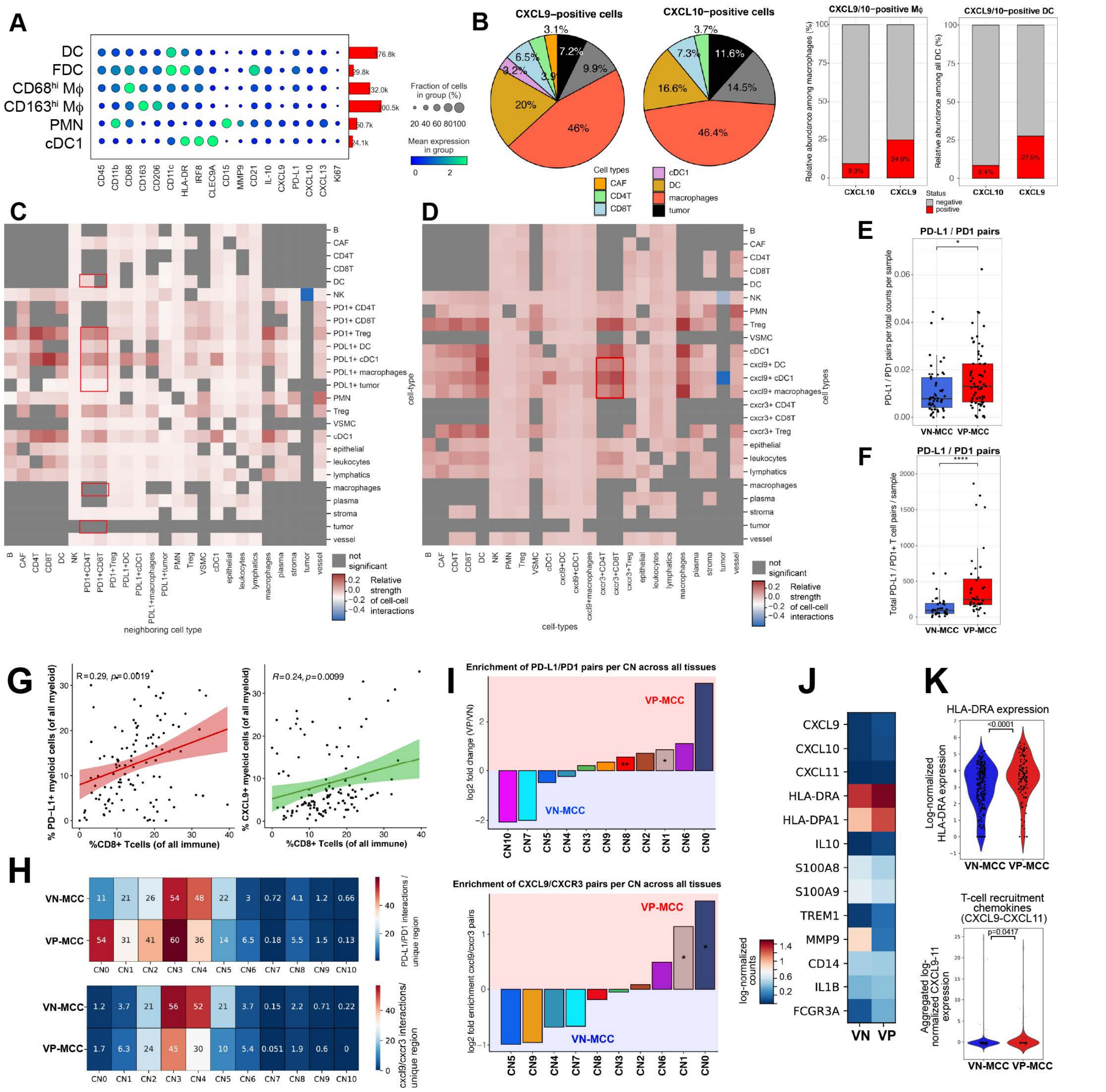


**(A)** Dotplot summarizing log-normalized mean marker expression of phenotypical and functional markers within the identified myeloid cell subtypes and their absolute cell type counts. **(B)** Piecharts (left) illustrating cell-types expressing the immunomodulatory CXCL9 and CXCL10 chemokines as detected in combined multiplex RNAscope and CODEX multiplex imaging and stacked barcharts (right) for CXCL9/CXCL10 expression in macrophages and DC. **(C)** Heatmap summarizing results from cell-cell-interaction analysis through random permutations[72], where interactions significantly enriched in VP-MCC are shown in red, non-significant interactions are colored in grey and cell-cell interactions found predominantly in VN-MCCs are marked in blue. Myeloid-cell types were annotated based on PD-L1 expression, while T-cell subsets were annotated based on PD1 positivity. **(D)** Heatmap summarizing results from cell-cell-interaction analysis through random permutations[72], where interactions significantly enriched in VP-MCC are shown in red, non-significant interactions are colored in grey and cell-cell interactions found predominantly in VN-MCCs are marked in blue. Myeloid-cell types were annotated based on CXCL9 expression, while T-cell subsets were annotated based on CXCR3 positivity. **(E)** Boxplots comparing the relative abundance of PD-L1/PD1 pairs in the myeloid/Tcell compartment normalized to the total number of cells within each investigated tumor core. Statistical significance was determined using Wilcoxon-rank sum test adjusted for multiple hypothesis testing (BH). **(F)** Boxplots comparing the mean abundance of PD-L1/PD1 pairs in the myeloid/Tcell compartment within each investigated MCC sample of cutaneous origin (primary tumors or cutaneous metastases). Statistical significance was determined using Wilcoxon-rank sum test adjusted for multiple hypothesis testing (BH). **(G)** Scatterplot highlighting correlations found between PD-L1$^{+}$ and CXCL9$^{+}$ myeloid cells and CD8$^{+}$ T cells within each investigated tumor sample as assessed using Spearman correlation analysis. **(H)** Heatmaps summarizing the enrichment of PD-L1/PD1 pairs (top) and CXCL9/CXCR3 pairs (bottom) between all CNs within the CODEX dataset. **(I)** Waterfall plots summarizing the log-fold enrichment of PD-L1/PD1 pairs within all CNs between VP and VN-MCC tumors amongst all investigated tumor samples (top) or log-fold enrichment of CXCL9/CXCR3 pairs within all CNs between VP and VN-MCC tumors. Statistical significance was determined using Wilcoxon-rank sum test adjusted for multiple hypothesis testing (BH). $*p < 0.05$, $**p < 0.01$. **(J)** Matrixplot derived from the scRNAseq dataset showing the enrichment of genes reflective of an inflammatory macrophage signature stratified by viral status. **(K)** Violinplots illustrating differences in mean expression levels from HLA-DR (top) and inflammatory chemokines CXCL9, CXCL10 and CXCL11 within the macrophages identified from scRNAseq. Statistical significance was determined using Bonferroni-corrected Wilcoxon-rank sum test with FDR<0.05.

**Supplemental Fig.8: CODEX antibody validation. Related to Fig.4.**

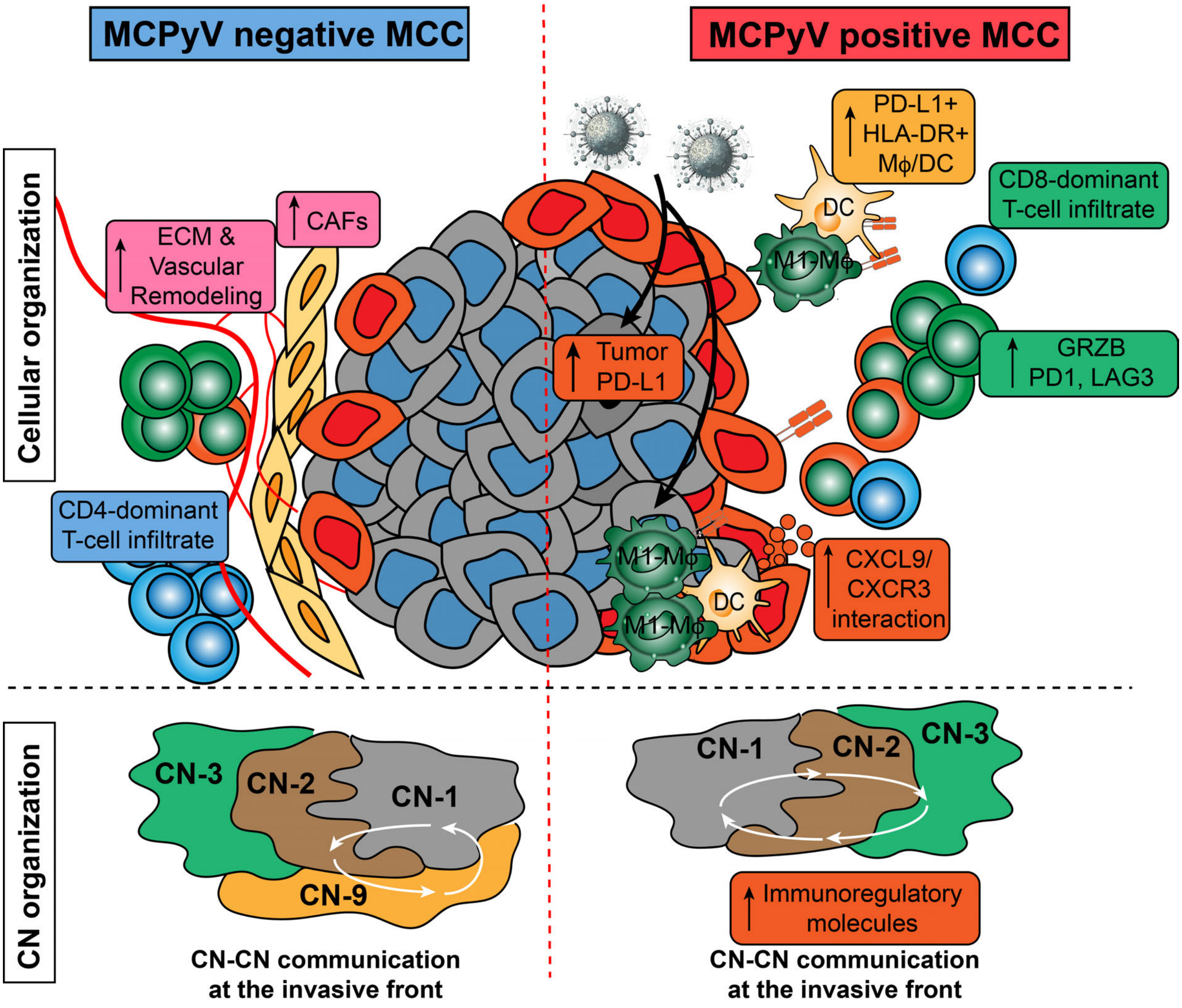


Cartoon of the proposed model for cellular (top), architecture (bottom), and mediator-driven differences in TMEs of VP-MCC and VN-MCC.

**Supplemental Fig.9: Skin-specific validation of cellular and spatial features characteristic to the VP-MCC-TME. Related to Fig.2-4.**

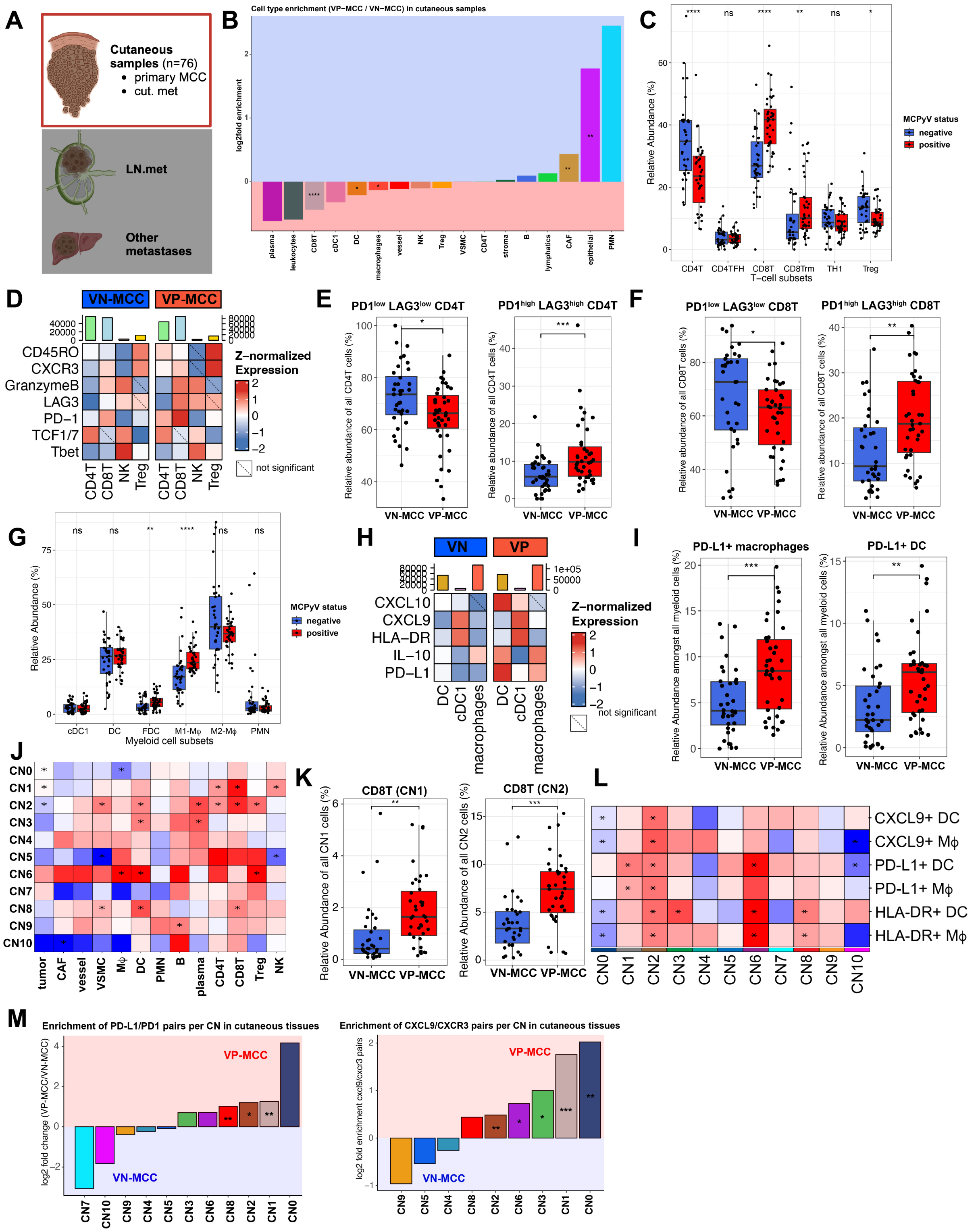


**(A)** Schematic summarizing the tissue samples investigated in this subset analysis, that included only tissues from cutaneous sites including primary tumors and cutaneous metastases. **(B)** Waterfall plot summarizing the log2-fold enrichment of cell-types outside of the tumor compartment between VP and VN-MCC cutaneous samples. Statistical significance was determined using Wilcoxon-rank sum test adjusted for multiple-hypothesis testing (BH). $*p < 0.05$, $**p < 0.01$, ***$p$<0.005, ****$p$<0.001. **(C)** Grouped boxplots comparing the relative abundance of T cell subtypes between VP and VN-MCC samples relative to the total T-cell compartment. Statistical significance was determined using Wilcoxon-rank sum test corrected for multiple-hypothesis testing. $*p < 0.05$, $**p < 0.01$, ***$p$<0.005, ****$p$<0.001. **(D)** Heatmaps comparing the mean z-normalized expression of selected markers within cells of the T-cell compartment between VP and VN-MCC samples. Non-significant differences are marked by dotted lines. Statistical significance was determined using Wilcoxon-rank sum test adjusted for multiple-hypothesis testing (BH). **(E)** Relative abundance of checkpoint-protein positive and negative CD4$^{+}$ T cells states normalized to the total number of T cells in VP-MCC vs VN-MCC. Each point represents the mean cell frequency per MCC-sample. Significance was determined using Bonferroni-adjusted t-test with FDR <0.05. * $p \leq 0.05$, ** $p \leq 0.01$, **** $p \leq 0.0001$. **(F)** Relative abundance of checkpoint-protein positive and negative CD8$^+$ T cell states normalized to the total number of T cells in VP-MCC vs VN-MCC. Each point represents the mean cell frequency per MCC-sample. Significance was determined using Bonferroni-adjusted t-test with FDR <0.05. * $p \leq 0.05$, ** $p \leq 0.01$, **** $p \leq 0.0001$. **(G)** Grouped boxplots comparing the relative abundance of Myeloid cell subtypes between VP and VN-MCC samples relative to the total myeloid cells within all cutaneous MCC samples. Statistical significance was determined using Wilcoxon-rank sum test corrected for multiple-hypothesis testing. $*p < 0.05$, $**p < 0.01$, ***$p$<0.005, ****$p$<0.001. **(H)** Heatmaps comparing the mean z-normalized expression of selected markers within cells of the myeloid cell lineage between VP and VN-MCC samples. Non-significant differences are marked by dotted lines. Statistical significance was determined using Wilcoxon-rank sum test adjusted for multiple-hypothesis testing (BH). **(I)** Relative abundance of PD-L1+ macrophages and DC normalized to the total number of myeloid cells in VP-MCC vs VN-MCC. Each point represents the mean cell frequency per MCC-sample. Significance was determined using Bonferroni-adjusted t-test with FDR <0.05. * $p \leq 0.05$, ** $p \leq 0.01$, **** $p \leq 0.0001$. **(J)** Heatmap illustrating the enrichment of selected cell-types within each CN compared to the average global distribution of cells within cutaneous tissue samples of the MCC CODEX data set. Asterixis indicate a significant enrichment of a given cell-type within this CN as computed through the statsmodel package in python. **(K)** Relative abundance of CD8$^+$ T cells normalized to the total number of cells in a given CN between VP-MCC vs VN-MCC samples. Each point represents the mean cell frequency per MCC-sample. Significance was determined using Bonferroni-adjusted t-test with FDR <0.05. * $p \leq 0.05$, ** $p \leq 0.01$, **** $p \leq 0.0001$. **(L)** Heatmap illustrating the enrichment of selected myeloid cell subtypes within each CN compared to the average global distribution of cells within cutaneous tissue samples of the MCC CODEX data set. Asterixis indicate a significant enrichment of a given cell-type within this CN as computed through the statsmodel package in python. **(M)** Waterfall plots summarizing the log-fold enrichment of PD-L1/PD1 pairs within all CNs between VP and VN-MCC tumors amongst cutaneous MCC tumor samples (left) or log-fold enrichment of CXCL9/CXCR3 pairs within all CNs between VP and VN-MCC tumors amongst cutaneous MCC samples (right). Statistical significance was determined using Wilcoxon-rank sum test adjusted for multiple hypothesis testing (BH). $*p < 0.05$, $**p < 0.01$.

**Supplemental Fig.10: Analytical framework used to identify clinical and molecular features associated with distant metastasis. Related to Fig.5.**

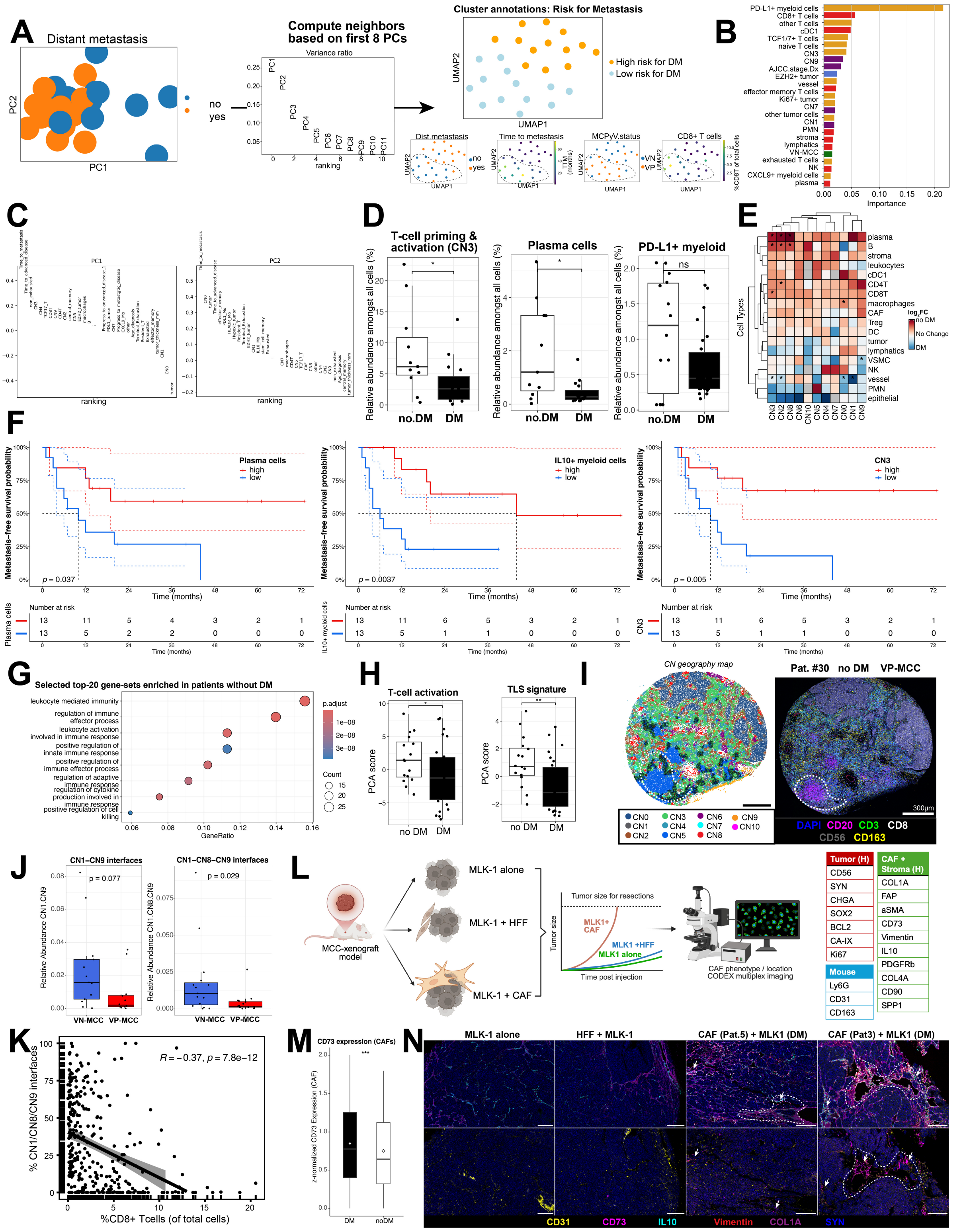


**(A)** Schematic of the analytical framework used to integrate molecular CODEX data and clinical metadata for patients with local stage I/II disease who received standard-of-care treatment. Shown is the initial computation of principal-components (left), choosing the first n=8 principal components for Leiden-clustering of integrated features (center) and exploration of features in UMAP dimension (right). **(B)** Relative importance of molecular and clinical features to the clustering results shown in the UMAP in (A). **(C)** Relative contribution of features to PC1 and PC2. **(D)** Boxplots illustrating the relative abundance of T-cell activation CN3, plasma cells and PD-L1$^{+}$ myeloid cells amongst all cells stratified by patients with / without distant metastasis. Each point represents the mean frequency from four TMA cores per tumor sample per patient. Significance was assessed using Bonferroni-corrected t-test with FDR<0.05. **(E)** Heatmap showing the relative enrichment of cell-types within a given CN between patients with initial local disease who developed (blue) or did not develop distant metastasis (red). Asterixis indicate statistically significant results as assessed by Wilcoxon-rank sum test adjusted for multiple hypothesis testing (BH). **(F)** Kaplan-Meier survival plots of DMFS probabilities stratified by cell type abundancies that were dichotomized using the median frequency of each investigated cell type (plasma cells, IL10$^+$ myeloid cells or CN3-frequency). **(G)** Dotplot illustrating selected results from gene-set enrichment analysis derived from LCMseq. Results were selected from the top-20 gene sets enriched in patients without distant metastasis and sorted by log-fold change difference and filtered adjusted p-values <0.01. **(H)** Boxplots comparing PCA scores for genes linked to T-cell activation or TLS-formation (see supplemental Table 6) derived from LCMseq. Statistical significance was determined using Wilcoxon-rank sum test corrected for multiple hypothesis testing (BH). **(I)** Representative example of a VP-MCC case with no distant metastasis showcasing the TLS-associated spatial context encompassing interfaces of CN3-CN4-CN5 using a CN-geography map (left) and corresponding multiplex immunofluorescence image (right). Interfaces are highlighted by black outline in CN geography map and by white dotted line in fluorescence image. Scale bar applies to both panels. **(J)** Boxplots depicting the relative abundance of CN-interfaces including CN1-CN9 (left) and CN1-CN8-CN9 (right) between VP and VN-MCC. Statistical significance was determined using Wilcoxon-rank sum test corrected for multiple-hypothesis testing (BH). Shown are mean relative abundances across all MCC samples from patients with initial local disease. **(K)** Scatterplot depicting the negative association of the spatial context, comprising CN1/CN8/CN9 interfaces and the infiltration by CD8+ T cells amongst all investigated tumor cores (n=356). Spearman correlation coefficient and results from statistical testing are shown. **(L)** Schematic summarizing the experimental design for the analysis of MCC xenografts from MLK-1-CAF co-cultures (left) including the CODEX multiplex imaging panel used for CAF phenotype detection. **(M)** Boxplots comparing the mean expression of CD73 in CAFs between patients who did or did not develop distant metastasis following initial local disease. Statistical significance was determined using Wilcoxon-rank sum test. **(N)** Representative examples from CODEX imaging of MCC xenografts depicting the enrichment of CD73+ CAFs together with CD31+ vessels at the tumor boundary of xenografts that did metastasize to distant sites. Scale bars of 200µm apply to all panels.

**Supplemental Fig.11: Dissection of spatial and phenotypical features linked to HPV-driven immunomodulation of the TME and patient outcomes in HNSCC. Related to Fig.6.**

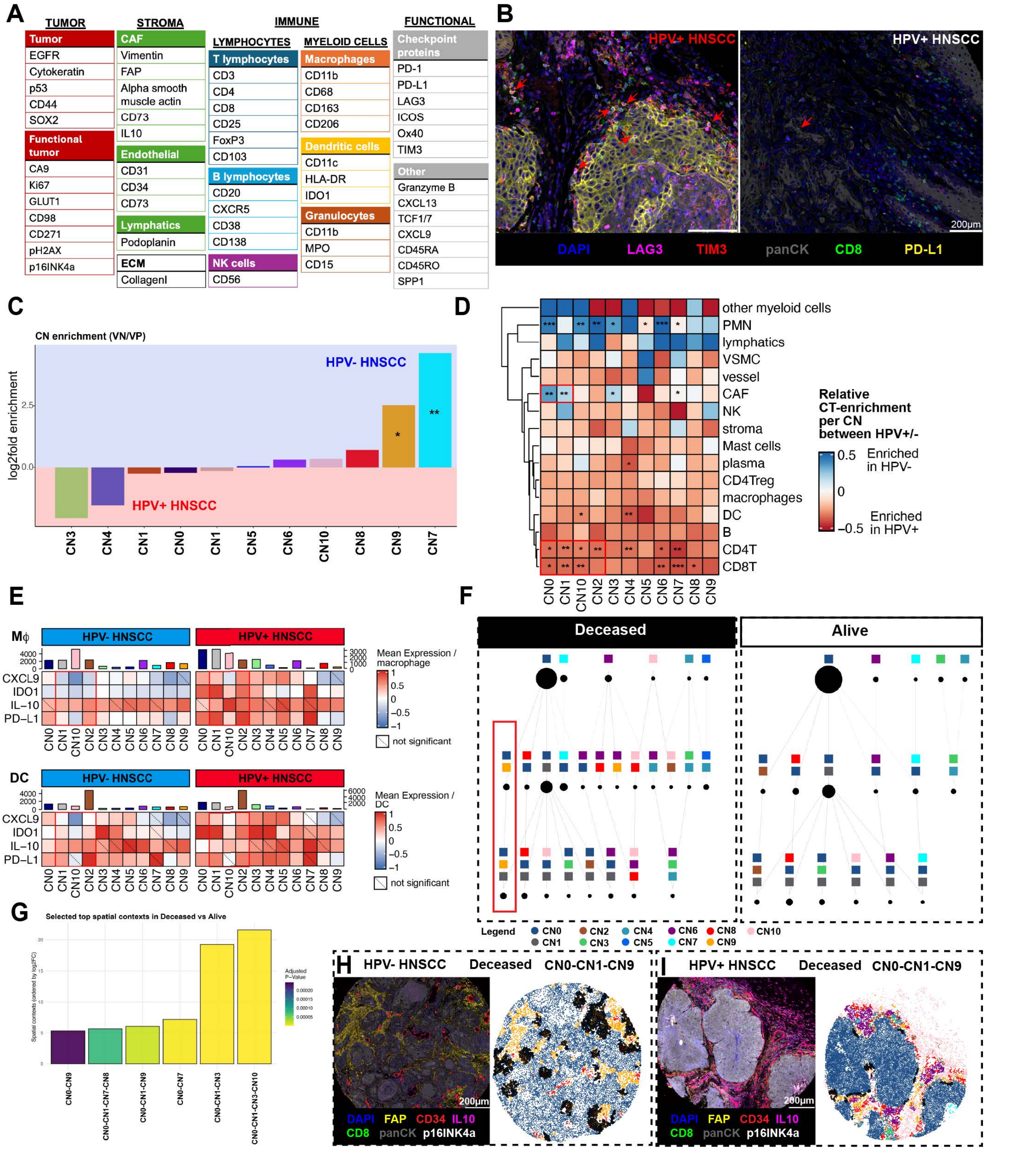


**(A)** Schematic summary of markers included in the CODEX multiplex imaging study of the OPSCC patient cohort. **(B)** Representative examples of T-cell phenotypes found in HPV+ OPSCC and HPV- OPSCC patients. **(C)** Waterfall plot depicting the log-fold enrichment of CNs between HPV+ and HPV- OPSCC patients. Statistical significance was determined using Wilcoxon-rank sum test adjusted for multiple hypothesis testing (BH). $*p < 0.05$, $**p < 0.01$. **(D)** Heatmap showing the relative enrichment of cell-types within a OPSCC CNs between HPV+ OPSCC (red) or HPV- OPSCC (blue). Asterixis indicate statistically significant results as assessed by Wilcoxon-rank sum test adjusted for multiple hypothesis testing (BH). **(E)** Heatmaps comparing the mean z-normalized expression of selected markers of macrophages (top) or dendritic cells (bottom) within OPSCC CNs between HPV+ vs HPV-OPSCC samples. Non-significant differences are marked by dotted lines. Statistical significance was determined using Wilcoxon-rank sum test adjusted for multiple-hypothesis testing (BH). **(F)** Spatial context maps showing the top10 characteristic CN-CN interfaces in OPSCC patients who deceased within the follow-up period (left). The CAF-enriched CN9 intersection at the tumor boundary CN1 (red box) is highlighted. **(G)** Barplots summarizing the differential enrichment of CN-CN interfaces in patients who deceased following OPSCC diagnosis compared to patients who did not decease within the follow-up period. Colors indicate results from statistical testing using adjusted p-values as computed through the R-package *DESeq2*. **(H)** Representative example of a HPV- OPSCC case who deceased within the follow-up period showcasing the CAF-enriched tumor boundary spatial context encompassing interfaces of CN0-CN1-CN9 as a raw CODEX image (left) or the corresponding CN-geography map (right). Interfaces are highlighted by black outline in CN geography map. Scale bar applies to both panels. **(I)** Representative example of a HPV+ OPSCC case who deceased within the follow-up showcasing the CAF-enriched tumor boundary spatial context encompassing interfaces of CN0-CN1-CN9 as a raw CODEX image (left) or the corresponding CN-geography map (right). Interfaces are highlighted by black outline in CN geography map. Scale bar applies to both panels.

**Supplemental Fig.12: Selection of patients with advanced MCC and identification of clinical and molecular biomarkers associated with ICB-response. Related to Figure 7.**

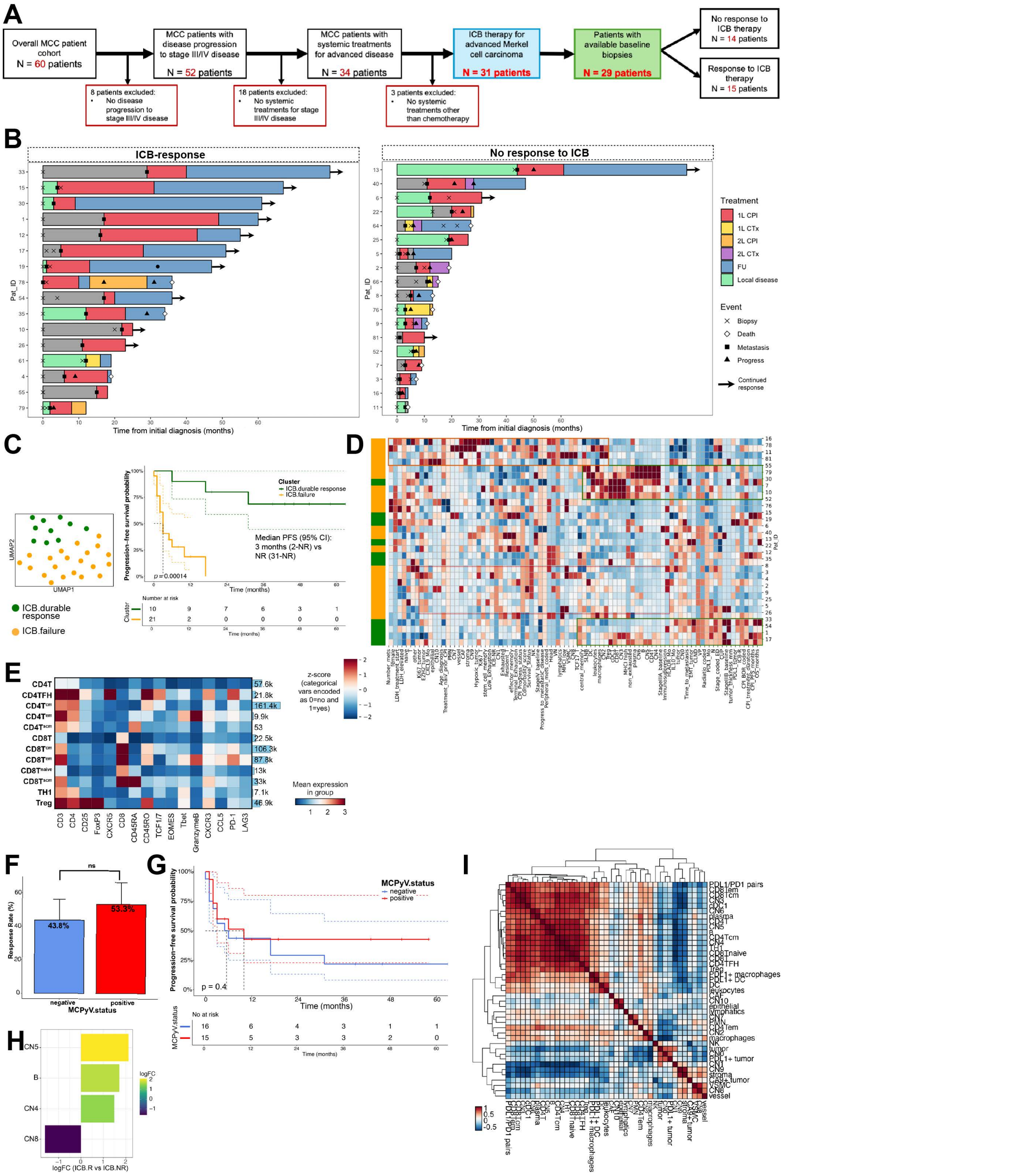


**(A)** CONSORT diagram showing the selection criteria applied to the initial cohort of 60 MCC patients to identify patients who responded to ICB therapy. Related to Figure 7A. **(B)** Swimmer plots depicting the individual course of disease and treatment for all patients that received systemic treatments for advanced MCC disease (n=34). Patients are stratified by response for first-line treatments. **(C)** UMAP depicting patient clusters following clinical-molecular data integration in low-dimensional space (left) with corresponding Kaplan-Meier plot illustrating the sharp separation of patient clusters based on integrated clinical and molecular features (right). Statistical significance was determined using log-rank test. Dotted lines indicate 95% confidence intervals. Related to Figure 7C. **(D)** Heatmap depicting clinical and molecular feature distribution across patients with advanced stage III/IV disease that were treated with ICB with rows and columns clustered by Euclidian distances. Colored boxes highlight selected clusters within the heatmap. Related to Figure 7C. **(E)** Heatmap depicting T cell memory subtypes, their absolute counts within the CODEX dataset and mean log-normalized expression of key phenotypic markers. Related to supplemental Table 5. **(F)** Bar plot with SEM comparing the response rate to immune-checkpoint inhibitors between VP and VN-MCC patients with advanced stage MCC. Statistical significance was determined using Wilcoxon-rank sum test. Related to Figure 7F. **(G)** Kaplan Meier plot for progression-free survival of patients with advanced stage MCC stratified by MCPyV-status. Statistical significance was determined using log-rank test. Related to Figure 7F. **(H)** Bar chart illustrating shared molecular predictors of ICB response between VP and VN-MCC patients amongst all parameters were p<0.05. Related to Figure 7F. **(I)** Heatmap illustrating the correlation of molecular features by relative abundance. Correlation was determined using Spearman correlation analysis and features were sorted according to hierarchical clustering. Related to Figure 7C.

**Supplemental Fig.13: Spatial distribution of $CD8^+$ central memory T cells within the immune TME of MCC. Related to Fig.7.**

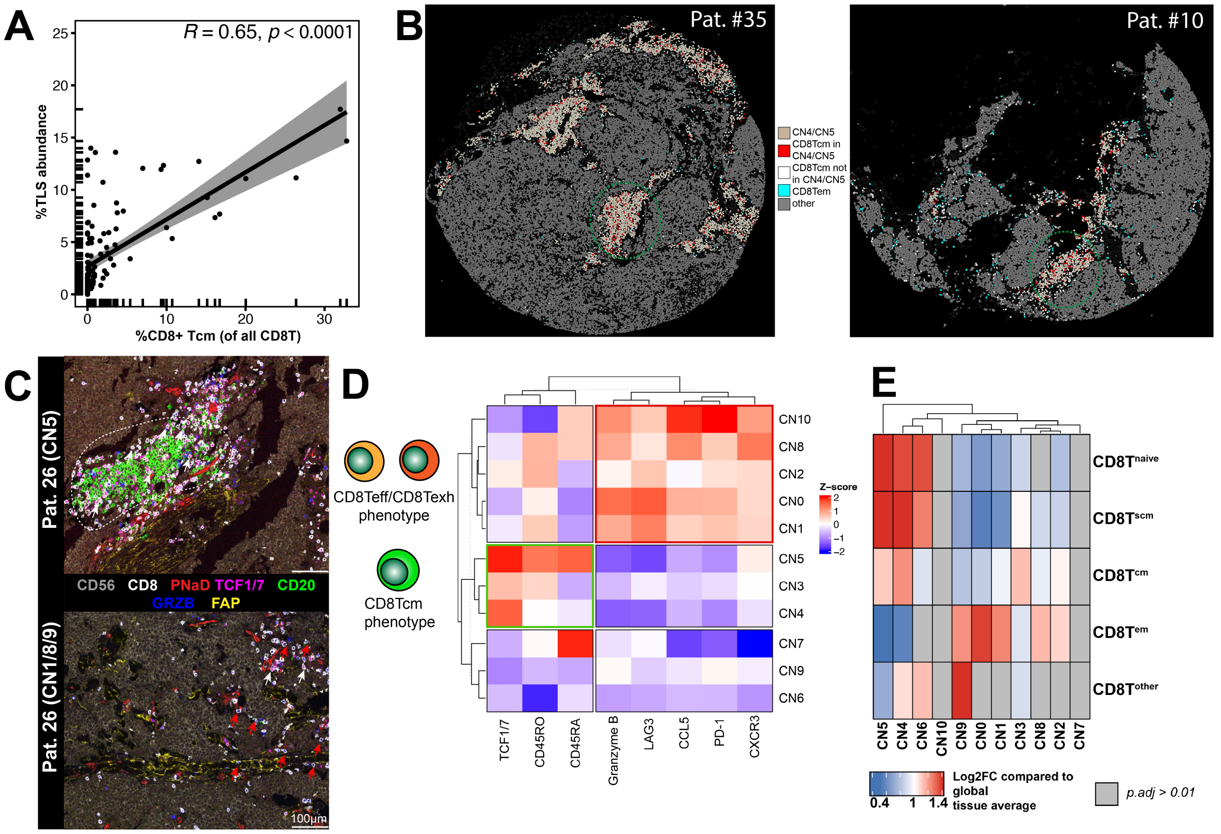


**(A)** Scatterplot showing the correlation between the abundance of CD8+ central memory T cells and the presence of B-cell enriched TLS CNs. Correlation was quantified using Spearman´s r. **(B)** Representative example showcasing the enrichment of central memory CD8+ T cells in TLS-linked CNs (red), while effector CD8+ T cells were predominantly found outside of these niches (cyan). TLS are highlighted in green circle. **(C)** Representative images showing the distribution of $CD8^+$ T cell memory subsets at different locations within the MCC TME, specifically highlighting the enrichment of effector-memory T cells at the tumor boundary (CN1/CN8/CN9; red arrows) (bottom) and $TCF1/7^+$ central memory $CD8^+$ T cells within TLS-associated CNs (white arrows) (top). Scale bar, 100µm. **(D)** Heatmap of log2-fold enrichment of markers used for T cell memory identification (left and top annotation) by cellular neighborhoods (y-axis labels) ordered by hierarchical clustering. Naïve-like memory T cells cluster in CN3-CN5, whereas T cells with an enrichment of markers indicative of an effector phenotype cluster within tumor-centric CN0, CN1, CN2, CN8, and CN10. **(E)** Relative enrichment of $CD8^+$ T cell memory phenotypes within each CN compared to global tissue average of $CD8^+$ T cell memory subsets. Statistical significance was determined using binomial testing corrected for multiple-hypothesis testing (BH).

### Supplemental Fig.14: Skin-specific validation of cellular, transcriptional and spatial features linked to ICB response in advanced MCC patients. Related to Fig.7.

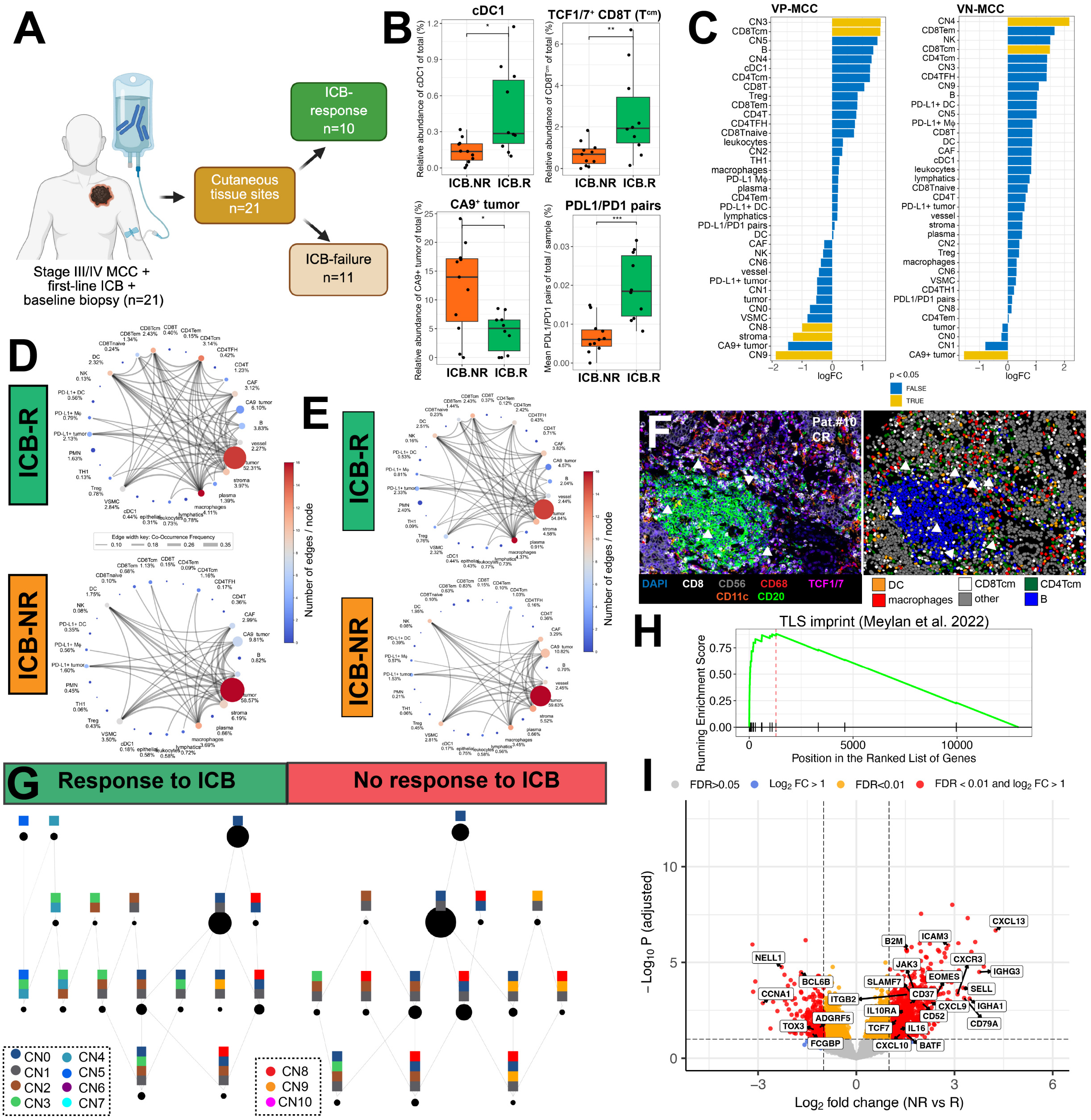


**(A)** Schematic summarizing the cohort of ICB-treated MCC patients with available cutaneous MCC baseline tumor samples. **(B)** Boxplots depicting relative abundance of molecular features associated with ICB-response in patients with only cutaneous tissue samples. Statistical significance was determined using Wilcoxon-rank sum test corrected for multiple-hypothesis testing (BH). $*p < 0.05$, $**p < 0.01$, *** $p<0.005$. **(C)** Barplots summarizing the key cellular and architectural features sorted by log-fold change difference associating with ICB-response in VP-MCC patients (left) and VN-MCC patients (right) filtered for cutaneous tissue samples only. Statistical significance was determined using a generalized linear model from the glm-package in R. **(D)** Circos plot illustrating cell-cell interactions that are significantly enriched in ICB-treated advanced MCC patients stratified by ICB response (response: top; no response: bottom). Size of edges depicts the strength of the interactions whereas the size of the nodes shows the relative abundance of cell-types compared to the global cell-type distribution. **(E)** Circos plot illustrating cell-cell interactions that are significantly enriched in ICB-treated advanced MCC patients with only cutaneous tissue samples stratified by ICB response (response: top; no response: bottom). Size of edges depicts the strength of the interactions whereas the size of the nodes shows the relative abundance of cell-types compared to the global cell-type distribution. **(F)** Representative example (left) and corresponding cell-type geography map showcasing the significant interaction between CD8Tcm, DC and B-cells observed particularly in proximity to TLS in ICB-response patients. Scale bars, 100µm. **(G)** Spatial context map depicts the enrichment of characteristic CN-interfaces in cutaneous samples of ICB-responders (left) and ICB-non-responders (right). Color codes are shown below. Size of nodes depicts the relative abundance of these CN-CN interfaces amongst the overall dataset. **(H)** Gene-set enrichment plot showing the enrichment of the TLS imprint signature in ICB-responders in only using cutaneous tissue samples. **(I)** Volcano plots of selected genes significantly enriched in cutaneous samples of ICB responders with FDR < 0.01 as assessed by LCM-seq.
