## Supplementary material for "Spatially organized inflammatory myeloid-CD8^+^ T cell aggregates linked to Merkel-cell Polyomavirus driven Reorganization of the Tumor Microenvironment": Data Figs.S1-S5

[1] Department of Microbiology and Immunology, Stanford University School of Medicine, 94305, CA, USA
[2] Department of Pathology, Stanford University School of Medicine, 94305, CA, USA
[3] Department of Dermatology, University Medical Center Mainz, 55131, Germany
[4] Institute of Pathology, University Medical Center Mainz, 55131, Germany
[5] Molecular Biosciences/Cancer Biology Program, Heidelberg University, Heidelberg, 48232, Germany
[6] German Cancer Research Center, DKFZ, Heidelberg, 48232, Germany
[7] Department of Cancer Immunology and Virology, Dana Farber Cancer Institute, Boston, 02215, MA, USA
[8] Department of Hematology, University Medical Center Heidelberg, Heidelberg, 48232, Germany
[9] Institute for Medical Microbiology, Virology and Hygiene, University Medical Center Hamburg-Eppendorf, Hamburg, Germany
[10] Virology Unit, Department of Translational Medicine, University of Piemonte Orientale, Novara, Italy
[11] Department of Biomedical Engineering, Duke University, 27708, NC, USA
[12] Lead contact
[13] These authors contributed equally

**Includes: Data Figs.S1-5.**

**Data Fig.S1:** Screening and Validation of CODEX antibodies, related to Fig.1, supplementary Table 1 and METHODS.

**Data Fig.S2.** H&E overview of the multi-tumor microarray used for validation of CODEX antibody panel. Related to Fig.1 and METHODS.

**Data Fig.S3.** Multiplex immunofluorescence overview of the multi-Tumor TMA used for validation of CODEX antibody panel. Related to Fig.1 and METHODS.

**Data Fig.S4.** Summary of all markers validated for the CODEX study. Related to Fig.1 and METHODS.

**Data Fig.S5.** Cell type annotations for the single-cell RNA-sequencing data. Related to Data Fig.S3 and S4. (A) Dotplot summarizing cell-type defining genes and total cell counts. (B) Illustration of cell-type defining markers in tSNE projection.

**Data Fig.S1: Screening and Validation of CODEX antibodies, related to Fig.1, Supplemental Table 1 and METHODS**. Following conjugation of purified antibodies to DNA oligonucleotides, those antibody-oligonucleotide conjugates were tested individually along with cross-validation in standard IHC using the same, non-conjugated antibody clone. Clones, manufacturer and staining specifications are listed along each antibody with examples for IHC staining, CODEX staining (false grey color fluorescence images), and similar areas on independent H&E-stained sections are shown. Brightness and contrast adjusted. Scale bars, 100 μm for CODEX and standard IHC images. Scale bars, 200 μm for H&E-stained images.

**Staining specifications**

**Antigen: alpha SMA**
**Clone: polyclonal**
Company:
abcam (ab5694)
Tissue: Breast
Dilution:
IHC: 1:150
CODEX: 1:100
CODEX Oligo: 69-ATTO550

**Antigen: B7-H3 (CD276)**
**Clone: D9M2L**
Company:
Cell signaling technologies (#58798)
Tissue: Tonsil
Dilution:
IHC: 1:100
CODEX: 1:40
CODEX Oligo: 17-alexa647

**Antigen: CA-9**
**Clone: polyclonal**
Company:
R&D Systems (AF2188)
Tissue: Head-and-neck cancer
Dilution:
IHC: 1:100
CODEX: 1:50
CODEX Oligo: 21-ATTO550

**Antigen: CCL5**
**Clone: E9S2K**
Company:
Cell signaling technologies (#57219)
Tissue: Tonsil
Dilution:
IHC: 1:200
CODEX: 1:50
CODEX Oligo: 11-alexa647

**Antigen: CD3**
**Clone: MRQ-39**
Company:
Cell marque (custom)
Tissue: Tonsil
Dilution:
IHC: 1:
CODEX: 1:100
CODEX Oligo: 77-ATTO550

**Antigen: CD4**
**Clone: EPR6855**
Company:
abcam (ab181724)
Tissue: Tonsil
Dilution:
IHC: 1:100
CODEX: 1:100
CODEX Oligo: 20-alexa647

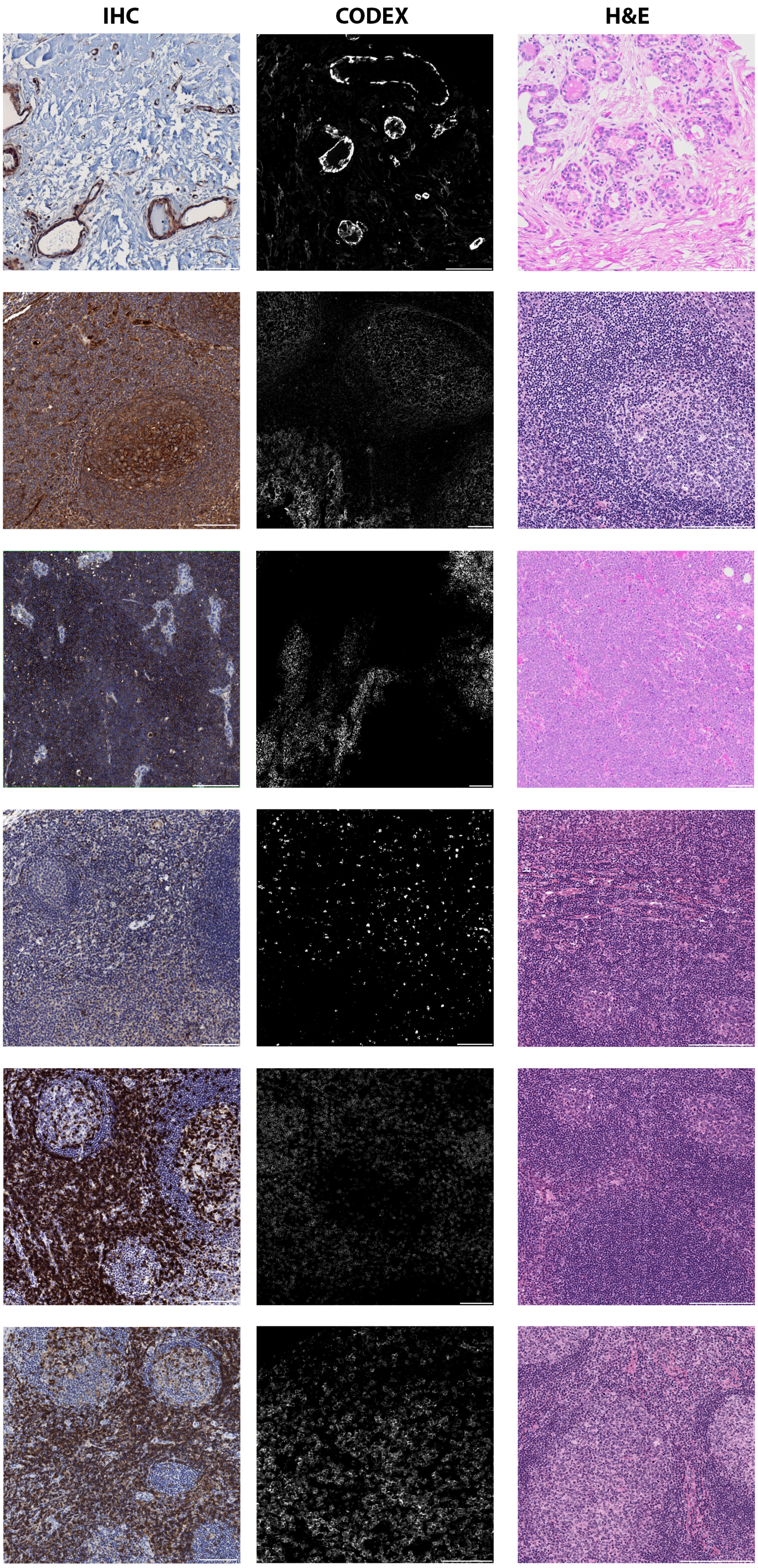

**Staining specifications**

**Antigen: CD8**
**Clone: C8/144B**

**Company:**
Cell marque (custom)

**Tissue: Tonsil**

**Dilution:**
IHC: 1:100
CODEX: 1:50

**CODEX Oligo: 8-ATTO550**

**Antigen: CD11b**
**Clone: EPR1344**

**Company:**
abcam (ab209970)

**Tissue: Tonsil**

**Dilution:**
IHC: 1:100
CODEX: 1:200

**CODEX Oligo: 28-ATTO550**

**Antigen: CD11c**
**Clone: EP1347Y**

**Company:**
abcam (ab216655)

**Tissue: Tonsil**

**Dilution:**
IHC: 1:200
CODEX: 1:150

**CODEX Oligo: 49-ATTO550**

**Antigen: CD15**
**Clone: MMA**

**Company:**
BD (559045)

**Tissue: Tonsil**

**Dilution:**
IHC: 1:100
CODEX: 1:1200

**CODEX Oligo: 14-alexa750**

**Antigen: CD20**
**Clone: IGEL/773**

**Company:**
Novus Biologicals (NBP2-54591)

**Tissue: Tonsil**

**Dilution:**
IHC: 1:250
CODEX: 1:200

**CODEX Oligo: 48-ATTO550**

**Antigen: CD21**
**Clone: SP186**

**Company:**
abcam (ab240987)

**Tissue: Tonsil**

**Dilution:**
IHC: 1:100
CODEX: 1:50

**CODEX Oligo: 44-alexa647**

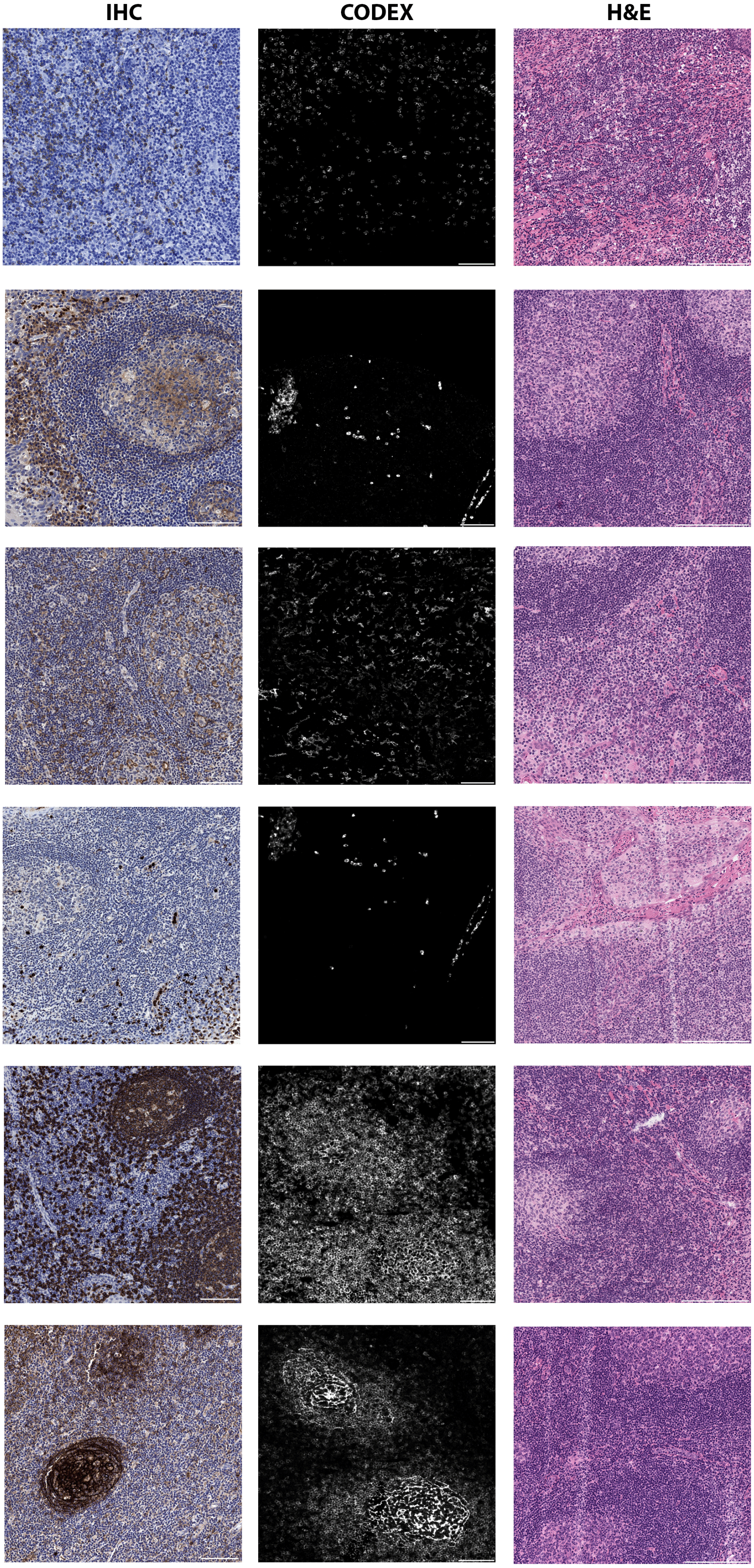

**Staining specifications** | **IHC** | **CODEX** | **H&E**

**Antigen: CD25**
**Clone: 4C9**
Company:
Cell Marque (custom)
Tissue: Tonsil
Dilution:
IHC: 1:50
CODEX: 1:75
CODEX Oligo: 24-ATTO550

**Antigen: CD31**
**Clone: C31.3+C31.7+C31.10**
Company:
Novus Biologicals (NBP2-47785)
Tissue: Tonsil
Dilution:
IHC: 1:100
CODEX: 1:200
CODEX Oligo: 68-alexa647

**Antigen: CD38**
**Clone: E7Z8C**
Company:
Cell signaling technologies (#43382)
Tissue: Tonsil
Dilution:
IHC: 1:100
CODEX: 1:150
CODEX Oligo: 66-alexa750

**Antigen: CD45**
**Clone: 2B11+PD7/26**
Company:
Novus Biologicals (Q-412898)
Tissue: Tonsil
Dilution:
IHC: 1:400
CODEX: 1:150
CODEX Oligo: 36-ATTO550

**Antigen: CD45RA**
**Clone: HI100**
Company:
Biolegend (304102)
Tissue: Tonsil
Dilution:
IHC: 1:50
CODEX: 1:250
CODEX Oligo: 72-ATTO550

**Antigen: CD45RO**
**Clone: UCH-L1**
Company:
Biolegend (304202)
Tissue: Tonsil
Dilution:
IHC: 1:50
CODEX: 1:50
CODEX Oligo: 2-alexa647

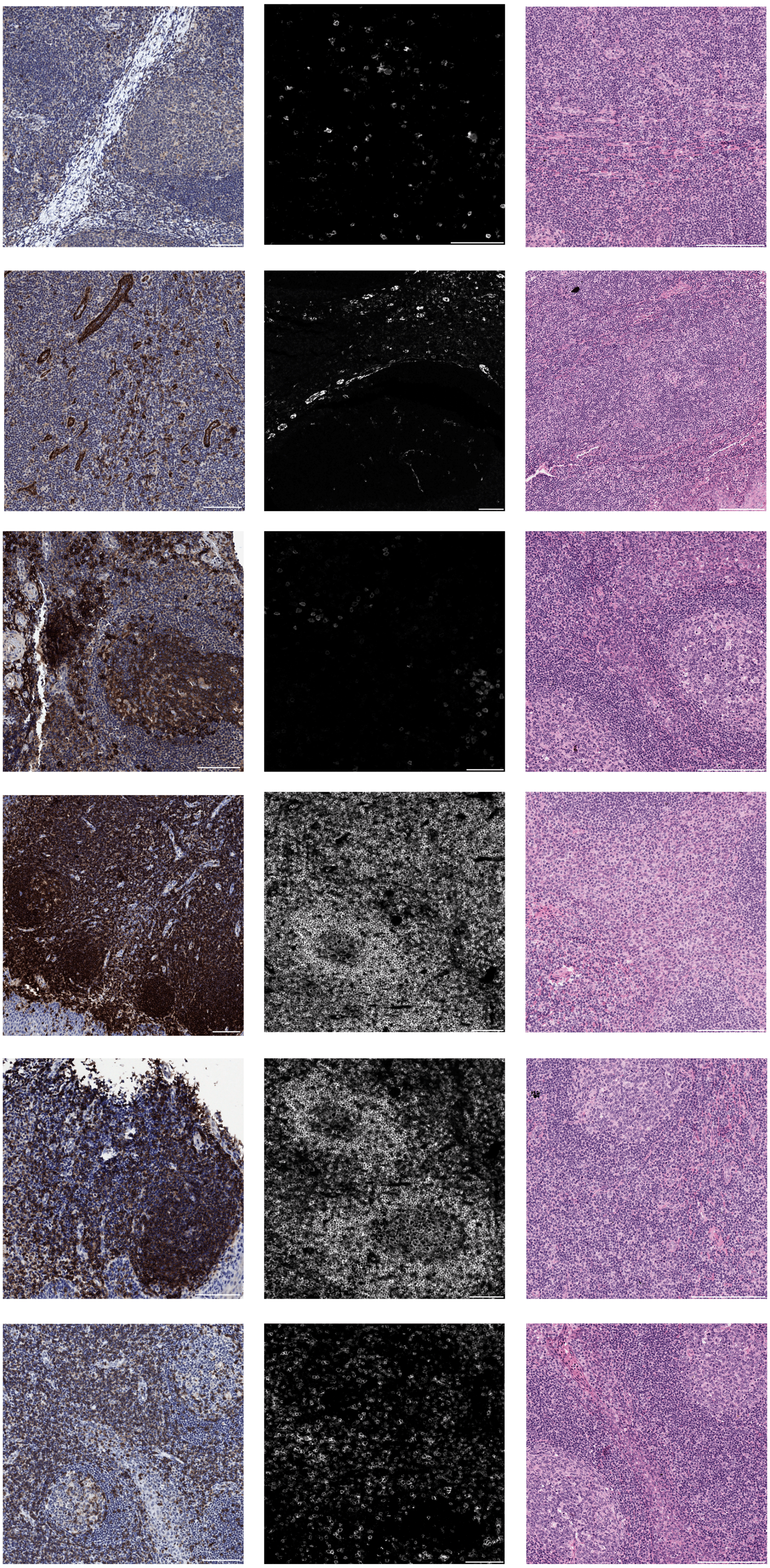

**Staining specifications**

**Antigen: CD56**
**Clone: MRQ-42**
Company:
Cell Maque (custom)
Tissue: Merkel cell carcinoma
Dilution:
IHC: 1:200
CODEX: 1:50
CODEX Oligo: 29-alexa647

**Antigen: CD57**
**Clone: HNK1**
Company:
Biolegend (359602)
Tissue: Tonsil
Dilution:
IHC: 1:100
CODEX: 1:100
CODEX Oligo: 30-ATTO550

**Antigen: CD68**
**Clone: KP-1**
Company:
Biolegend (916104)
Tissue: Tonsil
Dilution:
IHC: 1:100
CODEX: 1:200
CODEX Oligo: 70-alexa750

**Antigen: CD73**
**Clone: polyclonal (AF5795)**
Company:
R&D Systems (AF5795)
Tissue: Tonsil
Dilution:
IHC: 1:100
CODEX: 1:150
CODEX Oligo: 75-ATTO550

**Antigen: CD103**
**Clone: EPR22590-27**
Company:
abcam (ab254201)
Tissue: Tonsil
Dilution:
IHC: 1:500
CODEX: 1:50
CODEX Oligo: 38-alexa647

**Antigen: CD163**
**Clone: EDHu-1**
Company:
Novus Biologicals (NB110-40686)
Tissue: Tonsil
Dilution:
IHC: 1:100
CODEX: 1:50
CODEX Oligo: 45-ATTO550

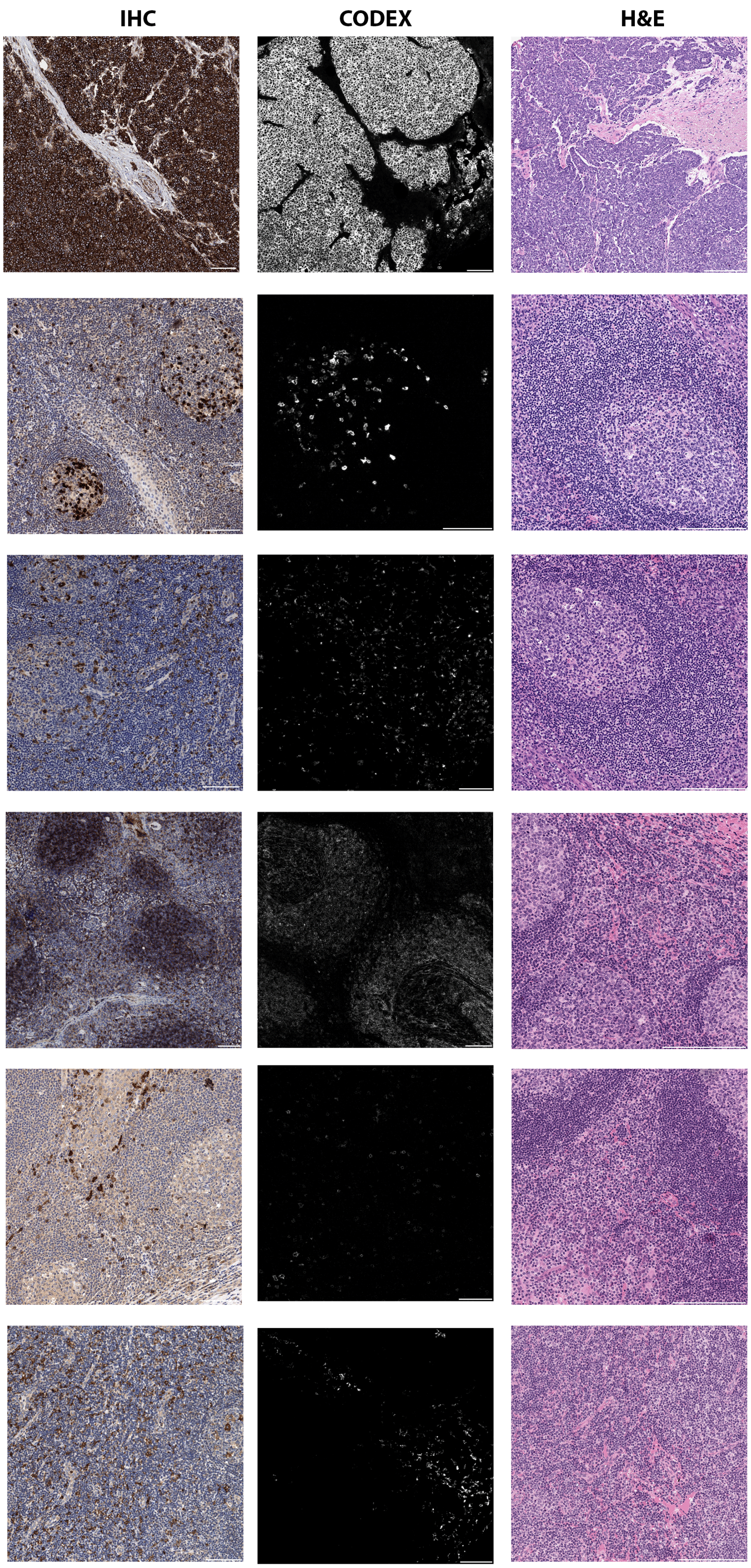

**Staining specifications** | **IHC** | **CODEX** | **H&E**

**Antigen: CD206**
**Clone: E2L9N**

Company:
Cell signaling technologies (#49243)

Tissue: Tonsil

Dilution:
IHC: 1:200
CODEX: 1:100

CODEX Oligo: 55-ATTO550

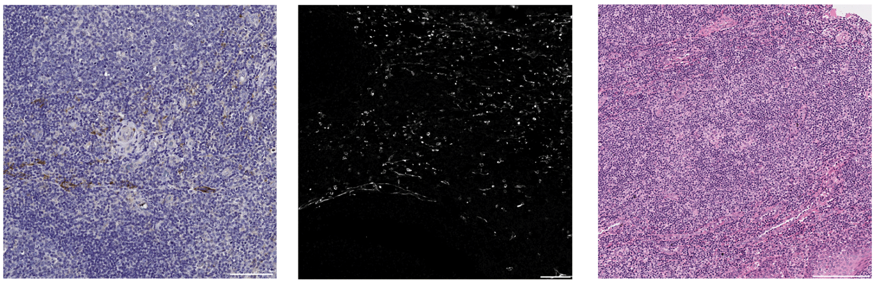

**Antigen: Chromogranin A**
Clone: LK2H10+PHE5+
CGA/414

Company:
Novus Biologicals (NBP2-34674)

Tissue: Merkel cell carcinoma

Dilution:
IHC: 1:200
CODEX: 1:50

CODEX Oligo: 80-ATTO550

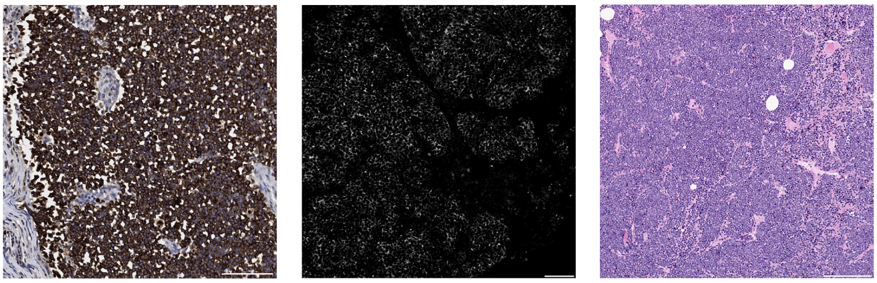

**Antigen: CLEC9A**
**Clone: EPR22324**

Company:
abcam (ab245121)

Tissue: Tonsil

Dilution:
IHC: 1:500
CODEX: 1:50

CODEX Oligo: 74-alexa647

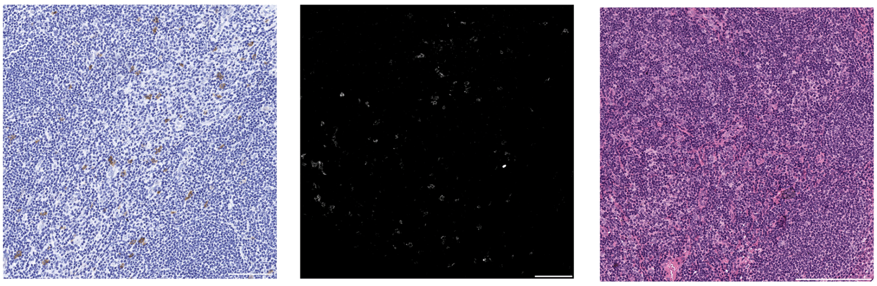

**Antigen: Collagen IV**
**Clone: EPR20966**

Company:
abcam (ab226485)

Tissue: Tonsil

Dilution:
IHC: 1:500
CODEX: 1:150

CODEX Oligo: 33-alexa647

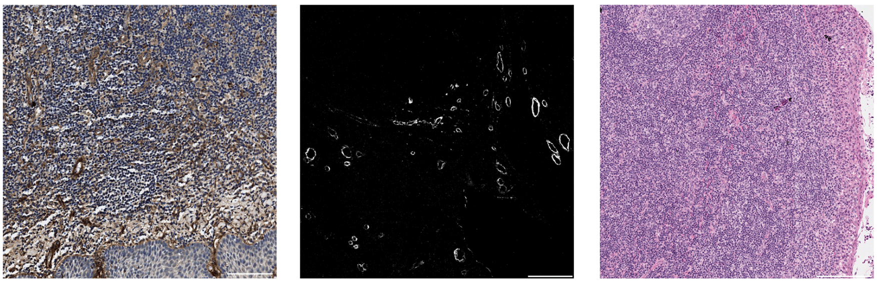

**Antigen: CXCL9**
**Clone: EPR26512-11**

Company:
abcam (ab290654)

Tissue: Tonsil

Dilution:
IHC: 1:100
CODEX: 1:40

CODEX Oligo: 58-alexa647

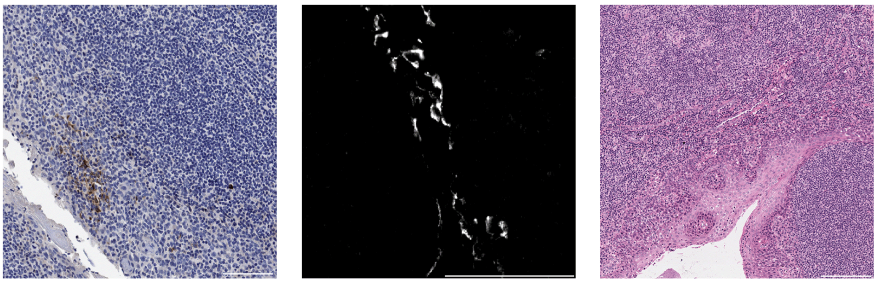

**Antigen: CXCL10**
**Clone: EPR24674-84**

Company:
abcam (ab306588)

Tissue: Tonsil

Dilution:
IHC: 1:1000
CODEX: 1:100

CODEX Oligo: 57-ATTO550

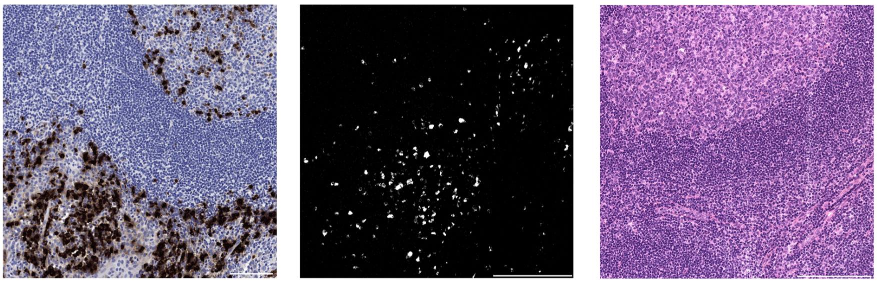

**Staining specifications** | **IHC** | **CODEX** | **H&E**

**Antigen: CXCL13**
**Clone: polyclonal**

Company:
R&D Systems (AF801)

Tissue: Tonsil

Dilution:
IHC: 1:50
CODEX: 1:100

CODEX Oligo: 41-alexa647

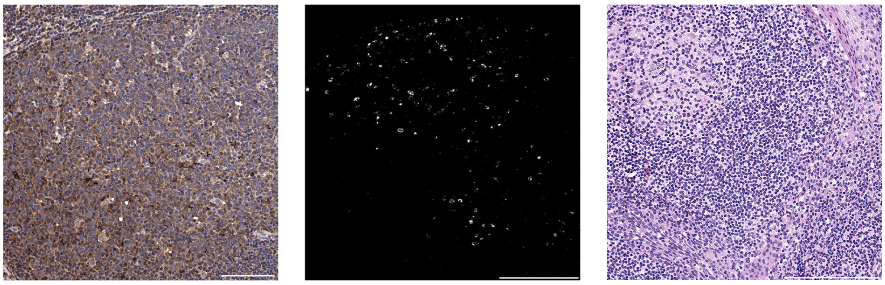

**Antigen: CXCR3**
**Clone: EPR25373-32**

Company:
abcam (ab288446)

Tissue: Tonsil

Dilution:
IHC: 1:500
CODEX: 1:50

CODEX Oligo: 59-alexa647

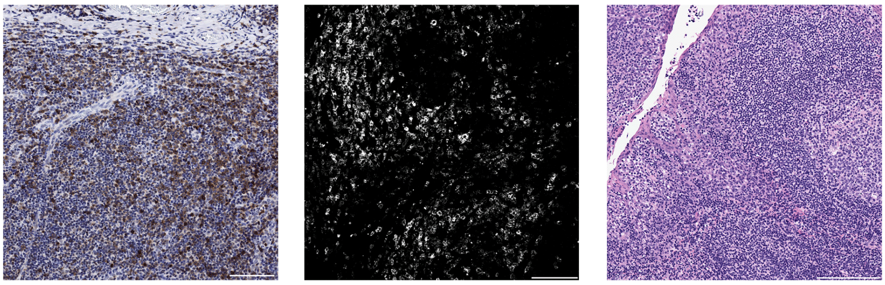

**Antigen: CXCR5**
**Clone: EPR23463-30**

Company:
abcam (ab272936)

Tissue: Tonsil

Dilution:
IHC: 1:5000
CODEX: 1:50

CODEX Oligo: 63-alexa647

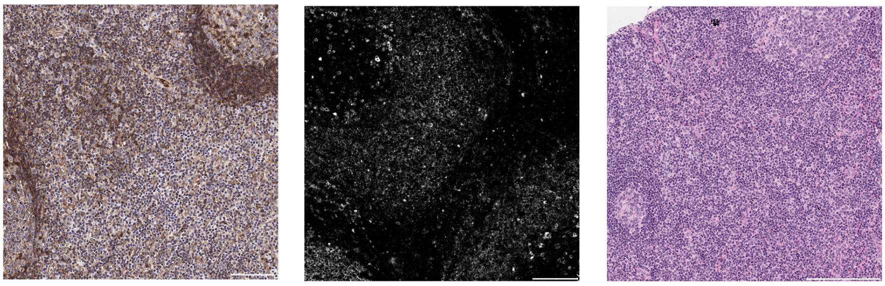

**Antigen: Cytokeratin 20**
**Clone: EPR1622Y**

Company:
abcam (ab219589)

Tissue: Merkel cell carcionma

Dilution:
IHC: 1:100
CODEX: 1:100

CODEX Oligo: 3-ATTO550

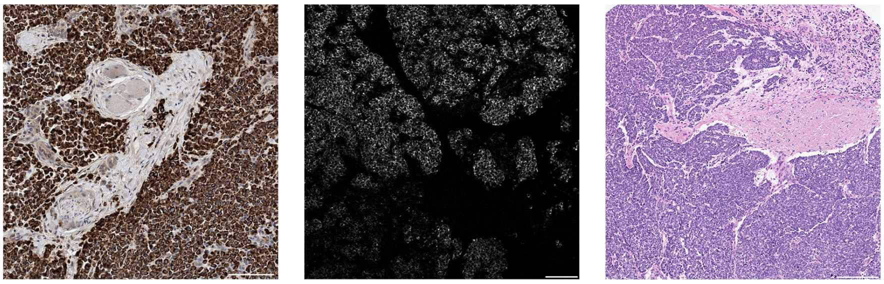

**Antigen: EOMES**
**Clone: EPR21950-24**

Company:
abcam (ab261913)

Tissue: Tonsil

Dilution:
IHC: 1:500
CODEX: 1:50

CODEX Oligo: 67-alexa647

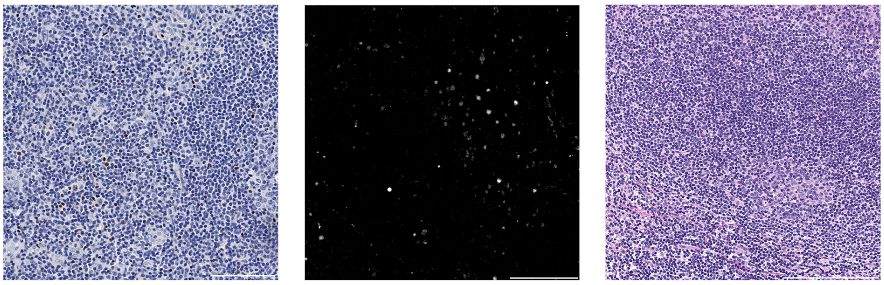

**Antigen: EZH2**
**Clone: D2C9**

Company:
Cell signaling technologies (#16098)

Tissue: Tonsil

Dilution:
IHC: 1:50
CODEX: 1:50

CODEX Oligo: 21-alexa647

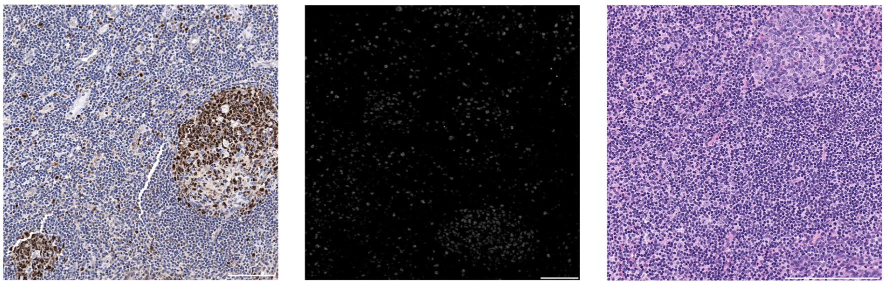

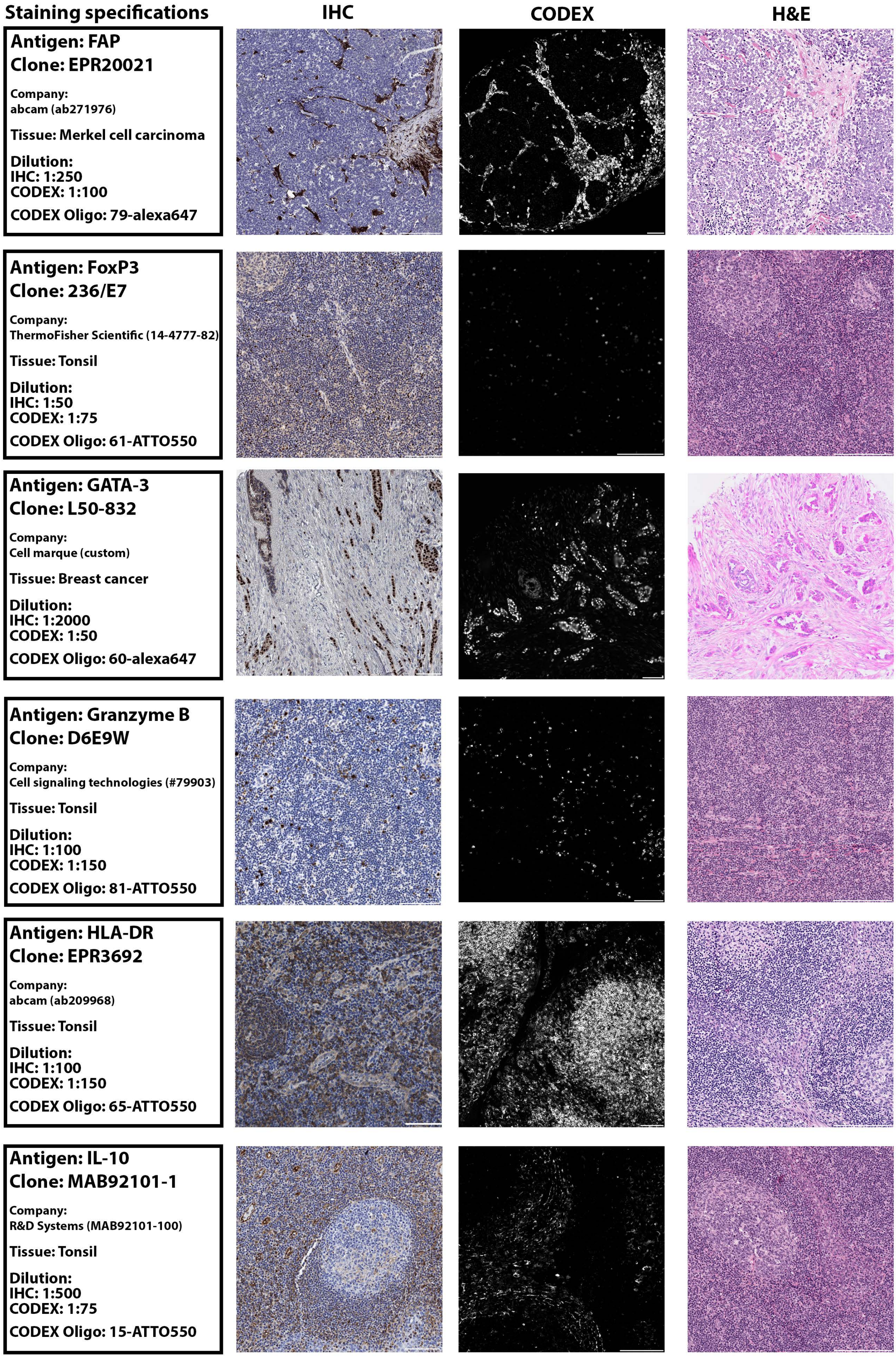
Staining specifications
IHC
CODEX
H&E
Antigen: FAP
Clone: EPR20021
Company:
abcam (ab271976)
Tissue: Merkel cell carcinoma
Dilution:
IHC: 1:250
CODEX: 1:100
CODEX Oligo: 79-alexa647
Antigen: FoxP3
Clone: 236/E7
Company:
ThermoFisher Scientific (14-4777-82)
Tissue: Tonsil
Dilution:
IHC: 1:50
CODEX: 1:75
CODEX Oligo: 61-ATTO550
Antigen: GATA-3
Clone: L50-832
Company:
Cell marque (custom)
Tissue: Breast cancer
Dilution:
IHC: 1:2000
CODEX: 1:50
CODEX Oligo: 60-alexa647
Antigen: Granzyme B
Clone: D6E9W
Company:
Cell signaling technologies (#79903)
Tissue: Tonsil
Dilution:
IHC: 1:100
CODEX: 1:150
CODEX Oligo: 81-ATTO550
Antigen: HLA-DR
Clone: EPR3692
Company:
abcam (ab209968)
Tissue: Tonsil
Dilution:
IHC: 1:100
CODEX: 1:150
CODEX Oligo: 65-ATTO550
Antigen: IL-10
Clone: MAB92101-1
Company:
R&D Systems (MAB92101-100)
Tissue: Tonsil
Dilution:
IHC: 1:500
CODEX: 1:75
CODEX Oligo: 15-ATTO550

**Staining specifications**

IHC | CODEX | H&E

**Antigen: IRF4**
**Clone: IRF4.3E4**

Company:
Biolegend (646402)

Tissue: Tonsil

Dilution:
IHC: 1:50
CODEX: 1:50

CODEX Oligo: 76-alexa750

**Antigen: IRF8**
**Clone: E6J8Q**

Company:
Cell signaling technologies (#28852)

Tissue: Tonsil

Dilution:
IHC: 1:500
CODEX: 1:50

CODEX Oligo: 43-alexa647

**Antigen: Ki67**
**Clone: B56**

Company:
BD Biosciences (556003)

Tissue: Tonsil

Dilution:
IHC: 1:200
CODEX: 1:300

CODEX Oligo: 6-alexa750

**Antigen: LAG3**
**Clone: D2G40**

Company:
Cell signaling technologies (#25848))

Tissue: Tonsil

Dilution:
IHC: 1:150
CODEX: 1:30

CODEX Oligo: 42-alexa647

**Antigen: MCPyV gp3 LT**
**Clone: Ab3**

Company:
abcam (ab202866)

Tissue: Merkel cell carcinoma

Dilution:
IHC: 1:100
CODEX: 1:50

CODEX Oligo: 51-alexa647

**Antigen: MHC-I**
**Clone: EMR8-5**

Company:
abcam (ab70328)

Tissue: Tonsil

Dilution:
IHC: 1:20.000
CODEX: 1:40

CODEX Oligo: 71-alexa750

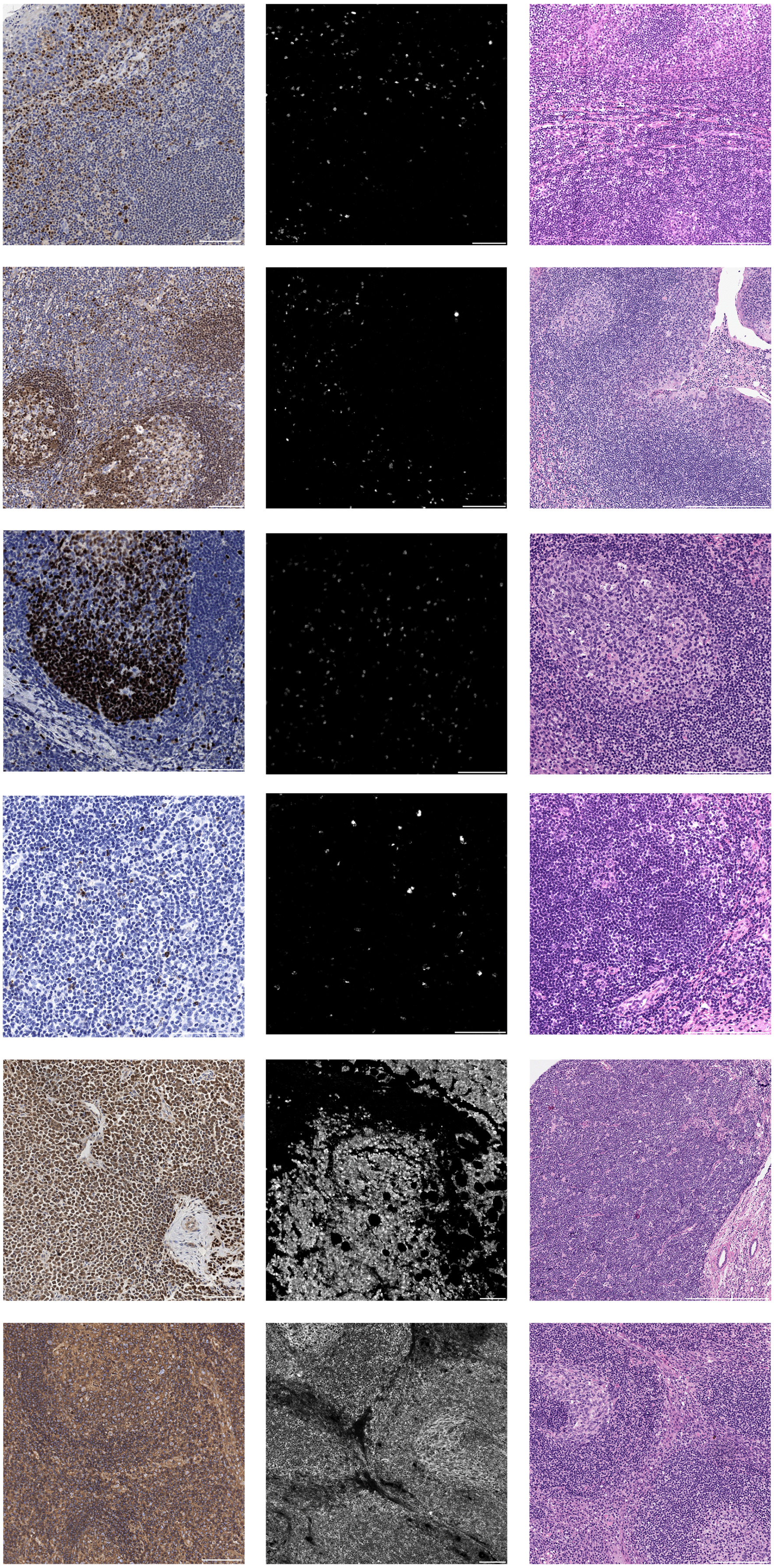

**Staining specifications**

**Antigen: MMP9**
**Clone: L51/82**
Company:
Biolegend (819701)
Tissue: Tonsil
Dilution:
IHC: 1:200
CODEX: 1:100
CODEX Oligo: 62-alexa750

**Antigen: p53**
**Clone: D-07**
Company:
Cell marque (custom)
Tissue: Tonsil
Dilution:
IHC: 1:100
CODEX: 1:50
CODEX Oligo: 52-ATTO550

**Antigen: PD-1**
**Clone: D4W2J**
Company:
Cell signaling technologies (#63815)
Tissue: Tonsil
Dilution:
IHC: 1:200
CODEX: 1:50
CODEX Oligo: 23-alexa647

**Antigen: PD-L1**
**Clone: 405.9A11**
Company:
Cell signaling technologies (#39356)
Tissue: Tonsil
Dilution:
IHC: 1:50
CODEX: 1:50
CODEX Oligo: 25-ATTO550

**Antigen: Podoplanin**
**Clone: D2-40**
Company:
Biolegend (916606)
Tissue: Tonsil
Dilution:
IHC: 1:100
CODEX: 1:50
CODEX Oligo: 32-alexa647

**Antigen: Phospho-Rb**
**Clone: D20B12**
Company:
Cell signaling technologies (#41359)
Tissue: Tonsil
Dilution:
IHC: 1:150
CODEX: 1:50
CODEX Oligo: 46-ATTO550

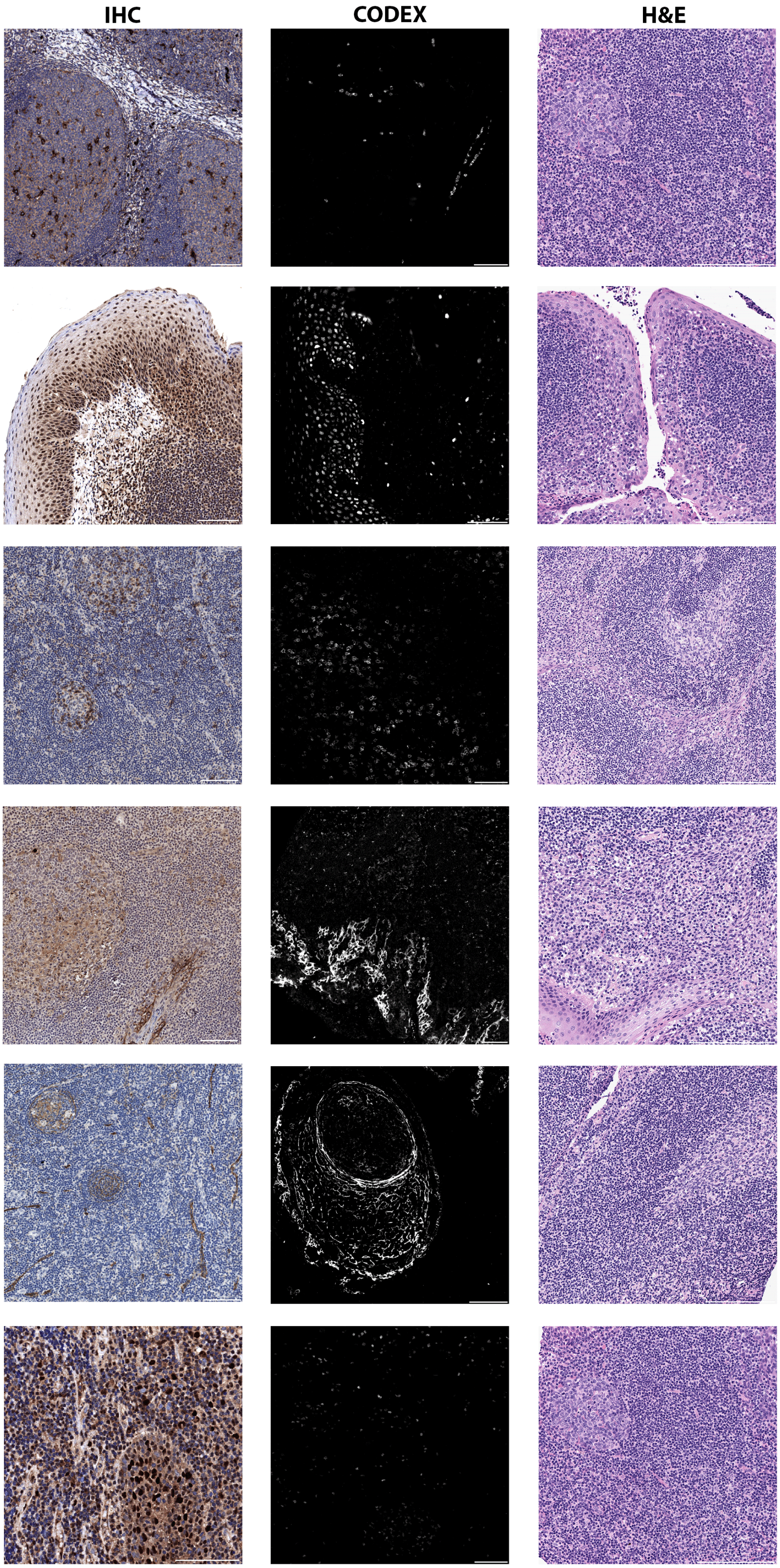

**Staining specifications**

**Antigen: Synaptophysin**
**Clone: 7H12**

Company:
Novus Biologicals (NBP1-47483)

Tissue: Merkel cell carcinoma

Dilution:
IHC: 1:200
CODEX: 1:100

CODEX Oligo: 26-ATTO550

**Antigen: Tbet**
**Clone: polyclonal**

Company:
BD Biosciences (561263)

Tissue: Tonsil

Dilution:
IHC: 1:200
CODEX: 1:50

CODEX Oligo: 5-alexa647

**Antigen: TCF1/7**
**Clone: C63D9**

Company:
Cell signaling technologies (#85942)

Tissue: Tonsil

Dilution:
IHC: 1:100
CODEX: 1:150

CODEX Oligo: 56-alexa647

**Antigen: Vimentin**
**Clone: RV202**

Company:
Novus Biologicals (NBP1-97672)

Tissue: Tonsil

Dilution:
IHC: 1:200
CODEX: 1:150

CODEX Oligo: 7-alexa647

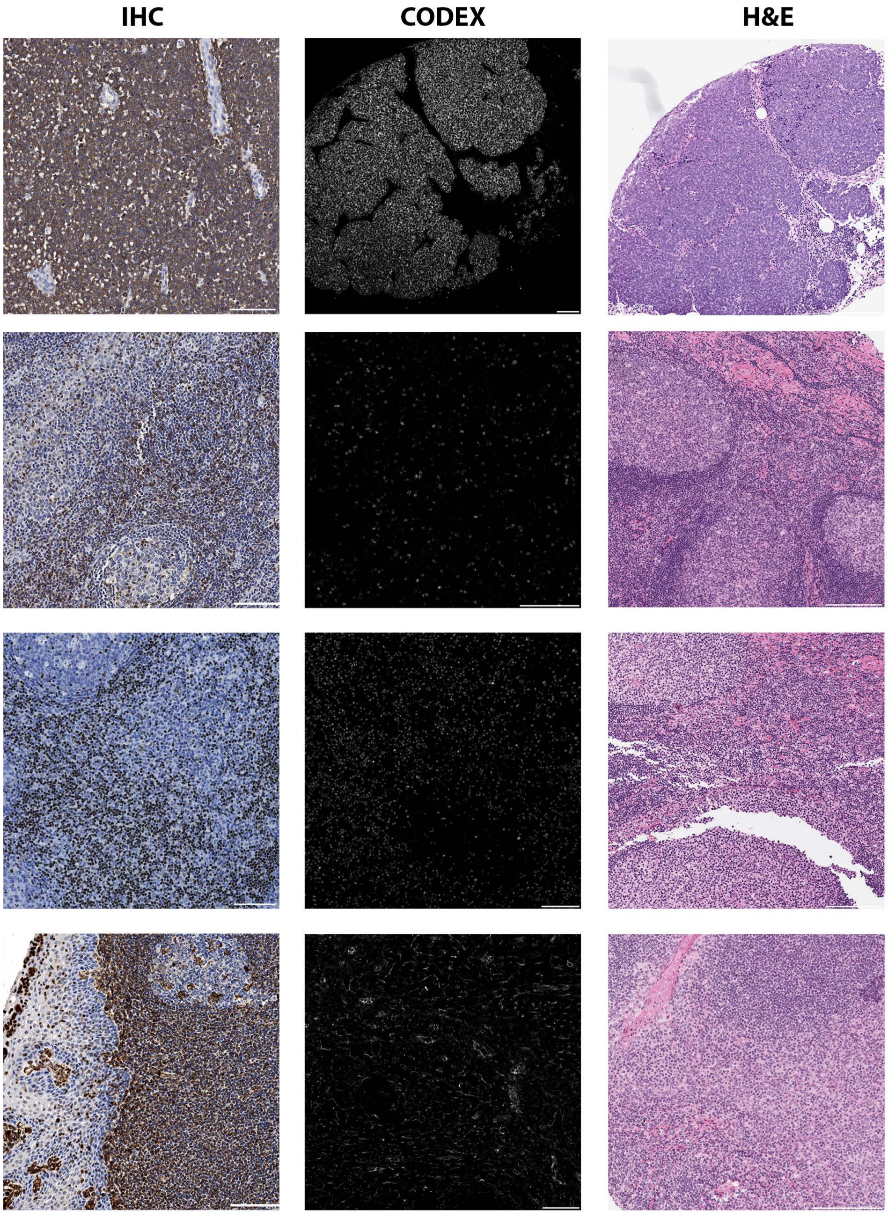

**Data Fig.S2. H&E overview of the multi-tumor microarray used for validation of CODEX antibody panel. Related to Fig.1 and METHODS.** Representative H&E-stained section of multi-tumor TMA. For details of tissues and tumor types see Supplemental Table 5. Scale bars, 1 mm.

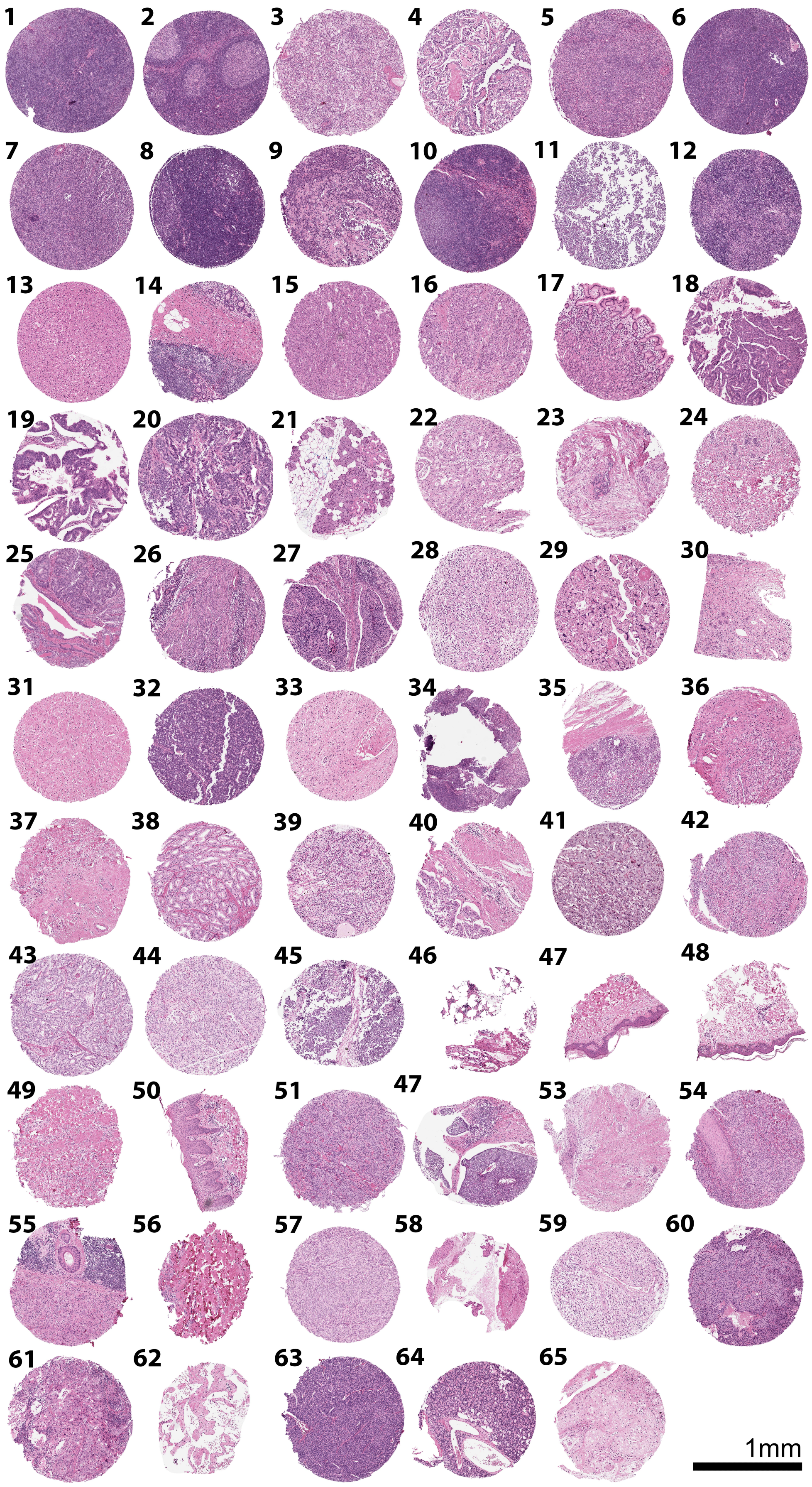

**Data Fig.S3. Multiplex immunofluorescence overview of the multi-Tumor TMA used for validation of CODEX antibody panel. Related to Fig.1 and METHODS.** Validation of the antibody panel in CODEX multiplex imaging of the multi-tumor TMA shown in Data Fig.S2.

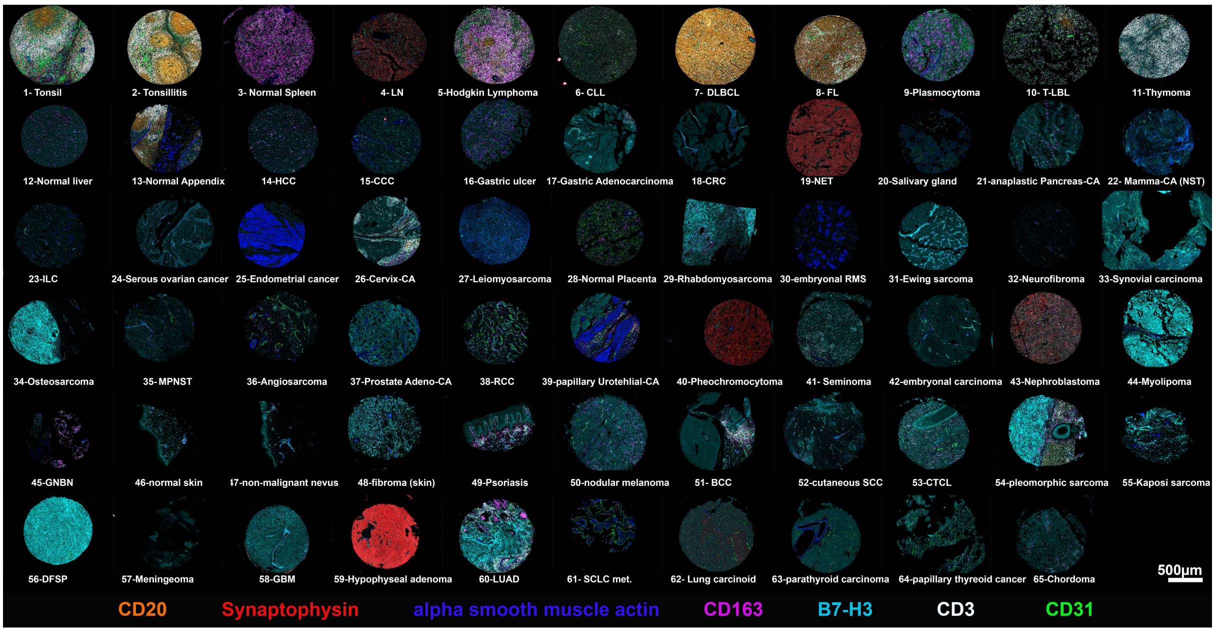

**Data Fig.S4. Summary of all markers validated for the CODEX study. Related to Fig.1 and METHODS.** Validation of the full antibody panel shown in two representative cores of one VN-MCC tumor core, which includes all markers except for MCPyV LT gp3 antigen which is shown for a VP-MCC tumor sample annotated from TMA1 core L1 on the bottom right (false grey color fluorescence images). Scale bars, 0.5 mm.

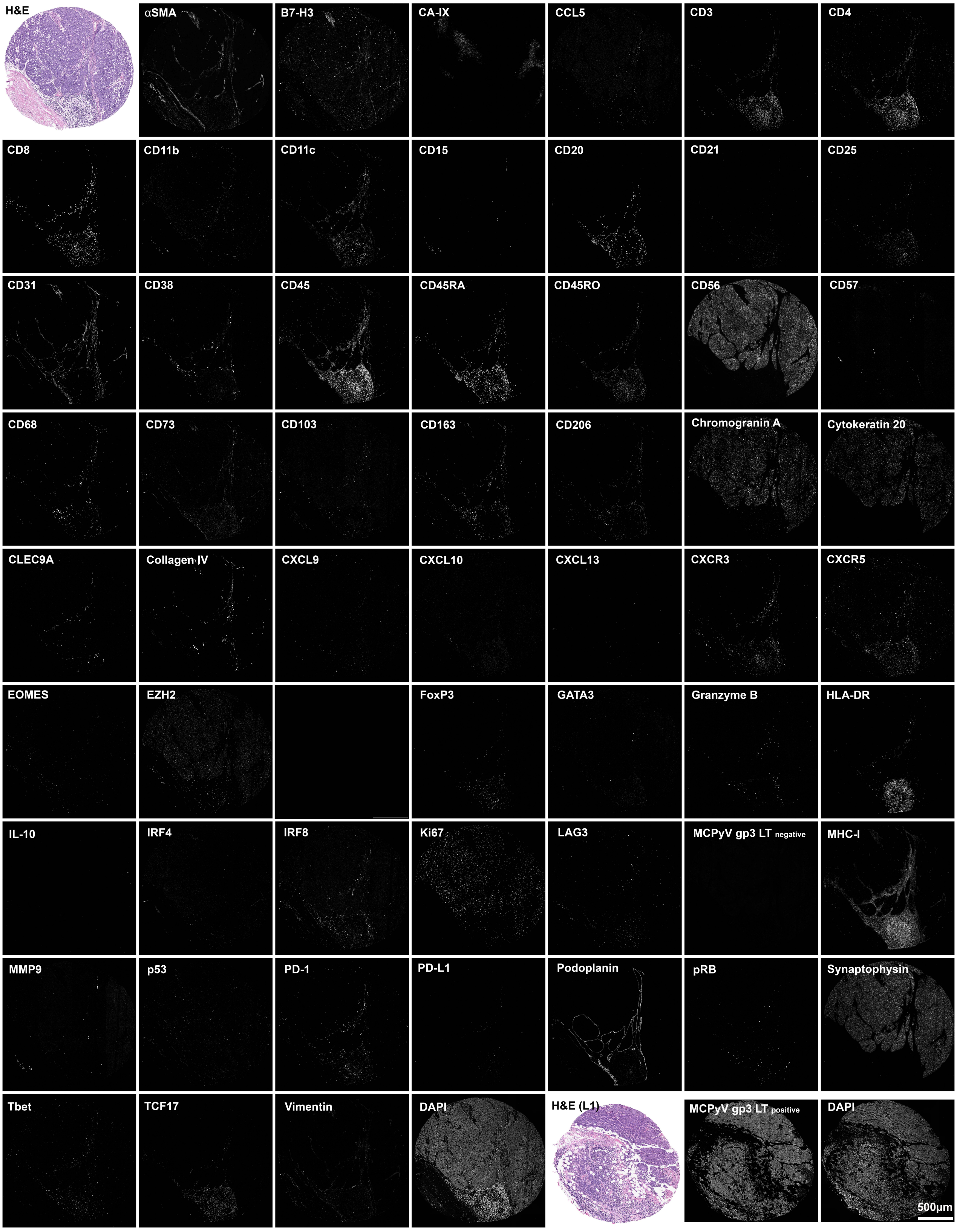

**Data Fig.S5. Cell type annotations for the single-cell RNA-sequencing data. Related to Data Fig.S3 and S4.** (A) Dotplot summarizing cell-type defining genes and total cell counts. (B) Illustration of cell-type defining markers in tSNE projection.

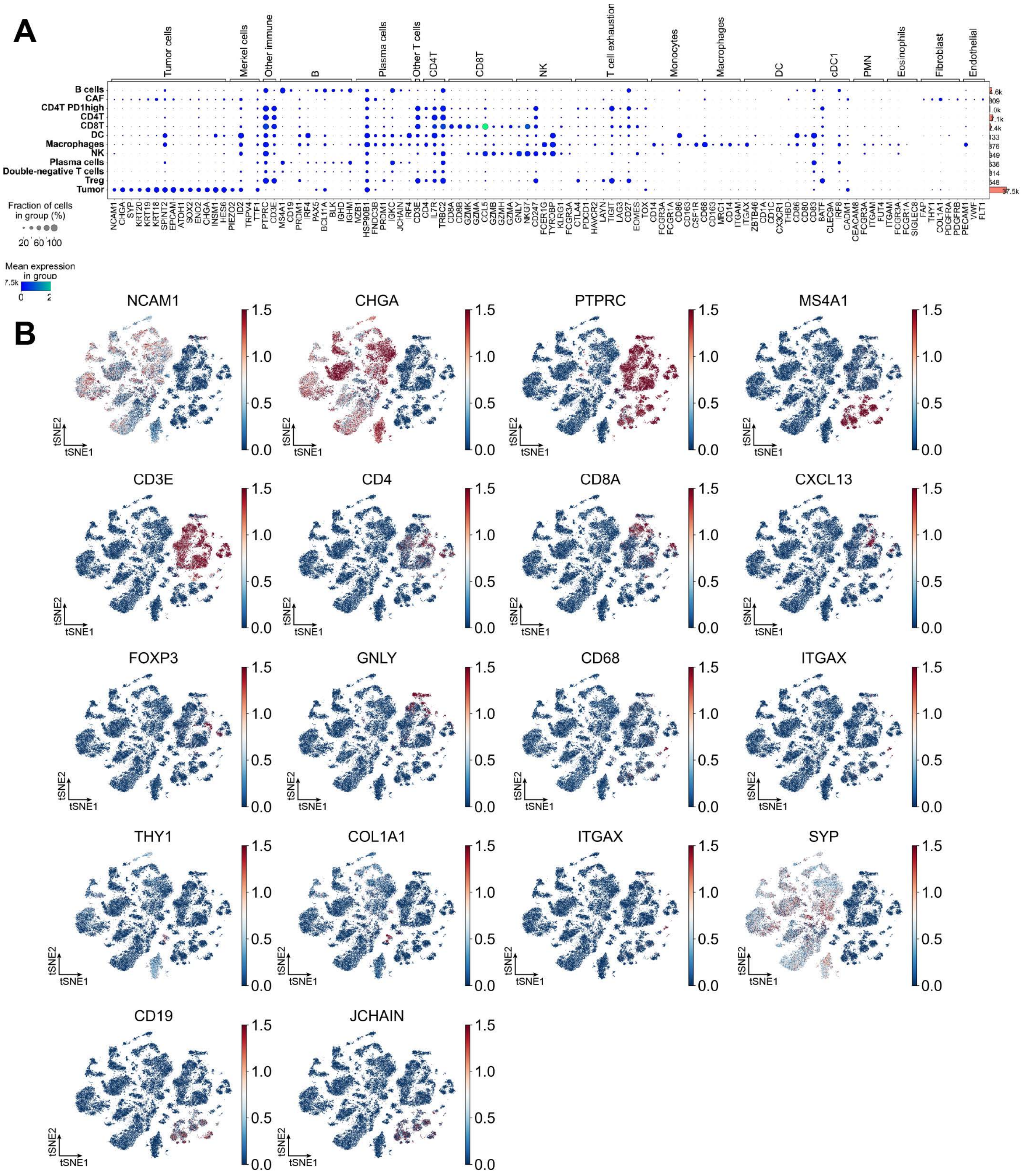
