## Supplemental Table 1 for "Spatially organized inflammatory myeloid-CD8^+^ T cell aggregates linked to Merkel-cell Polyomavirus driven Reorganization of the Tumor Microenvironment"

**Supplemental Table 1: Baseline clinicopathological characteristics stratified by MCPyV status. Related to Fig.1E.**

| | **Overall cohort** | **MCPyV positive (VP)** | **MCPyV negative (VN)** | **p-value** |
|---|---|---|---|---|
| Number of patients | 60 | 31 | 29 | - |
| **Demographics** | | | | |
| Median age at diagnosis (range) | 76.0 (44-94) | 76.0 (47-90) | 76.0 (44-94) | 0.782 |
| Gender | | | | 0.599 |
| - male | 36 (60.0%) | 20 (64.5%) | 16 (55.2%) | |
| - female | 24 (40.0%) | 11 (35.5%) | 13 (44.8%) | |
| Immunosuppression | 10 (16.7%) | 8 (25.8%) | 2 (6.9%) | 0.082 |
| ECOG performance status[1] | | | | 0.19 |
| - 0 | 22 (38.6%) | 13 (44.8%) | 9 (32.1%) | |
| - 1 | 16 (28.1%) | 10 (34.5%) | 6 (21.4%) | |
| - 2 | 18 (31.6%) | 6 (22.2%) | 12 (42.9%) | |
| - 3 | 1 (1.8%) | 0 | 1 (3.6%) | |
| **Primary tumor characteristics** | | | | |
| Localization of primary tumor | | | | **0.005** |
| - Head/neck | 20 (33.3%) | 6 (19.4%) | 14 (48.3%) | |
| - Upper limb | 12 (20.0%) | 11 (35.5%) | 1 (3.4%) | |
| - Trunk | 8 (13.3%) | 2 (6.5%) | 6 (20.7%) | |
| - Lower limb | 10 (16.7%) | 6 (19.4%) | 4 (13.8%) | |
| - CUP | 10 (16.7%) | 6 (19.4%) | 4 (13.8%) | |
| Median tumor thickness (range)[2] | 10.0mm (3.2-86.0) | 10.0mm (1.2-30.0) | 10.0mm (3.2-86.0) | 0.802 |
| MCPyV status | | | | - |
| - positive | 31 | 31 | 0 | |
| - negative | 29 | 0 | 29 | |
| T stage at diagnosis[3] | | | | 0.364 |
| - T1 | 13 (22.8%) | 10 (32.3%) | 3 (11.5%) | |
| - T2 | 23 (40.4%) | 10 (32.3%) | 13 (50.0%) | |
| - T3 | 6 (10.5%) | 3 (9.7%) | 3 (11.5%) | |
| - T4 | 5 (8.8%) | 2 (6.5%) | 3 (11.5%) | |
| - Tx | 10 (17.5%) | 6 (19.4%) | 4 (15.4%) | |
| N stage at diagnosis | | | | 0.118 |
| - N0 | 26 (43.3%) | 13 (41.9%) | 13 (44.8%) | |
| - N1 | 11 (18.3%) | 7 (22.6%) | 4 (13.8%) | |
| - N2 | 12 (20.0%) | 3 (9.7%) | 9 (31.0%) | |
| - N3 | 11 (18.3%) | 8 (25.8%) | 3 (10.3%) | |
| AJCC stage at initial diagnosis | | | | 0.421 |
| - I | 7 (11.7%) | 5 (16.1%) | 2 (6.9%) | |
| - IIA | 18 (30.0%) | 8 (25.8%) | 10 (34.5%) | |
| - IIB | 1 (1.7%) | 0 | 1 (3.4%) | |
| - IIIA | 6 (10.0%) | 2 (6.5%) | 4 (13.8%) | |
| - IIIB | 27 (45.0%) | 16 (51.6%) | 11 (37.9%) | |
| - IV | 1 (1.7%) | 0 | 1 (3.4%) | |
| Location of tissue biopsies | | | | 0.31 |
| - primary tumor | 51/115 (44.3%) | 24/64 (37.5%) | 27/51 (52.9%) | |
| - cutaneous metastasis | 24/115 (20.8%) | 15/64 (23.4%) | 9/51 (17.6%) | |
| - LN-metastasis | 28/115 (24.3%) | 17/64 (26.6%) | 11/51 (21.6%) | |
| - Other metastasis | 12/115 (0.4%) | 8/64 (12.5%) | 4/51 (7.8%) | |
| **Treatments for all disease stages** | | | | |
| Sentinel node biopsy[4] | | | | 0.384 |
| - None | 24 (45.3%) | 10 (35.7%) | 14 (56.0%) | |
| - Tumor-free | 20 (37.7%) | 12 (42.9%) | 8 (32.0%) | |
| - Tumor-inoculated | 9 (17.0%) | 6 (21.4%) | 3 (12.0%) | |
| Radiotherapy | 42 (70.0%) | 25 (80.6%) | 17 (58.6%) | 0.091 |

| | | | | |
|---|---|---|---|---|
| Median time to advanced stage III disease (95% CI)[5] | 12.0 months (7-NR) | 19.0 months (12-NR) | 7.0 months (4-NR) | **0.047** |
| Median time to metastatic stage IV disease (95% CI)[5] | 19.0 months (10-NR) | NR (19.0 – NR) | 7.0 months (4.0-NR) | **0.0034** |
| **Treatment for advanced disease stages** | | | | |
| AJCC stage at baseline<br>- IIIA<br>- IIIB<br>- IV | <br>7 (13.2%)<br>14 (26.9%)<br>32 (60.4%) | <br>4 (15.4%)<br>8 (30.8%)<br>14 (53.8%) | <br>3 (11.1%)<br>5 (17.2%)<br>18 (66.7%) | 0.6 |
| Multifocal metastatic disease (>2 metastatic sites) | 12 (36.4%) | 6 (37.5%) | 6 (35.3%) | 1.0 |
| Complete lymph node dissection (LAD) | 26 (43.3%) | 15 (48.4%) | 11 (37.9%) | 0.446 |
| Elevated LDH-serum levels at baseline (>245U/ml) | 21 (70.0%) | 10 (71.4%) | 11 (68.8%) | 0.596 |
| Systemic first-line (1L) treatments<br>- None*<br>- Chemotherapy<br>- Checkpoint-inhibitor therapy | <br>18 (34.6%)<br>5 (9.6%)<br>29 (55.8%) | <br>8 (30.8%)<br>4 (15.4%)<br>14 (53.8%) | <br>10 (38.5%)<br>1 (3.8%)<br>15 (57.7%) | 0.491 |
| Median duration of first-line treatments (range) | 6.0 (0-32) | 5.5 (1-32) | 6.0 (0-31) | 0.934 |
| Progress upon first-line treatments | 31 (64.6%) | 14 (58.3%) | 17 (70.8%) | 0.547 |
| Progression-free survival (95% CI) | 6.0 months (1.3-10.7) | 6.0 months (0-14.7) | 6.0 months (1.6-10.4) | 0.634 |
| Second-line treatment regimens<br>- Chemotherapy<br>- Checkpoint-inhibitor therapy<br>- Other | <br>5 (38.5%)<br>7 (53.8%)<br>1 (7.7%) | <br>3 (50.0%)<br>2 (33.3%)<br>1 (16.7%) | <br>2 (28.6%)<br>5 (71.4%)<br>0 | 0.388 |
| Progress following second line treatment | 8/12 (66.7%) | 6/6 | 2/6 (33.3%) | 0.061 |
| Subsequent treatment lines<br>- None<br>- One or more | <br>5<br>5 | <br>3<br>5 | <br>2<br>0 | - |
| **Follow-up data** | | | | |
| Median overall survival (95% CI) | NR | NR | NR | 0.054 |
| Deceased | 15 (25.0%) | 5 (16.1%) | 10 (34.5%) | 0.139 |
| 1-year survival probability (95% CI) | - | 100% (100%) | 73.9% (58.9-92.7%) | - |
| 3-year survival probability (95% CI) | - | 79.8% (65.0-98.0%) | 55.4% (37.1-82.8%) | - |
| Median Follow-up (95% CI) | 35.0 months (21.5-48.5) | 32.0 months (16.3-47.7) | 35.0 months (13.0-56.9) | 0.957 |

[1] Data for comorbidities and ECOG performance status have been available for n=29/31 VP patients and n=28/30 VN patients; [2] Data on tumor thickness was available for 38/60 patients (16 for VN and 22 for VP patients); [3] T stage has not been documented in n=3 cases. [4] Data on sentinel node biopsies have been available for n=53 patients. [5] Time to advanced disease and metastatic disease have been calculated for all patients with initial diagnosis of local stage I/II disease (n=26). *Among all patients that received no systemic treatment for advanced disease n=12 were diagnosed stage III disease and 5 were diagnosed with metastatic stage IV disease; Tests used for comparing continuous variables were Mann-Whitney U test and for categorical variables exact Fisher test.
