## Supplemental Table 2 for "Spatially organized inflammatory myeloid-CD8^+^ T cell aggregates linked to Merkel-cell Polyomavirus driven Reorganization of the Tumor Microenvironment"

**Supplemental Table 2: Univariate Cox-regression analysis for factors impacting overall survival across the entire patient cohort (n=60). Related to Fig.1E and Supplemental Table 1.**

| Parameter | | HR | 95% CI | p-value | p-value in multivariate |
|---|---|---|---|---|---|
| Gender | Male v female | 1.2 | 0.42-3.6 | 0.71 | |
| Age | | 1.02 | 0.97-1.07 | 0.49 | |
| Location of primary tumor | Other vs Head | 0.44 | 0.16-1.2 | 0.12 | |
| Tumor thickness | | 1.01 | 0.98-1.04 | 0.586 | |
| Immunosuppression | Yes vs no | 2.7 | 0.92-7.9 | 0.071 | |
| MCPyV status | Positive vs negative | 0.36 | 0.12-1.1 | 0.065 | 0.203 |
| Comorbidity score | Low vs High | 0.24 | 0.053-1 | 0.057 | 0.054 |
| T stage at initial diagnosis | T1 vs T2 vs T3 vs T4 | 2.05 | 1.3-3.3 | **0.0024** | 0.151 |
| AJCC stage at initial diagnosis | I vs II vs III vs IV | 1.6 | 0.74-3.3 | 0.24 | |
| LAD | Yes vs no | 1.63 | 0.58-4.6 | 0.35 | |
| Local radiotherapy | Yes vs no | 0.61 | 0.21-1.77 | 0.36 | |
| Distant organ metastases | Yes vs no | 3.4 | 1.2-9.7 | **0.024** | 0.519 |
| BOR | Response vs no response | 0.23 | 0.096-0.53 | **<0.001** | |
