## Supplemental Table 3 for "Spatially organized inflammatory myeloid-CD8^+^ T cell aggregates linked to Merkel-cell Polyomavirus driven Reorganization of the Tumor Microenvironment"

**Supplemental Table 3: Clinicopathological data on patients with initial local disease (stratified by progress to advanced disease). Related to Supplemental Fig.1D-E.**

| | **Patients with initial local disease** | **No progress to advanced disease** | **Progress to advanced disease** | **p-value** |
|---|---|---|---|---|
| Number of patients | 26 | 8 | 18 | - |
| **Demographics** | | | | |
| Median age at diagnosis (range) | 73.0 years (44-94) | 76.5 years (59-83) | 72.0 years (44-94) | 1.0 |
| Gender | | | | 0.216 |
| - male | 12 (46.2%) | 2 (25.0%) | 10 (55.6%) | |
| - female | 14 (53.8%) | 6 (75.0%) | 8 (44.4%) | |
| Immunosuppression | 5 (19.2%) | 0 | 5 (27.8%) | 0.281 |
| ECOG status[1] | | | | 0.405 |
| - 0 | 12 (48.0%) | 6 (75.0%) | 6 (35.3%) | |
| - 1 | 6 (24.0%) | 1 (12.5%) | 5 (29.4%) | |
| - 2 | 6 (24.0%) | 1 (12.5%) | 5 (29.4%) | |
| - 3 | 1 (4.0%) | 0 | 1 (5.9%) | |
| **Primary tumor characteristics** | | | | |
| Localization of primary tumor | | | | 0.585 |
| - Head/neck | 9 (34.6%) | 4 (50.0%) | 5 (27.8%) | |
| - Upper limb | 6 (23.1%) | 1 (12.5%) | 5 (27.8%) | |
| - Trunk | 5 (19.2%) | 2 (25.0%) | 3 (16.7%) | |
| - Lower limb | 6 (23.1%) | 1 (12.5%) | 5 (27.8%) | |
| Median tumor thickness (range)[2] | 9.2mm (1.2-86) | 10.5mm (3.0-19.0) | 8.3mm (1.2-86.0) | 0.667 |
| MCPyV status | | | | 0.395 |
| - positive | 13 (50.0%) | 5 (62.5%) | 8 (44.4%) | |
| - negative | 13 (50.0%) | 3 (37.5%) | 10 (55.6%) | |
| T stage at diagnosis | | | | 0.904 |
| - T1 | 7 (26.9%) | 2 (25.0%) | 5 (27.8%) | |
| - T2 | 13 (50.0%) | 5 (62.5%) | 8 (44.4%) | |
| - T3 | 5 (19.2%) | 1 (12.5%) | 4 (22.2%) | |
| - T4 | 1 (3.8%) | 0 | 1 (5.6%) | |
| AJCC stage at initial diagnosis | | | | 1.0 |
| - I | 7 (26.9%) | 2 (25.0%) | 5 (27.8%) | |
| - IIA | 18 (69.2%) | 6 (75.0%) | 12 (66.7%) | |
| - IIB | 1 (3.8%) | 0 | 1 (5.6%) | |
| Progress to advanced disease | 18/26 | 0 | 18 (100%) | **<0.001** |
| Median time to advanced disease (95% CI) | 8.5 months (0-15.3) | NA | 8.5 months (0-15.3) | - |
| Progress to metastatic disease | 15/28 | 0 | 15 (83.3%) | **<0.001** |
| Median time to metastatic disease (95% CI) | 19.0 months (0-24.5) | NR | 10.0 months (0-14.5) | **0.002** |
| **Treatments for all disease stages** | | | | |
| Sentinel node biopsy | | | | 0.586 |
| - None | 10 (38.5%) | 2 (25%) | 8 (44.4%) | |
| - Without tumor detection | 15 (57.7%) | 6 (75%) | 9 (50%) | |
| - Node positive | 1 (3.8%) | 0 | 1 (5.6%) | |
| Radiotherapy | 18 (69.2%) | 5 (62.5%) | 13 (72.2%) | 0.667 |
| **Treatment for advanced disease stages** | | | | |
| AJCC stage at baseline | | | | **<0.001** |
| - IIIA | 1 (5.6%) | NA | 1 (5.6%) | |
| - IIIB | 2 (11.1%) | | 2 (11.1%) | |
| - IV | 15 (83.3%) | | 15 (83.3%) | |

| | | | | |
|---|---|---|---|---|
| Systemic first-line (1L) treatments<br>- None**<br>- Chemotherapy<br>- Checkpoint-inhibitor therapy | <br>3 (16.7%)<br>3 (16.7%)<br>12 (66.7%) | <br>NA | <br>3 (16.7%)<br>3 (16.7%)<br>12 (66.7%) | - |
| Best overall response***<br>- PD<br>- SD<br>- PR<br>- CR/NED | <br>3 (20.0%)<br>7 (46.7%)<br>3 (20.0%)<br>2 (13.3%) | <br>NA | <br>3 (20.0%)<br>7 (46.7%)<br>3 (20.0%)<br>2 (13.3%) | - |
| Progress upon first-line treatments | 13 (86.6%) | NA | 13 (86.6%) | - |
| **Follow-up data** | | | | |
| Median overall survival (95% CI) | NR | NR | NR | 0.138 |
| Deceased | 5 (19.2%) | 0 | 5 (27.8%) | 0.150 |
| Median Follow-up (95% CI) | 31.0 months (12.4-49.6) | 21.0 months (11.3-30.7) | 47.0 months (15.5-78.5) | **0.047** |

[1] ECOG performance status was available for 25/26 patients; [2] Data on tumor thickness was unavailable for 2 patients; **Among all patients that received no systemic treatment for advanced disease n=1 were diagnosed stage III disease and 2 were diagnosed with metastatic stage IV disease; *** BOR was assessed among all patients that received systemic first line treatments (n=15). Tests used for comparing continuous variables were Mann-Whitney U test and for categorical variables exact Fisher test.
