## Supplemental Table 4 for "Spatially organized inflammatory myeloid-CD8^+^ T cell aggregates linked to Merkel-cell Polyomavirus driven Reorganization of the Tumor Microenvironment"

**Supplemental Table 4: Clinical factors impacting the time to metastasis for patients with initial local disease in univariate Cox-regression analysis (n=26). Related to Supplemental Fig.1D-E, and Supplemental Table 3.**

| Parameter | | HR | 95% CI | p-value | p-value in multivariate |
|---|---|---|---|---|---|
| Gender | Male v female | 1.50 | 0.54-4.15 | 0.4 | |
| Age | | 1.01 | 0.97-1.06 | 0.48 | |
| Primary tumor location | Head vs other | 1.16 | 0.38-3.3 | 0.84 | |
| Tumor thickness | | 1.08 | 1.005-1.169 | **0.037** | **0.05** |
| Immunosuppression | No vs yes | 0.84 | 0.23-3.04 | 0.788 | |
| MCPyV status | positive vs negative | 0.196 | 0.059-0.65 | **0.007** | **0.006** |
| ECOG | 0 vs ≥1 | 0.5 | 0.17-1.5 | 0.21 | |
| T stage | T1 vs T2 vs T3 vs T4 | 1.5 | 0.83-2.61 | 0.19 | |
| AJCC stage at baseline | Stage I vs Stage IIA vs Stage IIB | 2.8 | 0.74-10.26 | 0.13 | |
| Local radiotherapy | Yes vs no | 0.71 | 0.24-2.1 | 0.54 | |
| MHC-I status (baseline) | Low vs high | 1.4 | 0.17-12 | 0.74 | |
