## Supplemental Table 5 for "Spatially organized inflammatory myeloid-CD8^+^ T cell aggregates linked to Merkel-cell Polyomavirus driven Reorganization of the Tumor Microenvironment"

**Supplemental Table 5:** Markers employed for the identification of T cell and myeloid cell subsets.

| **Cell population** | **Subpopulation** | **Marker expression** |
|---|---|---|
| **CD8$^{+}$ T cells** | Resident CD8$^{+}$ T cells | CD8$^{+}$, CD103$^{+}$ |
| **CD8$^{+}$ T cells** | Naïve | CD45RA$^{high}$, CD45RO$^{-}$, TCF1/7$^{-}$, CXCR3$^{-}$ |
| | Stem-cell memory | CD45RA$^{high}$, TCF1/7$^{+}$, CD45RO$^{-}$, CXCR3$^{-}$ |
| | Central-memory | CD45RO$^{+}$, TCF1/7$^{+}$, CD45RA$^{low}$, CXCR3$^{low}$ |
| | Effector memory | CD45RO$^{+}$, TCF1/7$^{-}$, CD45RA$^{-}$, GRZB$^{high}$, CXCR3$^{high}$ |
| | Non-exhausted | PD1$^{low}$, LAG3$^{low}$, EOMES$^{-}$, |
| | “Precursor-exhausted” | PD1$^{high}$, LAG3$^{low}$, EOMES$^{low}$ |
| | “Exhausted” | PD1$^{high}$, LAG3i$^{nter}$, Tbet$^{inter}$, EOMES$^{+}$ |
| | “Terminally exhausted” | PD1$^{high}$, LAG3$^{high}$, EOMES$^{high}$ |
| **CD4$^{+}$ T cells** | **TFH CD4$^{+}$ T cells** | CD4+, PD1$^{high}$, CXCL13high |
| **CD4$^{+}$ T cells** | **TH1 CD4$^{+}$ T cells** | CD4+, Tbet$^{high}$ |
| **CD4$^{+}$ T cells** | Naïve | CD45RA$^{high}$, CD45RO$^{-}$, TCF1/7$^{-}$, CXCR3$^{-}$ |
| | Stem-cell memory | CD45RA$^{high}$, TCF1/7$^{+}$, CD45RO$^{-}$, CXCR3$^{-}$ |
| | Central-memory | CD45RO$^{+}$, TCF1/7$^{+}$, CD45RA$^{low}$, CXCR3$^{low}$ |
| | Effector memory | CD45RO$^{+}$, TCF1/7$^{-}$, CD45RA$^{-}$, GRZB$^{inter}$, CXCR3$^{high}$ |
| | Non-exhausted | PD1$^{low}$, LAG3$^{low}$, EOMES$^{-}$, |
| | “Precursor-exhausted” | PD1$^{high}$, LAG3$^{low}$, EOMES$^{low}$ |
| | “Exhausted” | PD1$^{high}$, LAG3i$^{nter}$, Tbet$^{inter}$, EOMES$^{+}$ |
| | “Terminally exhausted” | PD1$^{high}$, LAG3$^{high}$, EOMES$^{high}$ |
| **Macrophages** | CD68$^{high}$ macrophages | CD11b$^{+}$, CD68$^{+}$, HLA-DR$^{inter}$, CD163$^{-}$, CD206$^{-}$ |
| | CD163$^{high}$ macrophages | CD11b$^{+}$, CD163$^{+}$, CD206$^{+}$ |
| **Dendritic cells** | Conventional dendritic cells type 1 (cDC1) | CLEC9A$^{+}$, IRF8$^{+}$, HLA-DR$^{high}$ |
| | DC | CD11c$^{+}$, HLA-DR$^{high}$ |
| | Follicular dendritic cells (FDC) | CD11c$^{+}$, CD21$^{+}$ |
